# How the microbiome liberates iron from heme

**DOI:** 10.1101/2024.10.14.618232

**Authors:** Arnab Kumar Nath, Ronivaldo Rodrigues da Silva, Colin C. Gauvin, Emmanuel Akpoto, Victoria Adedoyin, Mensur Dlakić, C. Martin Lawrence, Jennifer L. DuBois

**Author notes:** These authors contributed equally (Arnab Kumar Nath, Ronivaldo Rodrigues da Silva, Colin C. Gauvin). Corresponding authors: Jennifer L. DuBois, C. Martin Lawrence **Email:**.

## Abstract

The microbiome is a hidden organ system with diverse metabolic roles. Using *Bacteroides thetaiotaomicron* (*B theta*) as a model, we asked how microbiome species process iron, the quintessential biometal supporting respiration and life as we know it, when iron is provided in complex with protoporphyrin IX (PPIX) as *heme*. An intestinal transporter for heme has not been unequivocally identified. Canonical pathways for reclaiming heme iron from host cells or the diet require O_2_, which is unavailable in the GI tract and many pathological microenvironments. HmuS, a homolog of cobalamine-(vitamin B12) and chlorophyll-building chelatases, is widespread in anaerobic microbial ecosystems. In this work, we provide direct physiological, biochemical, and structural evidence for the anaerobic removal of iron from heme by HmuS, a de-chelatase that deconstructs heme to PPIX and Fe(II). We show that heme can be used as a sole iron source by *B theta*, yielding PPIX, and that *hmuS* inactivation under these conditions is lethal. Iron removal from heme depends on the *B theta* membrane fraction, NADH, and O_2_ exclusion. Absorbance spectra indicate that heterologously expressed HmuS is isolated with a non-substrate heme bound and can accommodate multiple heme equivalents under saturating conditions. Solution of the cryoEM structure reveals heme and two cations at sites that are conserved across the HmuS family and the chelatase superfamily, respectively. The proposed structure-based mechanism for iron removal further links biosynthetic and biodegradative pathways for heme, chlorophyll, and vitamin B12, three ancient and biologically crucial metallocofactors.

**Significance Statement:** Iron in its heme and non-heme forms is essential for nearly all life. This work defines a microbial mechanism for metabolizing heme that is widespread in gastrointestinal and other anaerobic ecosystems. To date, an intestinal cellular transporter for heme has not been unequivocally identified. However, heme, abundant in both red meat diets and animal cells, is converted by the bacterial HmuS enzyme into a non-heme form from which the iron can be repurposed. HmuS enzymes form a distinct de-chelatase subset of a superfamily of chelatases that catalyze metal insertion into chlorophyll and vitamin B12, indicating convergence of metabolic pathways for biosynthesis and degradation of three of life’s essential metallocofactors. Understanding HmuS function supports engineered approaches to directing microbiome-host metabolism and offsetting heme-associated pathologies including anemia, inflammation, and cancer.

## Introduction

Iron is the flint that supplies the catalytic spark for almost all cellular life and a growth-limiting resource for humans and their resident microbiomes. Aerobic life, including pathogens in the highly aerated niches of the blood, lung, and urinary tract, depends on iron for trafficking, sensing, activating, detoxifying, and deriving respiratory energy from O_2_. By contrast, the parts of the gastrointestinal (GI) tract where dietary iron absorption occurs (duodenum and proximal jejunum) are anaerobic at their core and replete with anaerobes from the Bacteroidetes and Firmicutes phyla. These common microbiome species require iron for harvesting energy and building materials from the host’s diet. They are believed to have roles in facilitating host iron uptake and modulating the risk of diseases related to iron mismanagement, particularly anemia, inflammatory diseases, and cancers^1,2^. Pathological niches outside the GI tract, including tumors, anoxic wounds, atherosclerotic plaques, and the subgingival layer, harbor anaerobes from the same phyla. These strains steal iron from their hosts to support their survival^3^.

Most of the ∼2g of iron in the healthy human body is heme-associated^4-6^. 2.3 million new hemoglobin-packed red blood cells alone are produced each second. To meet such high metabolic demands, heme is rigorously recycled by macrophages, which retrieve damaged red blood cells, salvage the iron, and deliver it to the marrow for repackaging^7^. Humans also assimilate dietary iron by efficiently catabolizing heme, primarily from red meat^8^. Direct intake of dietary heme by nutrient-absorbing, intestinal epithelial cells has not been unequivocally demonstrated^9^. However, Bacteroidetes species and many Firmicutes that dominate the GI-tract are known to import heme, which they require but cannot make biosynthetically^10^. How they meet their iron and heme requirements, or whether they metabolize heme on behalf of the microbiome and host, is unclear.

We recently showed that *Bacteroides thetaiotaomicron* VPI 5482 (*B theta*), an obligate anaerobe, heme auxotroph, and model GI-tract species, can use heme as its sole source of iron^11,12^. Well-described heme disassembly mechanisms use O_2_ to cleave the PPIX macrocycle and free the iron^13-15^. An anaerobic, radical S-adenosyl methionine dependent, heme-cleaving pathway was recently identified in a handful of pathogens^16-19^. A more widespread role for this pathway among the commensal, non-pathogenic anaerobes of the GI tract has not been identified. *B theta* and related anaerobes, whether commensal or pathogenic, have instead been proposed to break down heme by separating iron from PPIX in a de-chelation reaction, leaving the porphyrin intact^20,21^. Biological de-chelation reactions, however, have never been clearly demonstrated, and their existence remains potently controversial^22,23^.

**Figure 1.**
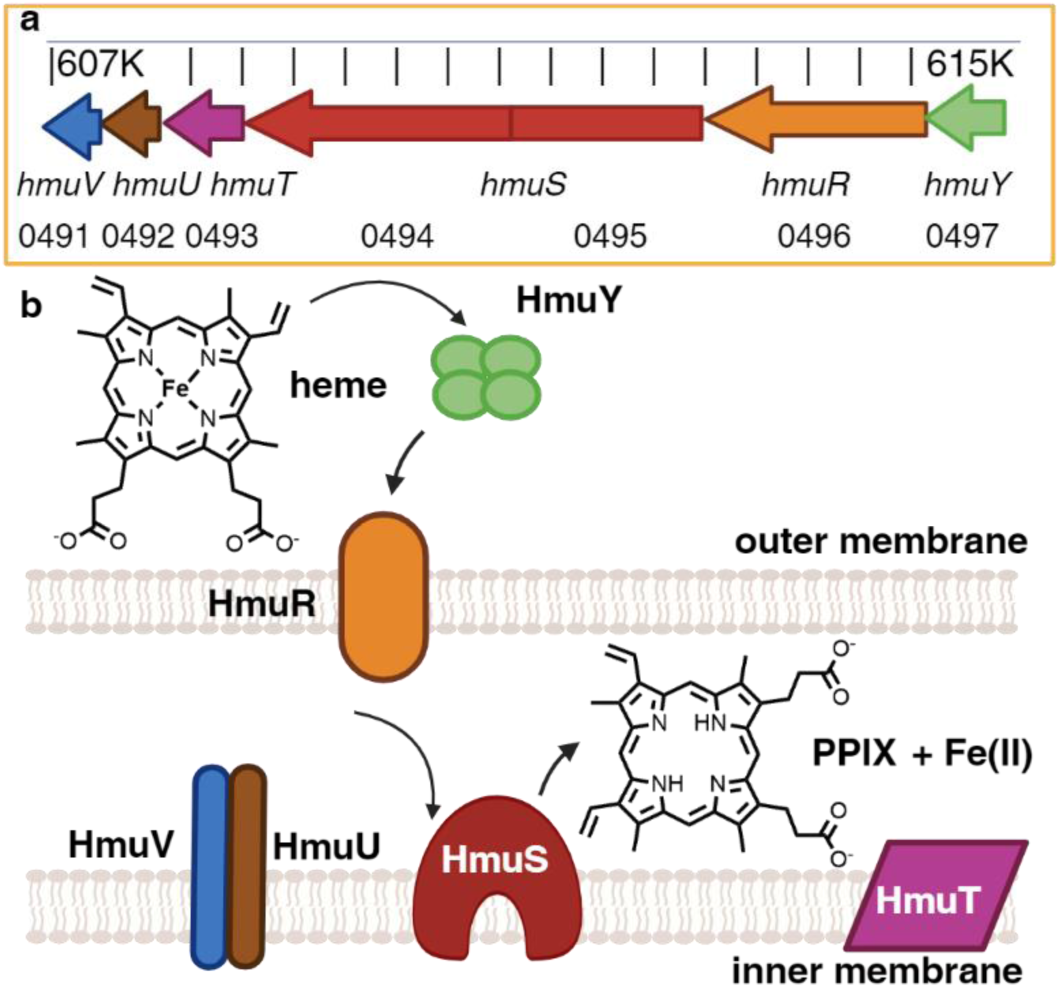
Organization of genes and proteins encoded by the heme utilization operon from *Bacteroides thetaiotaomicron* VPI-5482. (a) Magnified region of the complete genome between nucleotides 607K-616K (GenBank accession AE015928.1) is shown. Arrows indicate each of the *hmuYRSTUV* genes encoded on the antisense strand. BT loci numbers are given beneath the gene names. The *hmuS* gene was erroneously frame shifted in the initial genome assembly, though the error was corrected subsequently (GenBank assembly GCA_900624795.1). (b) Predicted organization of gene products. HmuY, a hemophore, is a monomeric lipoprotein that forms a tetramer when bound to heme^56^. HmuR is a TonB-dependent outer membrane heme transporter that, in conjunction with HmuV and HmuU, uses the inner membrane electrochemical gradient to power heme import to the periplasm. HmuS removes iron from heme, generating PPIX. HmuT is a predicted inner membrane protein that likely forms part of a larger complex.

Here, we provide a comprehensive *in vitro* and *in cellulo* description of the enzyme encoded by *hmuS*, part of a 6-gene operon (*hmuYRSTUV*) pervasive in members of phylum Bacteroidetes and the metagenomes of healthy human GI tracts (Fig. 1), definitively identifying it as a functional metal de-chelatase. Phenotypic analyses of *B theta* and its transposon insertion mutants demonstrate that HmuS is required for using heme as an iron source, with PPIX as the side-product^24^. Functional studies of the heterologously expressed HmuS protein demonstrate that it converts ferrous heme to PPIX and Fe(II)_aq_ in an NADH-dependent fashion in the absence of O_2_. As a critical step toward a mechanistic description of the de-chelation reaction, we report the first 3-dimensional structure for a protein from this family in its heme- and cation-bound form, obtained at 2.6 Å resolution by cryoelectron microscopy (cryoEM). We enumerate key differences between HmuS and homologous (class 1) chelatases responsible for inserting divalent manganese and cobalt, respectively, into chlorophyll (ChlH) and vitamin B12 (CobN) in the aerobic pathway^25-30^. This widespread bacterial heme lyase may promote the assimilation of dietary heme iron by the host and act as a critical regulator of microbiome and human health.

## Results

### Phenotypic analyses of wild type (*wt*) and *hmuS* mutant strains show HmuS is required for heme conversion to PPIX and Fe(II)

Cultures of *B theta* were grown on a minimal medium including residual non-heme iron (≤1 μM) and added hemin (15 μM ferric protoporphyrin IX chloride, referred to here as *heme*) as iron sources. Cultures were monitored optically over time until saturation (Fig. 2a), then pelleted and resuspended/lysed in a solvent that co-extracts heme and PPIX with equal efficiency. Extracts were analyzed for heme and PPIX by high performance liquid chromatography (HPLC)^31^ (Supplementary Fig. 1-2). These cells accumulated substantial heme (40 ± 0.9 μM) relative to a much smaller quantity of PPIX (0.035 ± 0.006 μM) (Fig. 2a, Supplementary Fig. 3).

The experiment was repeated but with 0.3 mM bathophenanthroline disulfonic acid (BPS), a non-metabolizable Fe(II) chelator that sequesters non-heme iron, added to the growth medium. These cells experienced a longer lag phase before entering exponential growth and saturated at lower optical density (Fig. 2a). Moreover, while the heme concentration in the cell pellets was similar to the unstressed condition, PPIX increased by more than an order of magnitude (Fig. 2a, Supplementary Fig. 3). This indicates that *B theta* can both incorporate heme into the cell mass and catabolize it to meet its non-heme iron needs, leaving PPIX as a by-product.

**Figure 2.**
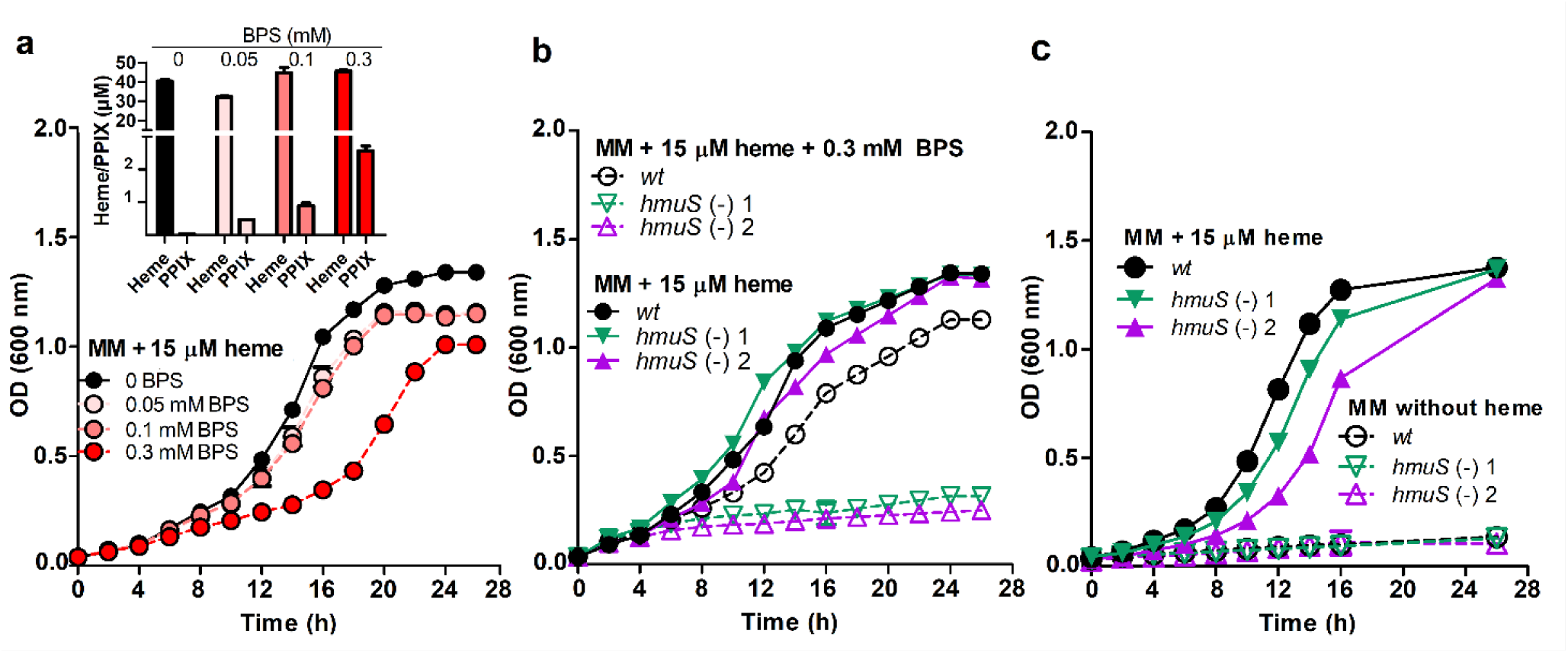
*Bacteroides thetaiotaomicron* requires *hmuS* to grow with heme as its sole iron source. (a) Growth curves illustrate response of *B theta* wild-type (*wt*) strain in 10 mL minimal medium + 15 µM hemin chloride (inoculated to optical density (OD) at 600nm = 0.03) to different concentrations of Fe(II)-sequestering BPS. Inset: heme/PPIX were extracted and quantified from cell pellets collected at the end of the growth. ACN: 12M HCl:DMSO (41:9:50, v/v/v) was used for the extraction of heme and PPIX (0.12 g wet cell pellet mL^-1^ of extraction solution). Heme and PPIX were separated and quantified relative to standards using HPLC (Supplementary Fig. 3). (b) Wild type and *hmuS* mutant strains were grown in minimal medium + 15 µM hemin chloride in the presence/absence of 0.3 mM BPS to eliminate available non-heme iron. Under the +BPS condition, *wt B theta* can grow with heme as the sole iron source, but the mutants cannot. (c) Serial passaging of *B theta* cultures into heme-limited growth medium was carried out to eliminate heme carryover in the inoculum. Four passages (18 h growth for each passage) into minimal medium without heme were carried out prior to inoculating cells into 15 µM heme-containing or heme-free minimal medium containing sufficient non-heme iron. No strains grew without heme. For all experiments, the optical monitoring of cell growth was carried out in crimp-topped Balch tubes under an atmosphere composed of 2.5% H_2_/97.5% N_2_, incubated at 37 °C, 150 rpm and monitored every 2 hours for 26 hours. See Table S1 and Supplementary Fig. 3.

We next grew two strains of *B theta* in which *hmuS* had been inactivated by insertion of a transposon (mutants 1 and 2), each at a unique site in the *hmuS* gene (Supplementary Fig. 4)^32^. Both grew indistinguishably from *wt* in the standard medium (Fig. 2b), accumulating similar amounts of heme and a trace amount of PPIX (not shown). This result suggested that both mutants could meet their cytoplasmic heme and non-heme iron requirements. However, neither strain grew in the medium amended with BPS. This result indicates that *B theta* requires a functional HmuS protein to obtain non-heme iron from heme. As a control, *wt* and mutant *B theta* strains were also grown with sufficient non-heme iron but without added heme. Surprisingly, all appeared to grow under these conditions (not shown), suggesting that carryover of heme stored by the inoculum could sustain the cultures. However, when cells were passaged four times into heme-free medium to eliminate carryover, the mutant strains were viable in a heme-/non-heme-iron-sufficient medium, but failed to grow without heme (Fig. 2c).

### Conversion of heme to PPIX by *B theta* depends on the membrane fraction and NADH

We next examined the small-molecule dependence of heme catabolism using lysed *B theta* cells (0.067 g cell pellet or 1.7 x 10^9^ *B theta* cfu resuspended per mL of 20 mM Tris-HCl, pH 7.1). The lysates (300 μL) were incubated anaerobically with 100 μM hemin for 30 or 40 min, after which an equal volume of extraction solvent was added to capture PPIX and unreacted heme for HPLC analysis (Supplementary Fig. 5). For lysed but undialyzed cells, ∼85% of the porphyrin content was unaltered heme and 15% was PPIX, with no other porphyrins detected by UV/visible (UV/vis) absorbance or fluorescence spectroscopy (Fig. 3a, Supplementary Fig. 5a and Supplementary Table 1). The ratio of PPIX:heme decreased only slightly following extensive dialysis to remove loosely affiliated small molecules. This suggests that the hydrophobic PPIX detected in either case was released only upon extraction.

**Figure 3.**
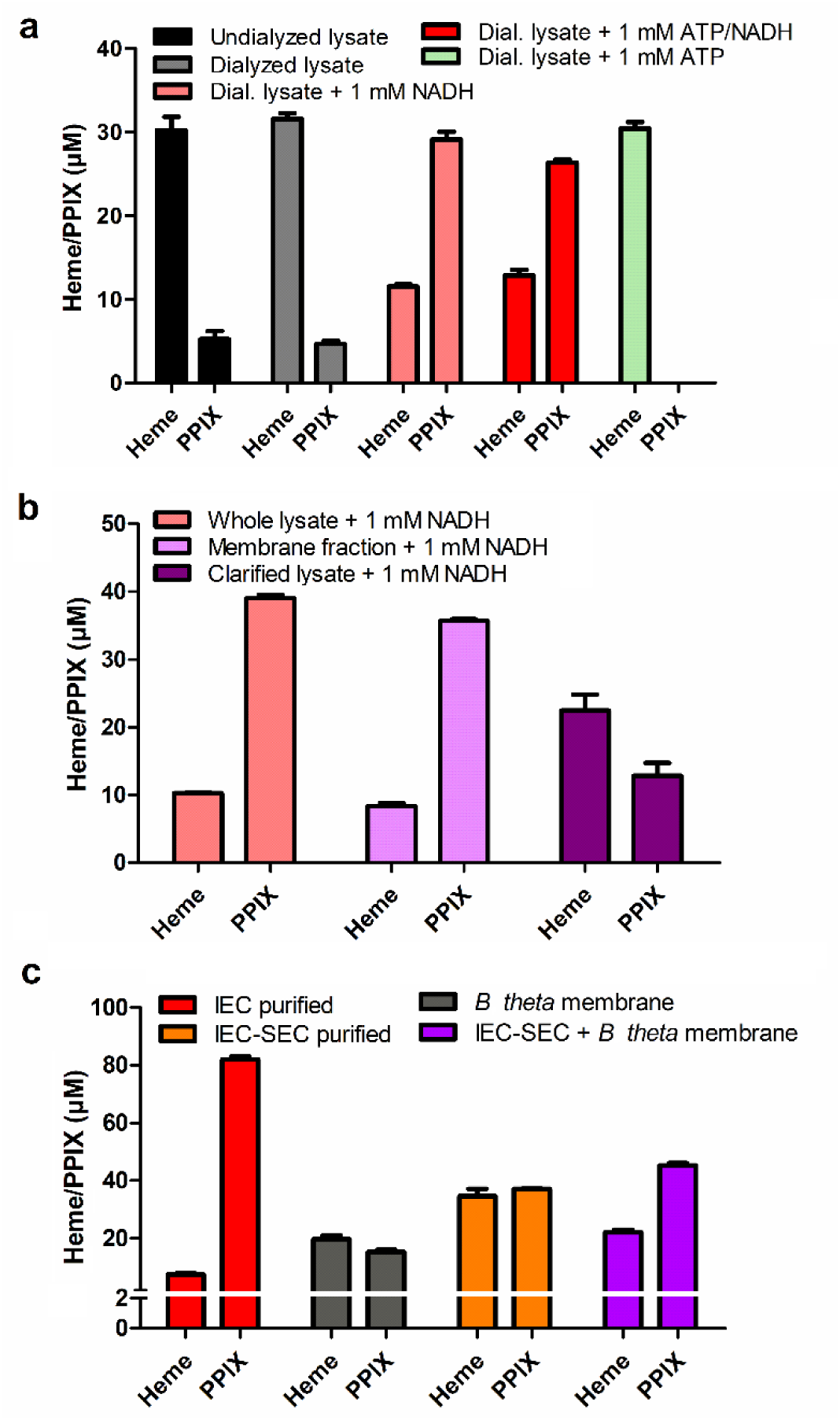
PPIX production by *B theta* cell fractions or purified recombinant HmuS incubated with heme illustrates the reaction dependence on NADH. The role of small molecules is monitored in a), and the subcellular fraction associated with enzymatic activity is identified in b). (a) The effects of 1 mM NADH and/or 1 mM ATP on 100 μM heme conversion to PPIX using 300 μL undialyzed/dialyzed *B theta* cell lysate (1 g cell pellet resuspended in 15 mL of 20 mM Tris-HCl buffer, pH 7.1) are shown. (b) *B theta* whole cell lysate (post-dialysis), soluble fraction (clarified lysate), and membrane fractions were reacted with 1 mM NADH and 100 μM heme. All reaction mixtures were prepared in 20 mM Tris buffer, pH 7.1 and incubated anaerobically at 25 °C for 30 min (a) and 40 min (b), followed by extraction into 330 μL ACN:12M HCl:DMSO (41:9:50, v/v/v)) and HPLC quantification of heme and PPIX. Concentrations of analytes are referenced to the final reaction volumes (330 μL). Triplicate results were averaged. Error bars represent ±1 standard deviation. See Table S1 and Supplementary Fig. 5. (c) Recombinant HmuS converts hemin to Fe(II) and PPIX in the presence of NADH and a probable mediator protein. Reaction mixtures contained 100 μM hemin, 1 mM NADH, in 330 μL total reaction volumes (25 mM Tris-HCl, 250 mM NaCl, pH 7.1) and the following protein fractions: partially (IEC) purified HmuS (3.2 mg mL^-1^ protein; 2.3 mM HmuS-heme) (red); IEC and SEC purified HmuS (3.2 mg mL^-1^ protein; 1.9 mM HmuS-heme) (orange); *B. theta* membrane fraction (150 μL from a stock prepared from 0.1 g cells mL^-1^) (gray); the same mixture as in gray but with added IEC-SEC-purified HmuS (final concentration in reaction: 1.6 mg mL^-1^; 0.95 μM HmuS-heme). Reactions were incubated for 40 min at room temperature prior to extraction. Averages of 3 measurements are shown as bar charts. (See Supplementary Fig. 12 and Table S1). Error bars represent ±1 standard deviation.

Reactions were carried out under identical conditions but with 1 mM NADH added, yielding 71% PPIX and only 29% residual heme. 1 mM ATP had no added effect when co-presented with 1 mM NADH. However, in the presence of 1 mM ATP alone, no PPIX was observed. The reasons for the apparent ATP-dependent inhibition of PPIX production are unclear; however, unlike the reactions catalyzed by type 1 chelatases, the heme de-chelation does not require ATP^33,34^ (Fig. 3a).

To determine whether catalysis depended solely on the membrane fraction (to which HmuS is anchored, Fig. 1), the dialyzed cell lysate was fractionated by ultracentrifugation. The supernatant (soluble component) and washed/resuspended pellet (membrane component) were each anaerobically incubated with 100 μM hemin and 1 mM NADH for 40 min before quantifying PPIX and heme by HPLC. The membrane fraction yielded approximately 80% PPIX and 20% heme after extraction, while the soluble fraction gave 36% PPIX and 64% heme (Fig. 3b, Supplementary Fig. 5b and Table 1). These results are consistent with an NADH-dependent, heme-converting activity localized in the membrane fraction, where we hypothesize that the PPIX in the soluble fraction could be due to HmuS solubilized by sonication.

### Heterologous expression yields HmuS in complex with ferrous heme

To examine the de-chelation reaction directly, soluble HmuS was expressed in *E. coli* BL21(DE3)-Lemo without its predicted N- and C-terminal membrane-anchoring helices (Supplementary Fig. 6-7). The HmuS-containing cells produced a pink lysate enriched in a protein with the expected HmuS molecular weight. The centrifuge-clarified lysate had a UV/visible absorbance spectrum typical of a 6-coordinate, low-spin ferrous hemoprotein with a prominent Soret band (λ_max_ = 425 nm), a weakly absorbing, sharp α band at 560 nm, and a less intense 530 nm β band^35^ (Fig. 4a). The spectrum was remarkably stable, remaining unchanged for >24 hours in air at room temperature (Supplementary Fig. 8).

**Figure 4.**
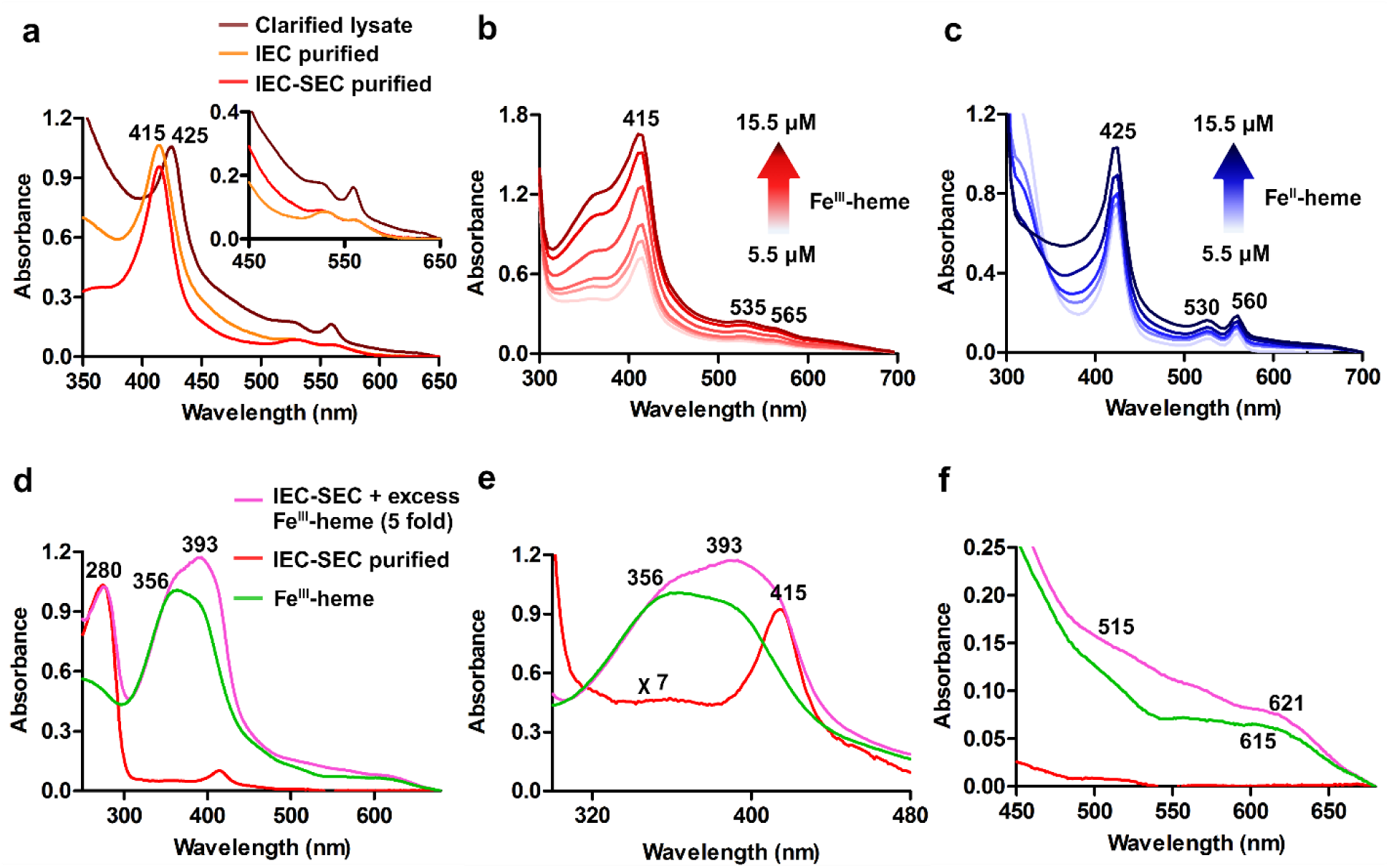
UV/vis absorbance spectra of recombinant HmuS report on heme binding. (a) Rose-colored *E. coli* clarified lysates exhibited a reduced heme-protein spectrum following HmuS expression. Following purification by anion exchange (IEC) and then size exclusion (IEC-SEC) chromatographies, the protein solution became orange and then red, respectively. The absorbance spectra indicated an oxidized, 6-coordinate heme protein (all spectra represent *ca.* 10 mM heme-protein content measured by pyridine hemochromagen assay). (b) Ferric heme was added to partially heme-loaded, IEC-SEC-purified HmuS (9.5 mg mL^-1^ protein; 5.5 mM HmuS-heme complex). Final heme concentrations: 7.0, 8.5, 10, 11.5, 15.5 mM. The increasing intensity of the spectral features, without changes in their energies, suggested increasing occupancy of a single heme-binding site without conversion to PPIX. (c) The same concentration of purified HmuS with 5.5 mM heme bound was reduced by addition of dithionite. Dithionite-reduced heme was added in the same increments as in panel b. (d-f) HmuS (1 mg mL^-1^ protein, 0.8 mM HmuS-heme, red spectrum) was incubated with excess heme (30 mM) followed by 3 cycles of concentration/resuspension in fresh buffer to remove loosely bound heme, and the spectrum of the resulting protein (27 mM HmuS-heme) measured (purple). The spectrum for a 22 mM ferric heme stock solution is shown for comparison. Spectra are shown (d) from 250-680 nm to visualize the protein peak at 280 nm, (e) focused on the Soret region (red spectrum shown on a magnified scale), and (f) in the Q-band region. Peak maxima are identified. All spectra were measured in 20 mM TrisHCl, 250 mM NaCl, pH 7.1. Heme/protein concentrations were determined using the pyridine hemochromagen and Bradford assays.

### The HmuS-bound heme oxidizes during purification

HmuS was purified from the clarified lysate in air using anion exchange chromatography (IEC). The eluted fractions featured a contaminant band near 60 kDa on SDS polyacrylamide gels (SDS-PAGE) that correlated with the density of the HmuS band at 158 kDa (Supplementary Fig. 9). Mass spectrometric (MS) analysis confirmed that the excised band derived mainly (73%) from HmuS peptides, while the 158 kDa was 90% HmuS (Supplementary File 01). Protease inhibitors, metal chelators, centrifuge filtration (100 kDa MWCO), and affinity purification (using HmuS with an N-terminal His6-tag) had little effect on these peptide contaminants (not shown).

IEC fractions enriched in full-length HmuS on SDS-PAGE were pooled, yielding an orange-hued solution containing >30% HmuS-derived peptides according to proteomic analysis of the trypsin-digested fraction (Supplementary Fig. 10, Supplementary File 02). Further purification by size exclusion chromatography (SEC) largely removed contaminants below 60 kDa, including a yellow protein that eluted in the 20-30 kDa range (Supplementary Fig. 8g). SDS-PAGE and proteomics analyses indicated that the pooled, red, IEC-SEC purified protein was predominantly full-length HmuS. (Supplementary Fig. 8a. See Supplementary Table 2 for a typical purification and Supplementary File 02 for proteomics analyses of IEC- and IEC-SEC purification fractions.) The UV/vis spectra for column-purified HmuS (Fig. 4a, Supplementary Fig. 8b) indicated a ferric, low-spin, 6-coordinate heme, with peak maxima (nm): 415 (Soret), 365 (shoulder), 535 (β), and 565 (α)^36,37^.

### Heme is associated with distinct sites on HmuS

We examined HmuS/heme association via several approaches. First, we attempted to remove the ferric heme bound to the purified protein by dialysis against imidazole as a competitive heme ligand. However, dialysis had no effect on the protein’s UV/vis spectrum or the amount of HmuS-heme measured by the pyridine hemochromagen assay, suggesting that the heme was tightly bound.

Second, we added ferric hemin in 1.5 μM increments to a concentrated solution of purified HmuS, monitoring changes in the UV/vis spectrum following each substoichiometric addition. Pyridine hemochromagen analysis indicated 5.5 μM heme bound to HmuS at the start of the experiment (apparent HmuS-heme *ε*_(415nm)_ ≈ 110 mM^-1^ cm^-1^). We observed increases in the Soret/shoulder and α/β band intensities after each addition up to a final HmuS-heme = 15.5 μM, with no additional UV/vis bands (Fig. 4b).

Third, we observed that excess NADH (1 mM) added to either the IEC or IEC-SEC purified HmuS-ferriheme complex (5.5 μM HmuS-heme), converted the oxidized heme into a reduced, low spin 6-coordinate heme (Supplementary Fig. 11) with a UV/vis spectrum closely resembling that of the clarified *E. coli* lysate^36,37^ (Fig. 4a). The spectrum did not change over 1h, indicating that reduction did not lead to conversion of the bound heme to PPIX and Fe(II). To monitor ferrous heme binding to HmuS, a 5.5 μM HmuS-heme sample was reduced with a 3-fold excess of dithionite. The resulting UV/visible spectrum of the HmuS-heme complex had Soret (425 nm) and visible bands (530, 560 nm) consistent with the UV/visible features observed in the lysate (Fig. 4a). Reconstitution of the complex with dithionite-reduced heme (to 15.5 μM HmuS-heme) gradually increased each absorbance band with no appearance of additional bands (Fig. 4c). Together, these observations suggested that adding small increments of heme to HmuS, in the ferric or ferrous state, did not result in the occupation of multiple heme binding sites or conversion of the heme to PPIX.

Finally, we incubated purified, partially heme-bound HmuS (1 mg mL^-1^ protein or ≤6.3 μM HmuS including 0.8 μM HmuS-ferriheme complex) with excess hemin (30 μM) to saturate the heme binding sites. Weakly bound heme was removed by 3 cycles of buffer exchange. Comparing the spectra before and after incubation (Fig. 4d) showed an identical band at 280 nm, reflecting the unchanged protein concentrations of the two samples. However, the pyridine hemochromagen assay indicated 27 μM HmuS-heme complex, or at least 4 equivalents of heme bound per HmuS molecule. The UV/visible absorbance spectrum (Fig. 4e-f) revealed a broadened Soret band with apparent peaks at 356, 393, and 415 nm, and a broad Q-band region with maxima at 515, 560, and 621 nm. The similarity of the λ_max_ values to those for both the HmuS-ferriheme complex and ferriheme in aqueous solution suggests that weakly HmuS-associated heme is present in the heme-saturated sample. However, the added peak maxima, particularly at 393 nm in the Soret region, pointed toward occupancy of an additional, stably-ligated heme site.

### Expressed HmuS converts ferroheme to PPIX

Conversion of heme to PPIX and iron was monitored under conditions similar to those used for *B theta* cellular fractions but using HmuS expressed in *E. coli* (Fig. 3c, Supplementary Fig. 12). Reactions were incubated anaerobically for 40 min with 100 μM heme and 1 mM NADH before extraction of PPIX and heme for analysis by HPLC. When IEC-purified HmuS was the catalyst, 92% of the added heme was converted to PPIX. An equal mass of IEC-SEC purified enzyme also converted heme to PPIX, though with fewer turnovers (54%), despite containing a greater proportion of HmuS. The purified enzyme furthermore augmented the heme-converting activity of *B theta* membranes when the two fractions were combined, relative to the activity of the membrane fraction alone (Fig. 3c Supplementary Table 1).

We hypothesized that SEC partly removed an *E. coli* protein supporting HmuS activity, potentially a reductase capable of mediating electron transfer from NADH. Consistent with that hypothesis, SEC partially separated a yellow, 20-30 kDa protein from HmuS. The proteins eluting in this mass range had absorbance features characteristic of both heme and a yellow flavin cofactor (Supplementary Fig. 8). Proteomic analyses of the IEC- and IEC-SEC-purified HmuS and these small-molecular-weight SEC fractions identified a single 26 kDa *E. coli* NAD(P)H flavin reductase common to all 3 samples (Fre, UniprotID P0AEN1), that IEC partly removed^38^ (Supplementary File 02). *E. coli* Fre generates reduced flavins that interact with multiple redox partners, including ferrisiderophores and heme proteins^39^. Fre or a functionally similar protein could enable electron transfer from NADH to a flavin cofactor and ultimately to HmuS-heme. An analogous, membrane-associating reductase may be present in the *B theta* membrane fraction. Consistent with this hypothesis, a flavodoxin was among the most upregulated proteins in a recent survey of *B theta* transcriptional responses to a low iron challenge^40^.

### Structure of HmuS

The structure of HmuS was determined by cryo-EM single particle analysis at a global resolution of 2.6 Å. DALI and PDBeFold searches each identified the greatest similarity to cobaltochelatase from *M. tuberculosis* (Mt CobN, PDB ID 7c6o), followed closely by Mg chelatase from *Synechocystis* (Syn ChlH, PDB IDs 4zhj and 6ysg)^26,28,29^. HmuS de-chelatase shares the overall 6-domain architecture of these class I chelatases, including the N-terminal domains termed “head” (domain I) and “neck” (domain II), followed by domains III, **-**VI, with domain IV inserted within domain III, which collectively form the body. However, structural superposition on CobN or ChlH showed a substantially different conformation of the head domain in HmuS relative to those in both CobN and ChlH (Fig. 5; see below).

**Figure 5.**
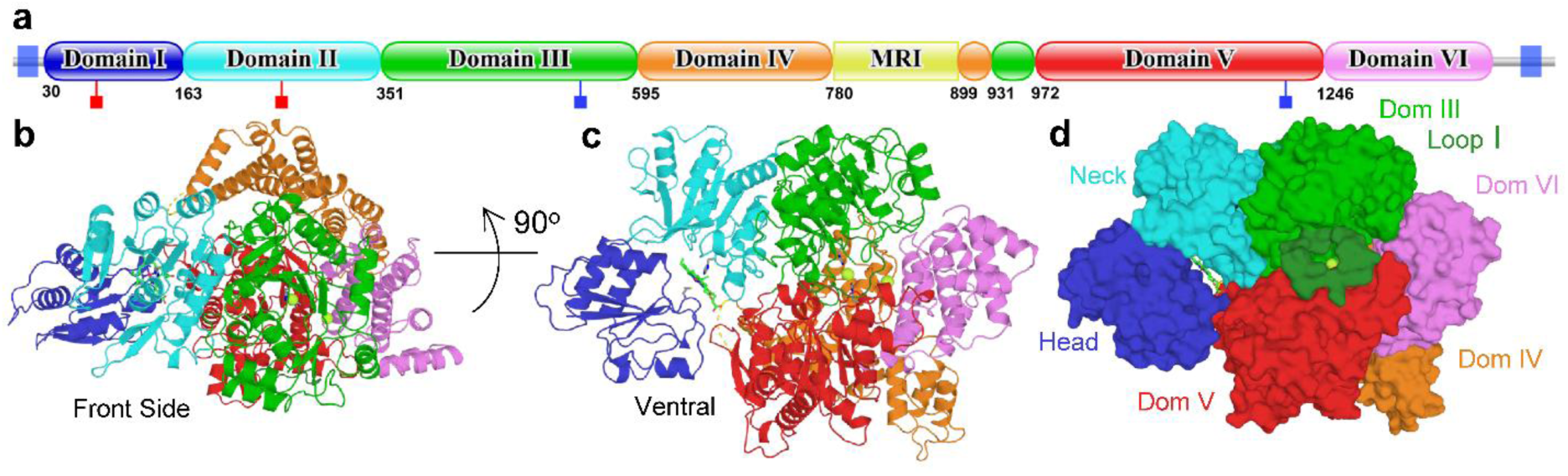
Structure of HmuS. (a) Domain organization of HmuS. Blue squares at each end indicate transmembrane regions. Domains I-VI are equivalent to known class 2 chelatase domains. The first residue number for each domain is shown. The cryo-EM map lacks density for the 120 residues Methionine Rich Insertion (MRI). While the MRI is particularly enriched in Met residues, Lys and Ala are also overrepresented. Red sticks indicate residues interacting with heme at the head/neck interface (Met-79 and His-254, from left to right), while blue sticks indicate ion-coordinating residues (His-538 and His-1209, from left to right). (b) Cartoon diagram of Hmus shown from the “front side”. Individual domains are colored as in panel A, the MRI is not present, but is an insertion in Domain IV (orange), predicted to lie above domains I (dark blue) and II (cyan), which form the N-terminal “head” and “neck” domains of HmuS. The body is formed by domains III-VI, which houses a central cavity formed by domains III, V, and VI, with domain IV running along the top (“backbone”). (c) HmuS is rotated 90 degrees about the horizontal axis, tipping the molecule to expose the underside (ventral view). The heme (green sticks) at the interface of the head and neck domain becomes apparent. (d) Surface representation of panel c (ventral view). Domain III is green. Loop I, thought to gate entry into the cleft is in the darker forest green. The *bis*-His coordinated sodium ion (Na1) is shown in “limon” and is visible through the Loop I pore. See also Supplementary Fig. 20 for views from the reverse side.

#### Methionine Rich Insertion

A structure-based multiple sequence alignment identified a 120 residue, low complexity, methionine rich insertion (MRI, residues 780-898, 21 Met) within domain IV (Fig. 5, Supplementary Fig. 13-14). Consistent with this low complexity, density for the MRI is absent, suggesting these residues are intrinsically disordered in the absence of an appropriate ligand. A conservative set of *bona fide* HmuS sequences was identified by focusing on complete, 6-gene *hmu* operons, and by selecting only HmuS sequences with at least 50% identity to *B theta* HmuS. This resulted in 680 non-redundant HmuS sequences that were multiply aligned to identify a set of strictly conserved residues across the HmuS family (Supplementary Fig. 13-14). The alignment confirmed the presence of an MRI in each HmuS sequence, though varying in length: 119 residues in *B theta,* a mean length of 82 residues, and a minimal length of just 22 (Supplementary Fig. 15). The MRI is clearly present in each HmuS sequence and appears to be a feature that distinguishes the HmuS de-chelatase family from other members of the class I chelatase superfamily. In addition to Met, Lys and Ala are also over-represented compared to their natural frequencies, with almost no other hydrophobic residues of the aromatic or aliphatic type.

#### HmuS body

Like ChlH and CobN, domains III through VI form a globular structure, known as the “body”, that houses a central cavity. In the chelatase enzymes, this cavity represents the enzyme active site, where the macrocycles are bound and metalated. The N-terminal half of the body is formed by domains III and IV, in which domain III adopts an α/β fold with a central, predominantly parallel, 6-stranded β sheet (β_3↑_β_4↑_β_1↑_β_5↑_β_6↑_β_7↑_), and domain IV is a predominantly helical, extended domain (Fig. 5, Fig. 6). Similarly, the C-terminal half of the body is formed by domains V and VI, with V present as an α/β domain with a central, predominantly parallel, 5-stranded β-sheet (β_4↑_β_1↑_β_5↑_β_6↑_β_7↓_), and an extended helical domain VI, containing two successive armadillo-like repeats (α1-α3, α4-α6) followed by a 3-helix bundle (α7-α9). The central cavity is at the nexus of domains III, V and VI. When the structure is aligned such that the predicted N-terminal membrane anchor points downward, holding the head domain close to the inner membrane, domain IV runs like a backbone across the top of the structure, with major contacts to domains III and VI.

**Figure 6.**
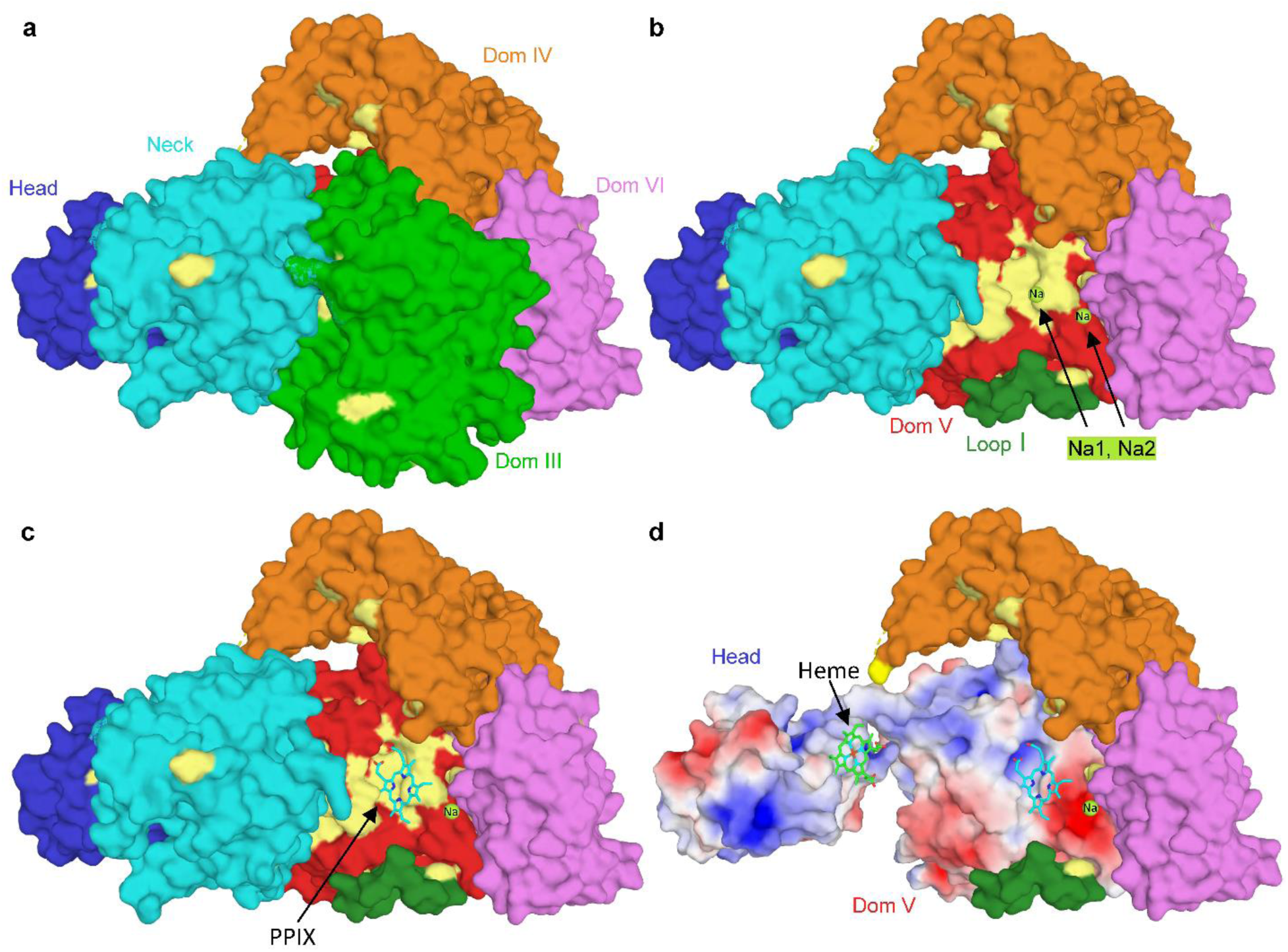
Ligand binding sites. (a) HmuS is shown in the same “side view” orientation as Fig. 5b, colored by domain. Strictly conserved side chains are in pale yellow; in this view strict surface conservation is limited. (b) Domain III has been removed, revealing the central cleft with 2 bound Na ions (limon), Na1 and Na2. Loop I, thought to gate access to the central cleft is, also visible in forest green. In contrast to the exterior surface, domain V present a large, strictly conserved surface to the central cleft (pale yellow on red). Na1 is coordinated by His-538, which lies in the center of this conserved face. (c) The central cleft is the chelatase active site in ChlH and CobN, suggesting it represents the de-chelatase active site in HmuS. We thus docked the HmuS product, protoporphyrin IX (PPIX), into the central cleft with AutoDock Vina. The docking model suggests HmuS also utilizes the strictly conserved surface on domain V for recognition of PPIX. (d) Domain II (cyan) has also been removed to reveal the heme binding site at the Head/Neck interface. The heme and PPIX molecules are separated by nearly 30 Å. Electrostatic surfaces are also shown for domains I and V (+/- 5kT/e). The propionate groups for both ligands are accommodated by significant positive charge, and in PPIX they are accommodated by a conserved basic pocket in domain V.

Mapping the HmuS multiple sequence alignment (above) on the structure shows the central cavity of HmuS is highly conserved (Fig. 6). Notably, an HFG(T/A)HG motif (residues 534-539) in the β6/α8 loop of domain III and an LSLDHxxEFMGG motif in the β5/α14 loop and α14 of domain V are strictly conserved. His-538 in the first motif and His-1209 in the second extend toward each other, directly across the middle of the putative active site, where they coordinate strong spherical density, which we have modeled as a sodium cation due to the inclusion of NaCl in the buffer (Fig. 6-7, Supplementary Fig. 18). Strictly conserved Glu-1212 forms a hydrogen bond with the His-1209 side chain, likely fixing the position and fine-tuning the acidity of the imidazole. The observed cation coordination by His-538/His-1209, in conjunction with docking studies, suggests HmuS could utilize these residues for *bis*-His heme coordination within the central cavity. The axial histidines are also well placed to act as general acids, protonating the porphyrin as iron is excised. While strictly conserved in our limited set of HmuS sequences, His-538 also appears strictly conserved across the chelatase superfamily. In addition to these motifs, Ser-1014, Ser-1067, Gln-1198, Ser-1199 and Trp1202 are among the strictly conserved residues lining the surface of the HmuS central cleft, and along with the unsaturated N-terminus of domain V helix α3, are predicted to coordinate the propionate groups of a bound heme/PPIX. Consistent with this, while sequence identity to CobN and ChlH are low, structural superposition on CobN or ChlH shows greatest similarity to domains III and V (Supplementary File 03).

**Figure 7.**
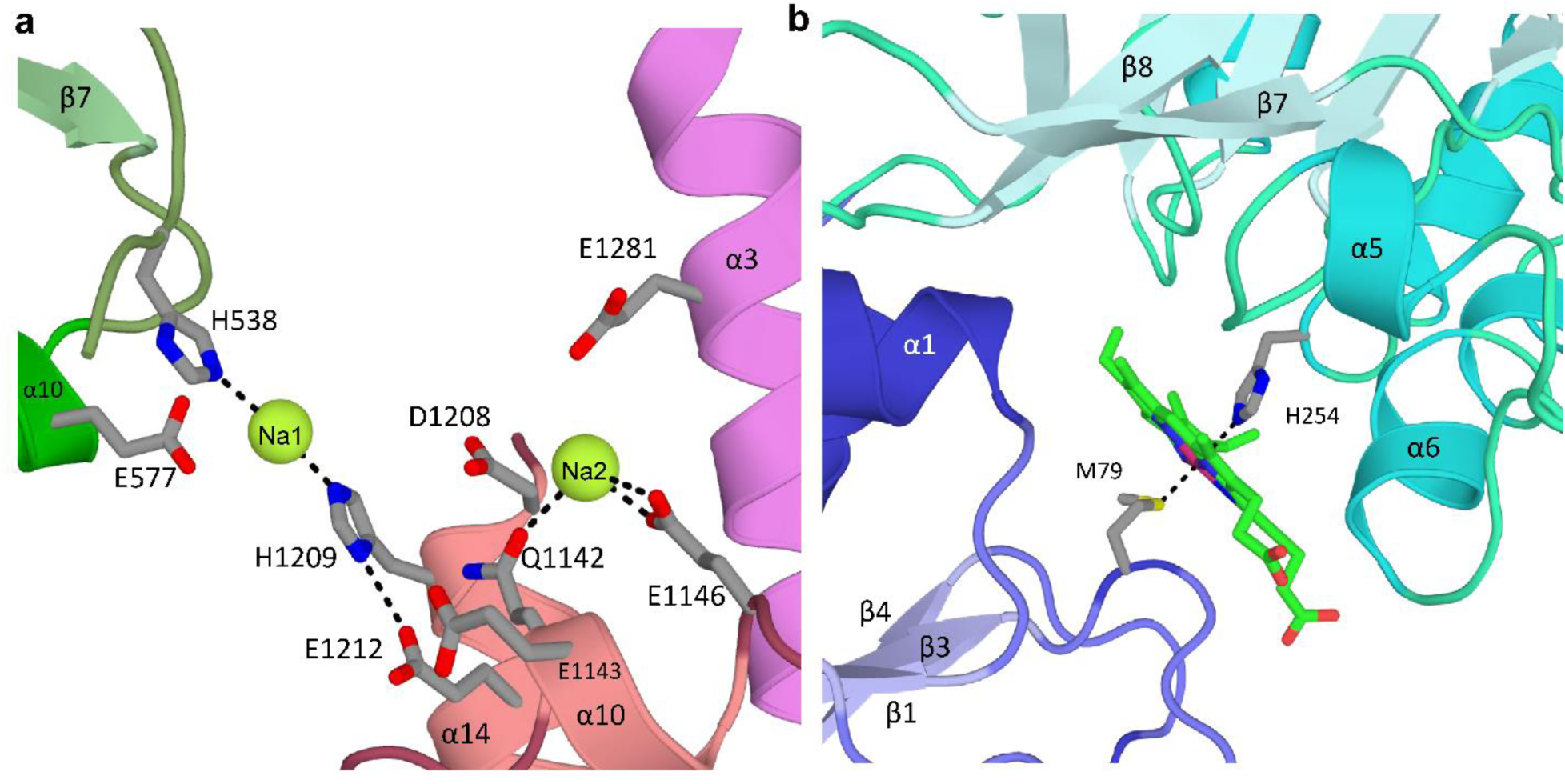
Ligand Coordination. (a) HmuS is shown from the ventral side (Fig. 5d). Elements of domains III, V and VI involved in Na^+^ ion coordination within the central cleft are shown in green, salmon and magenta respectively. Na1 is coordinated by His-538 and His-1209, with Glu-1212 coordinatingHis-1209. Na2 is coordinated by Gln-1142 and Glu-1146. Residues coordinating Na2 are not strictly conserved. Additional residues contributing to a patch of strong negative charge within the central cleft are also shown (Glu-1143, Asp1208 and Glu-1281). (b) Coordination of heme at the interface of the Head and Neck domains by Met-79 and His-254. The Head domain is in shades of blue, and Neck domain in shades of cyan. In both panels, secondary structure elements are labeled relative to their specific domain.

#### A non-Pro cis peptide

The potential importance of His-538 may also be indicated by *a cis* peptide bond between Thr-537 and His-538. Interestingly, a *cis* peptide bond is also present in CobN between an equivalent His residue (His-522) and the preceding Lys residue^26^. In contrast, a roughly equivalent His residue in ChlH is in the *trans* configuration, with the side chain more distal to the porphyrin metal binding site. Whether these residues undergo *cis/trans* isomerization during the catalytic cycle, as they insert or remove metal ions, remains to be determined.

#### Active site accessibility

In ChlH, access to the central cavity is blocked by Loop I, which is in a closed conformation^25 28^. In contrast, the equivalent loop in CobN was found in an open, partially disordered state^26^. Like CobN, the central cavity is also solvent accessible in HmuS, as the β5/α7 loop in domain III is locked in an ordered, “partially open” conformation. However, when modeled in the closed conformation by analogy to ChlH, strictly conserved Phe-502 and His-506 are predicted to lie over the solvent exposed (propionate distal) edge of the heme, proximal to His-538 and His-1209 in a heme/protoporphyrin free structure.

*Structural similarity with class II chelatases*. The head (domain I) and neck (domain II) domains show distant structural similarity to those in CobN and ChlH. Domain I exhibits a 4-stranded parallel β-sheet (β_2↑_β_1↑_β_3↑_β_4↑_) with right-handed helical crossovers and two additional C-terminal helices, while domain 2 builds on this core β-sheet structure with two additional strands at the N-terminus to give a 6-stranded β-sheet, and a C-terminal β-finger (Fig. 5, Fig. 7).

However, unlike CobN and ChlH, the central β-sheets in each of these domains “points” toward the other, with a distinct cleft between the two domains. We, and others^26,28^, also note distant similarity between the head and neck domains of the class I chelatase structures with human (class II) ferrochelatase, and to bacterial periplasmic binding proteins in general.

**Figure 8.**
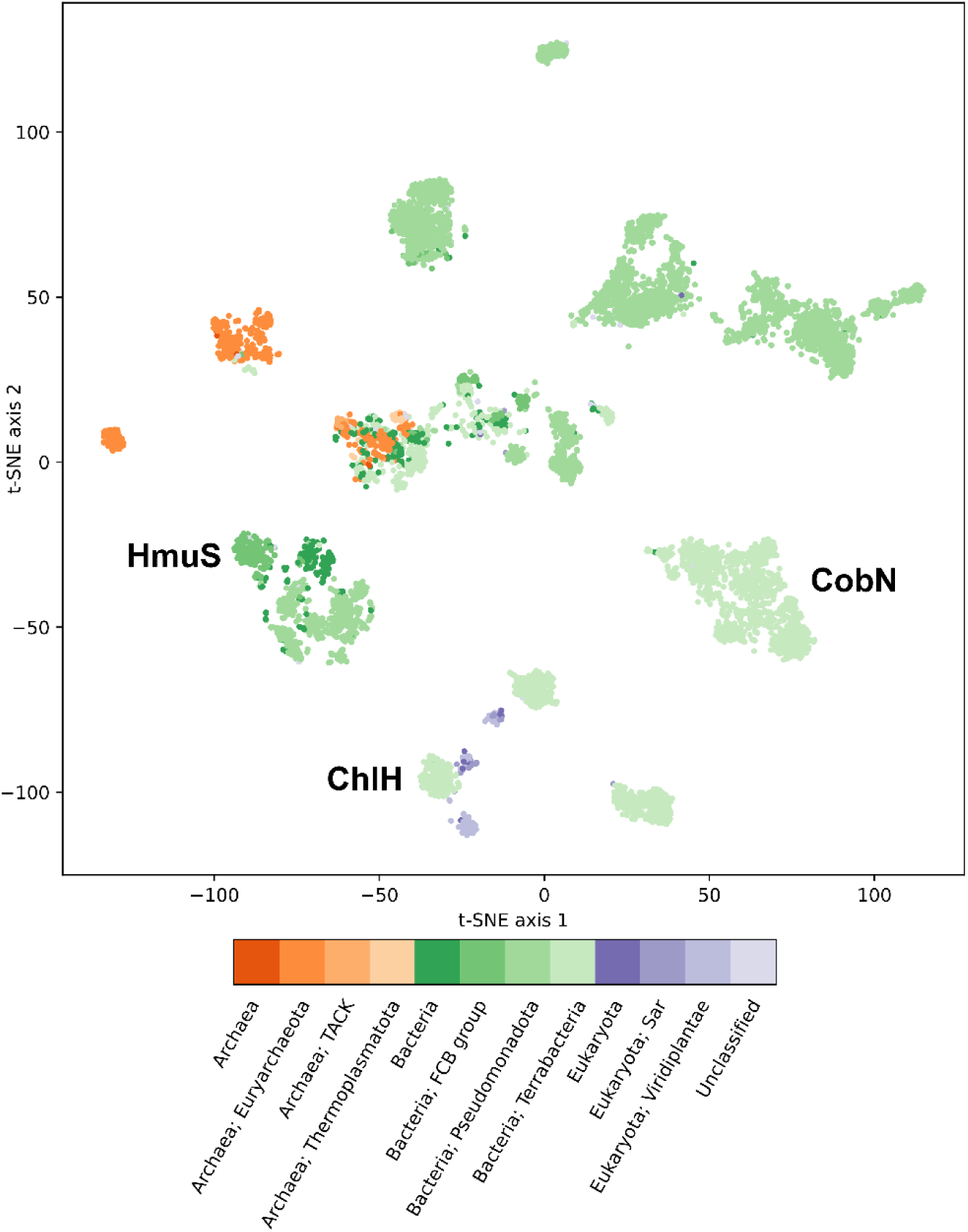
2D embedding of chelatase superfamily sequences. High-dimensional numerical representations of protein sequences were reduced to 2 dimensions by t-SNE^51^. Each data point was colored by its taxonomic domain membership: Archaea (orange), Bacteria (green), and Eukarya (purple). Phyla with largest memberships within these domains are colored separately in shades of a similar color. Representative clusters for each of the three major families are labeled. Family assignments (HmuS, ChlH, and CobN) are made on the basis of biochemically characterized anchoring sequences, of which there are very few. Only the clusters nearest the labels on the diagram can be ascribed with a known function. Most of the remaining bacterial and archaeal sequences cannot be reliably annotated without further investigation.

#### Heme binding site at the head/neck interface

The HmuS multiple sequence alignment identifies only a few strictly conserved, solvent-exposed residues in the head and neck domains. Notably, these include Met-79 and Gly-80 in the head, as well as a His-Gly-Arg motif (residues 254-256) and Gln-299 in the neck (Fig. 7, Supplementary Fig. 18). Each of these residues lies at the interface between the head and neck domains. Critically, we also identified a bound heme at this interface that copurified with HmuS, with the iron coordinated by the strictly conserved Met-79 and His-254 residues (Fig. 6-7). Given the structural similarity of the head and neck domain to class II chelatases, there has been speculation that class I chelatases might utilize this site to coordinate ligands. However, this is the first time a tetrapyrrole has been found at this site. It should be noted that Met-79 and His-254, while conserved among HmuS sequences, are not universally conserved in the chelatase superfamily, raising the possibility that HmuS binds heme at the head/neck interface in a way that is distinct from other superfamily members. Further, superposition of the HmuS head domain on CobN or ChlH requires rotations of 50° and 100°, respectively, away from the neck domain (Supplementary Fig. 19). Our HmuS structure is thus in a closed conformation, which in the absence of heme, might sample open conformations, perhaps similar to that seen for CobN.

## Discussion

Well-described cellular mechanisms for releasing iron from heme use O_2_ to produce a linearized porphyrin, most commonly bilirubin plus carbon monoxide^15,41^. The portion of the GI-tract in which the microbiome breaks down complex nutrients, however, is anaerobic at its core, as are several pathological microenvironments within tumors. Anaerobes from the Bacteroidetes phylum, which require but cannot make heme, numerically dominate in these O_2_-limited niches when heme is attainable. The heme-microbiome interaction has been directly implicated in the induction of tissue damage and disease in these pathogenic niches as well as the GI-tract, where the microbiome constitutes a missing link between red meat diets, cellular damage, and cellular hyperproliferation that precedes colon cancer^42^. How diverse GI-tract organisms promote or potentially mitigate tissue damage is not fully understood. It is unknown whether these organisms metabolize dietary heme on behalf of the host, perhaps contributing to the enhanced bioavailability of heme iron in the human diet. Importantly, the molecular mechanism by which they break down heme, without the benefit of O_2_, has remained the subject of particularly spirited debate^20-22,43^. Here, we used multiple experimental approaches to demonstrate that an anaerobic process for <u>h</u>e<u>m</u>e <u>u</u>tilization, encoded by the *hmuS* gene from the widely distributed *hmu* operon, removes iron from heme, leaving PPIX intact^11^.

*B theta* is a model GI-tract species bearing just one copy of the intact *hmu* operon, including a single copy of the catalytic class I chelatase encoded by *hmuS*. The lone *hmuS* gene greatly simplifies genetic approaches for describing HmuS function^21^. When both heme and non-heme forms of iron were supplied in a chemically defined growth medium, we observed that *wt* and *hmuS* mutant strains of *B theta* grew indistinguishably. However, when iron was provided only in its Fe(III)-heme form, *wt B theta* experienced a growth lag and accumulated PPIX, while the *hmuS* mutants failed to grow. We localized the *in cellulo* de-chelatase activity to the membrane component, consistent with the sequence-based prediction that HmuS is integrally attached to the inner membrane. However, unlike its CobN or ChlH homologs, which catalyze ATP-driven metal chelation reactions, the HmuS-containing membranes required added NADH but did not expend ATP. The *hmu* operon likewise does not encode ATP-dependent accessory proteins homologous to ChlD/ChlI^44-46^ or CobS/CobT, which are necessary for the function of their accompanying chelatases^27^.

To examine the de-chelation reaction under well-defined conditions, we expressed the ∼1400 amino acid HmuS monomer without its membrane-anchoring helices. Surprisingly, the protein emerged with sub-stoichiometric heme bound in a low spin hexacoordinate and reduced/ferrous state that oxidized upon column purification. The same site could be augmented with added heme and reduced in an apparent flavin-dependent reaction by NADH. These observations led us to conclude that the as-isolated HmuS heme may be bound in a non-catalytic site.

Solution of the cryoEM structure of HmuS revealed a monomeric 6-domain protein similar to available ligand-free structures of the CobN and ChlH chelatases that have been zoomorphically described as “chicken-like”^26-28,30,46^. However, unlike prior structures of homologs, HmuS featured a heme molecule tucked between the craning head and neck (domains I-II). The heme iron was axially ligated by Met-79 and His-254, residues conserved specifically in HmuS and not CobN/ChlH sequences. This is the non-substrate heme observed in the UV/visible absorbance spectrum of as-isolated HmuS. However, 34 Å away and inside body’s cavernous interior, two cations (assigned as Na^+^(1) and Na^+^(2)) were identified amid a zone of highly conserved residues. This region, featuring a strikingly well-conserved hydrophobic surface that readily accommodates PPIX, marks a likely location for substrate conversion to products, which then travel outward from the protein (Fig. 6).

ATP-independent ferrochelatases catalyze the same reaction as HmuS, though biased in the reverse direction (Fe(II) + PPIX^2-^ ↔ [Fe(II)PPIX]^2-^ + 2H^+^)^43,47^. The ferrochelatase paradigm and the experimental observations presented here suggest a model for HmuS function. The substrate heme could easily fit in the vicinity of Na^+^(1), which has conserved residues suggestive of heme ligation, including proximal His-1209/Asp-1208 and distal His-538/Glu-577 histidine-carboxylate dyads that could act as axial ligands or acid-bases, Ser/Gln residues and main chain amines at the N-terminus of helix V-α3 that could interact with propionates, and a surface of conserved side chains lining the pocket. The enzyme’s dependence on NADH, the lack of detectable porphyrin products other than PPIX, and the spectroscopically observed reduction of the HmuS-bound heme (Fig. 3, Supplementary Figs. 11-12) together suggest that NADH reduces the heme iron^11^. A flavin-dependent *E. coli* NADH oxidoreductase is clearly present in the IEC-purified HmuS and depleted by SEC, along with the enzymatic activity of HmuS (Fig. 4). *B theta* has at least one flavodoxin that is highly upregulated under iron-deficient conditions^40^. The flexible hinge connecting the head domain to the neck and the proximity of the MRI suggests that ferric heme can access a binding site at the head/neck interface, possibly fed by the MRI. This could be the site of heme-iron reduction or receipt of a reduced heme^16^ which might be transferred by HmuS into the Na^+^(1) site. Alternatively, heme bound at the HmuS head/neck interface could receive electrons from NADH via a reductase, transferring one to a ferric heme at the Na^+^(1) site and reducing it *in situ*. However it arrives, we expect the substrate heme to be in the ferrous state. Deformation of the planar PPIX ligand would expose the pyrrole nitrogens^48^ to proton-donating side chains, from alternate faces of the porphyrin plane or from the same face with both pyrrole nitrogens aspiring toward the same active site acid. Protonation should restore planarity to the deformed PPIX as Fe(II) is liberated. A conserved pathway for metal egress, leading to the inner membrane Fe(II) transporter (FeoB), could begin with a way station near the less-conserved Na^+^(2) site^49^. A defined pathway for PPIX release is also likely.

Confirming the function of HmuS as a *de-chelatase* motivates redrawing the boundaries of the family within the chelatase superfamily. Alignment of HmuS sequences derived from complete *hmu* operons supplied a core set of family-defining sequence features, including the MRI, the Met-79 and His-254, and the series of characteristic motifs described above. Aligning the much larger superfamily of sequences, including the HmuS set, identified a far shorter list of residues conserved superfamily-wide, placing HmuS distinctions in even sharper relief (Supplementary file 03). To perform a deeper analysis of the chelatase superfamily, we combined search results for representative members of the three major families. After converting these ∼10,000 sequences into complex numerical representations using protein language models (ProtT5), a 2D representation of the whole dataset was created by t-distributed stochastic neighbor embedding (t-SNE)^50,51^.Subdivision of the entire superfamily is not done just according to sequence similarity, because it also includes a complex encoding of physical and chemical properties of individual sequences. HmuS sequences form an island amid an archipelago containing more than the minimally anticipated 3 familial clusters (Fig. 8). Biochemically, genetically, and/or structurally characterized representatives of these groups functionally anchor some of the clusters. The existence of further subdivisions beneath these known functional umbrellas, in some cases, appears to follow taxonomic lines; for example, green plants and phototrophic cyanobacteria possess Mg^2+^ chelatases whose divergence may reflect the distance between the taxa. Other sequences, annotated as CobN subunits, come predominantly from Methanobacteriota and cluster according to taxonomic class into sibling groups from *Halobacteria* and thermophilic or methanogenic eubacteria, though their functions are unclear. It is possible that members of some of these sequence clusters share (de)chelatase functions with HmuS/CobN/ChlH, but preferentially act on modified porphyrin or corrin rings. Some may lack (de)chelatase functions entirely.

HmuS-utilizing Bacteroidetes sp. are often dominant members of the human microbiome. Whether heme is derived from the host diet or the recycled lining of the intestine, HmuS-mediated heme de-chelation presents a means of recovering the iron from heme. The lengths to which cells go to protect themselves from Fe(II)-driven redox stress, which can lead to DNA, lipid, and protein damage, especially near aerobic/anaerobic interfaces, are well known. PPIX can also be highly toxic. When exposed to light, it fluoresces, offering a means for endoscopically identifying porphyrin-overproducing infectious agents like *Bacteroides fragilis* or *Helicobacter pylori* inside surgical wounds or in the stomach^52,53^. Photoexcited PPIX catalyzes the conversion of triplet O_2_ to the highly reactive singlet state, which is leveraged in photodynamic therapies to destroy cancerous or other diseased tissue, and which has pathological consequences, for example, for patients with porphyrias who accumulate PPIX in the skin. Porphyrins likewise hyperaccumulate in diseased or intoxicated liver and in tumor microenvironments and associated serum, where they π-stack, intercalate into membranes, or precipitate^54^. Even in dark or anaerobic spaces, the redox activity and water-insolubility of PPIX and heme make each a troublesome guest. Because heme, PPIX, and iron are not appreciably excreted by healthy individuals, we expect that an optimal anaerobic microbiome manages these critical molecules, ideally supporting their nutritional and metabolic benefits while mitigating their potential for harm.

## Materials and Methods

Summary methods are given here, with further details in Supplementary Information.

### Strains and genes

*Bacteroides thetaiotaomicron* V-5482, ATCC 29148 contains one complete *hmuYRSTUV* operon (loci (BT0497-BT0491)) (Scheme 1). The original genome annotation erroneously assigned *hmuS* to two loci (BT0495, BT0494) though they were subsequently identified as a single gene in all Bacteroidetes strains (1463 aa, RefSeq: WP_008765019.1)^11^. A transposon insertion library for *B. theta* V-5482 generated by Shiver *et al.^24^* was the source of two hmuS-deficient strains, each containing a single transposon insertion into the *hmuS* coding region (Supplementary Fig. 3). The *wt* strain was acquired from the ATCC.

### Monitoring *B theta* growth

*B theta* was cultivated in a chemically defined minimal medium (MM). Balch-type, crimp-topped tubes containing 10 mL MM, 15 µM hemin, ± 0.3 mM BPS were kept in the anaerobic glove box for 4 hours before use (2.5% H_2_/97.5% N_2_ atmosphere). *B theta* frozen glycerol stocks were used to prepare a liquid pre-inoculum in MM + 15 µM hemin (37 °C, 150 rpm, 14 h). The pre-inoculum was centrifuge-pelleted, washed twice in hemin-free MM, and added to fresh media to an optical density (OD_600nm_) of 0.03. Bacterial growth was monitored every 2h and metabolites analyzed from pelleted cells at saturation. 70 mL cultures were grown similarly in crimp-topped bottles.

### Extracting and quantifying heme and PPIX from *B theta*

An extraction solvent containing acetonitrile, 12M HCl, and DMSO in a 41:9:50 volume ratio (final [HCl] = 1.08 M) was used to recover heme and PPIX from cells or reaction mixtures^31^. 10 mL cultures were centrifuge-pelleted, washed, resuspended (0.12 g mL^-1^) in extraction solvent, transferred into FastPrep Lysis B-matrix tubes, and lysed. Following centrifugation, the well-resolved upper layer was removed for quantifying PPIX and heme by high-performance liquid chromatography (HPLC).

### *B theta* fractionation

Cell pellets (1g) were resuspended (20 mM Tris-HCl buffer, pH 7.1) and lysed by sonication. Whole lysates were dialyzed in air to remove small molecules. Soluble proteins were isolated by centrifugation. The pelleted membrane fractions were washed and resuspended in the same buffer.

### Heterologous expression, purification, and UV/visible absorbance characterization of HmuS

The soluble HmuS protein from *B theta* (accession WP_022471467.1) was heterologously expressed in *E. coli* BL21(DE3)-Lemo cells (NEB) without the predicted single transmembrane spanning helices at its N- and C-termini (Supplementary Fig. 6). Cultures were pelleted by centrifugation and stored at –80 °C, then thawed/resuspended in 20 mM Tris pH 7.1. Suspensions were lysed by sonication in the presence of multiple protease inhibitors and ultracentrifuged to obtain a clear supernatant (clarified lysate). The clarified lysate was purified by anion exchange chromatography using a linear elution gradient of 0-500 mM NaCl. Fractions enriched ≥50% in a protein of the expected molecular weight (158 kDa) were pooled (ion exchange fraction). This fraction was centrifuge concentrated (100 kDa MWCO) to ≤1 mL and further purified by preparative size exclusion chromatography. Proteins were eluted in 20 mM Tris pH 7.1, 250 mM NaCl. HmuS-enriched fractions were identified by SDS -PAGE, concentrated, flash frozen in liquid N_2_, and stored at –20 °C, ≥10 mg mL^-1^ until further use. Protein and heme concentrations were routinely determined by Bradford and pyridine hemochromagen assays^53^.

### Sequence conservation and clustering analyses

Using cblaster (https://github.com/gamcil/cblaster) against the Identical Protein Groups database (https://www.ncbi.nlm.nih.gov/ipg/) all HmuS proteins that are found in the same operon with the other 5 Hmu proteins were considered to fulfill the operonic criterion^11^. We selected only complete HmuS sequences with at least 95% coverage and at least 50% identity relative to *B theta* HmuS.

Sequences of *B. theta* HmuS, *M. tuberculosis* CobN (PDB code 7C6O) and *Synechocystis spp* magnesium chelatase ChlH (PDB code 6YT0) were used as queries for three iterations of PSI-BLAST searching. All hits were pooled together, and all redundant sequences were removed. ProtT5 protein language model was used to create an average 1024-value vector for each protein sequence^50^. All numerical vectors were subjected to dimensionality reduction by t-SNE to create a 2D grouping of sequences^55^.

### Structural analysis

Tools used for single particle transmission cryoelectron microscopy (CryoTEM) data collection, analysis, and model building are described in Supporting Information. The model was deposited in the Protein Data Bank with accession code PDB ID 9D26 and the map was deposited in the Electron Microscopy Data Bank with accession code EMD-46483.

## Author Contributions

A.K.N. carried out experiments with expressed HmuS. R.R.d.S. performed experiments with *B theta* strains and developed/implemented methods for co-extracting and quantifying heme and PPIX. C.C.G. measured the cryo-EM data and solved the structure. E.A. helped express, purify, and quantify HmuS, and conducted bioinformatics studies. V.A. assisted R.R.D.S. with experiments. M.D. made the AlphaFold2 model of HmuS and performed bioinformatics analyses. C.M.L. helped solve the cryo-EM structure, coordinated the study, and co-wrote the manuscript draft. J.L.D. obtained funding, designed and coordinated the study, analyzed the data, and wrote the manuscript draft. All authors prepared the Figures and tables, provided critical revisions, contributed to manuscript editing, and approved the final manuscript. A.K.N., R.R.d.S., and C.C.G. contributed equally.

## Supporting information

Supplemental Information HmuS

## Acknowledgments

AKN, RRS, EA, VA, and JLD thank the National Institutes of Health grant R35 GM136390 (US Health and Human Services Department) for funding. Funding for the Montana State University Cryo-EM Facility was contributed by National Science Foundation (DBI-1828765), the MJ Murdock Charitable Trust, The National Institute of General Medical Sciences (P30GM140963) and the MSU Office of Research, Economic Development and Graduate Education. Structure calculations and other analyses were performed on the Tempest High Performance Computing System, operated and supported by University Information Technology Research Cyberinfrastructure at Montana State University. We thank Dr. Jessica Lusty Beech for technical support and Garrett Moraski for helpful discussions.

