## Supplemental Information HmuS for "How the microbiome liberates iron from heme"

### Supporting Information for
How the microbiome liberates iron from heme

Arnab Kumar Nath[1], Ronivaldo Rodrigues da Silva[1], Colin C. Gauvin[1], Emmanuel Akpoto[1], Victoria Adedoyin[1], Mensur Dlakić[2], C. Martin Lawrence[1*], Jennifer L. DuBois[1*]

Address correspondence to Jennifer L. DuBois


**This PDF file includes:**

Supporting experimental methods text
Figures S1 to S20
Tables S1 to S3
Legends for Datasets S1 to S03
SI References

**Other supporting materials for this manuscript include the following:**

Datasets (Supplementary Files) 01 to 03

#### Supporting Information Text

##### Detailed Methods

**Biochemical stock solutions and equipment.** Chemicals were purchased through Fisher unless otherwise stated. Hemin (ferric heme chloride, Calbiochem®) stocks were prepared in 1M NaOH (pH 8) or DMSO and concentrations measured via UV/vis absorbance ($\varepsilon_{385}$ = 58.44 $mM^{-1}$ $cm^{-1}$). 200 µM PPIX stocks were prepared in acidified acetonitrile (ACN): ACN:1.7 M HCl (82:18, v/v). Both stocks were prepared in air and handled in amber bottles before storing at -20 °C. Reductant stocks (NADH, dithionite) were prepared in an anaerobic chamber (Coy Labs, 2.5% $H_2$ and 97.5% $N_2$) at 100 or 50 µM concentrations, respectively. Hemin and reductant stocks were aliquoted into single-use tubes, frozen, and discarded at the end of a day's use. Working solutions were similarly discarded within a day. An extraction solvent consisting of acetonitrile, 12M HCl, and DMSO in a 41:9:50 volume ratio (final [HCl] = 1.08 M) was used to equivalently recover heme and PPIX from cells and reactions[1]. Standards and reductants were diluted immediately prior to use. 20 mM TrisHCl buffer, pH 7.1 was used to resuspend *B theta* cell pellets prior to lysis and subsequent analyses. The same buffer was used as a storage and reaction medium for purified, heterologously expressed HmuS, though with added 250 mM NaCl.

UV/visible absorbance (UV/vis) spectra were measured in quartz cuvettes using a Cary 50 or in test tubes using a ThermoScientific Genesis instrument. Cell pellets (>1g) were lysed via Branson model 102C (CE) sonicator. High pressure liquid chromatography (HPLC) was performed on a Shimadzu Prominence-i LC-2030C 3D Plus with UV/visible detection (190-800 nm) using a Hypersil GOLD™ column (Thermo Scientific™, 4.6 mm x 250 mm, 5 µm particle size). Protein chromatography was carried out with a Next Generation Chromatography System (BioRad).

**Strains***.* The sequenced genome for the strain of *B theta* used in this work (*Bacteroides thetaiotaomicron* V-5482, ATCC 29148, GenBank accession: AE015928.1) contains one complete *hmuYRSTUV* operon (loci (BT0497-BT0491)) with the *hmuS* gene on the complement strand at nucleotide positions 608,809 – 613,201 (Scheme 1). The original genome annotation erroneously assigned this gene to two loci (BT0495, BT0494) though they were subsequently identified as a single gene in all Bacteroidetes strains (1463 aa, RefSeq: WP_008765019.1)[2]. A transposon insertion library for *B. theta* V-5482 was generated by Shiver *et al*[3,4]. Gene-inactivating, single transposon insertions into the *hmuS* coding region were identified in two of the library strains, mapping to nucleotide positions 610,214 and 612,897, respectively (Supplementary Fig. 3). These mutants were obtained from the Shiver group at Stanford University[3,4]. The *wt* strain was acquired from the ATCC.

**Monitoring *B theta* growth in chemically defined media.** *B theta* was cultivated in a pH buffered, chemically defined minimal medium (MM) composed of: 6.6 mM potassium dihydrogen phosphate, 15.4 mM NaCl, 98 µM $MgCl_2{\cdot}(H_2O)_6$, 176.5 µM $CaCl_2{\cdot}(H_2O)_2$, 4.2 µM $CoCl_2{\cdot}(H_2O)_6$, 50.5 µM Mn(II)$Cl_2{\cdot}(H_2O)_4$, 9.3 mM $NH_4Cl$, 1.75 mM $Na_2(SO_4)$, 134 µM L-methionine, 23.8 mM sodium bicarbonate, 8.25 mM L-cysteine (free base), 28 mM D-glucose, and 15 µM hemin chloride. The pH of the medium was adjusted to 7.1 using 5 M HCl. The MM supplemented with 15 µM hemin was transferred into Balch type, crimp topped tubes (10 mL MM) and kept in the anaerobic glove box for 4 hours before use (2.5% $H_2$/97.5% $N_2$ atmosphere).

*B theta* glycerol stocks were used to prepare the pre-inoculum in liquid MM + 15 µM hemin. After 14 h growth, the liquid pre-inoculum was centrifuged at 3,260 x g for 10 min at 4 °C. The cell pellet was washed twice and resuspended with MM without hemin, and used as an inoculum for monitoring bacterial growth and/or metabolite production. For optical monitoring of growth, MM (10 mL) in Balch type, crimp-topped tubes containing 15 µM hemin with or without added 0.3 mM BPS was inoculated with sufficient cell mass to produce an initial optical density ($OD_{600nm}$) of 0.03 in the Balch tube (2.5% $H_2$/97.5% $N_2$ atmosphere), incubated at 37 °C, 150 rpm, and monitored every 2 hours for 26 hours. For larger scale B theta fractionation experiments, 70 mL cultures were grown in crimp-top bottles containing anaerobic MM medium supplemented with 15 µM hemin, prepared in the anaerobic chamber as described above.

**Extracting heme and PPIX from *B theta* at the end of growth experiments.** 10 mL Balch tube cultures of B theta cells were pelleted by centrifugation, washed twice in ultrapure water, resuspended (0.12 g $mL^{-1}$) in extraction solvent, transferred into FastPrep Lysis B-matrix tubes, and lysed using a FastPrep 24 5g instrument (2 cycles: 6.0 meters/second for 40 sec). Samples were centrifuged (9,600 x g, 25 °C, 15 min), yielding a well-resolved upper layer that was removed for quantifying PPIX and heme by HPLC.

***B theta* cell-free extracts.** 1 g pellets (on the order of 2.6 x $10^{11}$ *B theta* cfu[11]) from 200 mL *B theta* cultures grown as described above were resuspended in 15 mL 20 mM Tris-HCl buffer (pH 7.1) and lysed by sonication on ice (Branson instrument: 7 min, with a pulse sequence of 10 s on and 25 s off, at 40% amplitude). Whole lysates were dialyzed in air against the same buffer at 4 °C using 12 kDa molecular weight cut off (MWCO) tubing. Soluble proteins were separated from the lysate by centrifugation (9,600 x g, 25 °C), and contained 1.65 mg $mL^{-1}$ protein (estimated by the Bradford assay, below).

**Heme conversion to PPIX.** 300 µL samples of the dialyzed lysate and soluble fractions were placed in Eppendorf tubes inside the glove box and amended to contain 1 mM NADH, 1 mM ATP, both, or neither. Reactions were initiated by adding 100 µM hemin (330 µL total reaction volume) and incubated in an anaerobic chamber at room temperature for 40 min prior to adding 300 µL extraction solvent, which denatured the proteins and solubilized both PPIX and unreacted hemin[1]. Proteins were precipitated by centrifugation (9,600 x g, 25 °C, 20 min) and the organic layer removed for analysis by HPLC.

**High-performance liquid chromatographic (HPLC) analysis of heme and protoporphyrin IX**. Chromatography was carried out using a linear gradient with solution A (ultrapure water + 0.1% trifluoroacetic acid, TFA) and solution B (ACN + 0.1% TFA), flow rate of 1 mL min$^{-1}$, oven temperature at 25 °C (Supplementary Fig. 1). 30 µL samples were injected onto the column. Standard curves of hemin chloride and PPIX (0.1, 0.5, 1, 5, 10, 50 mM) were prepared to quantify the extracted heme and PPIX via peak integration.

**Heterologous expression of HmuS.** The soluble HmuS protein from *B theta* (accession WP_022471467.1) was heterologously expressed in *E. coli* BL21(DE3)-Lemo cells (NEB) without the predicted single transmembrane spanning helices at its N- and C-termini (Supplementary Fig. 6). The gene was codon-optimized for *E. coli* and synthetically introduced between the NdeI/XhoI sites of pET28a(+) by GenScript (Supplementary Fig. 7).

A single colony from a freshly streaked Lysogeny Broth (LB)-agar plate was used to inoculate an overnight liquid LB culture on a rotary shaker (37 °C, 200 rpm). This subsequently seeded 6 x 1L flasks containing terrific broth. All media contained selective antibiotics (50 µg mL$^{-1}$ kanamycin, 34 µg mL$^{-1}$ chloramphenicol). Flasks were incubated on a rotary shaker to mid-logarithmic growth ($OD_{600}$ = 0.6-0.8). Isopropyl β-D-1-thiogalactopyranoside (IPTG, 1 mM) was added to induce protein expression at 16 °C, 200 rpm, for 16 h. The cultures were pelleted by centrifugation and stored at –80 °C. Cell pellets were resuspended in 100 mL of 20 mM Tris pH 7.1 and lysed by ultrasonication on ice with added protease inhibitors (0.5 mM phenylmethylsulfonate, 1.0 mM EDTA, and a protease inhibitor tablet (Protease Inhibitor Tablets, EDTA-free, Thermo Scientific). The lysate was ultracentrifuged (39,000 × g, 30 min, 4 °C) to obtain a clear pink supernatant (clarified lysate).

**HmuS purification**. Working in ambient air at 4 °C, the clarified lysate (100 mL) was loaded onto 80 mL HiPrep DEAE FF 16/10 DEAE sepharose fast flow anion exchange resin (Cytiva) equilibrated with 20 mM Tris pH 7.1. The column was washed with 800 mL 20 mM Tris-HCl and eluted using a linear gradient of the same buffer containing 0-500 mM NaCl (3 mL min$^{-1}$, 70 min). Fractions were analyzed by SDS-PAGE (12% acrylamide) and UV/visible absorbance spectroscopy. Fractions enriched ≥50% in a protein of the expected molecular weight (158 kDa) were pooled (ion exchange fraction). This fraction was centrifuge concentrated (100 kDa MWCO) to ≤1 mL and loaded onto a 140 mL size exclusion column (HiPrep 16/60 Sephacryl S-300 HR, Cytiva) pre-equilibrated with 20 mM Tris pH 7.1, 250 mM NaCl. Proteins were eluted isocratically in the same buffer at 0.1 mL min$^{-1}$. HmuS-enriched fractions were identified by SDS - PAGE, concentrated, flash frozen in liquid $N_2$, and stored at –20 °C, ≥10 mg mL$^{-1}$ until further use.

**Protein concentrations.** These were determined by Bradford assay. 0.1 mg mL$^{-1}$ bovine serum albumin (BSA from BioRad, stock concentration 2 mg mL$^{-1}$) was used to make 2.5, 5 and 10 mg mL$^{-1}$ standards. Absorbance at 595 nm was recorded (corresponding to the protein-dye complex). Standards and unknowns were recorded in triplicate.

**Heme concentrations by pyridine hemochromagen assay[5]**. The heme content of purified HmuS was measured by mixing 400 mL HmuS-heme complex (10 mg mL$^{-1}$) with 400 mL 40% pyridine in 0.2M NaOH(aq) along with 2 mL 0.1M potassium ferricyanide and measuring the absorption spectrum of the oxidized *bis*-pyridine bound heme. The oxidized heme-pyridine complex was then reduced by 20 mL 0.5M sodium dithionite in 0.5M NaOH (excess) and the characteristic absorption spectrum of the reduced heme-pyridine complex was obtained with a Soret band at 420 nm and β/α-bands at 525 and 557 nm. The concentration of the bis-pyridine bound reduced heme was determined using the extinction coefficient at 557 nm ($\varepsilon_{557}$ = 34.7 mM$^{-1}$cm$^{-1}$) corrected for dilution. The [heme] in HmuS-heme samples determined by this method was used in assigning an approximate e-value for the complex at its Soret peak maximum.

**Heme binding to HmuS.** *Dialysis to remove bound heme*. We attempted to remove heme from the as-isolated HmuS-heme complex by dialysis against imidazole-containing buffer. Before dialysis, the absorption spectrum of the heme-bound purified HmuS protein (4 mg mL$^{-1}$) was recorded to compare with the spectrum of the dialyzed protein. The protein (2.5 mL) was dialyzed in 12-14 KDa MWCO dialysis tubing against 20 mM Tris-HCl pH 7 containing 250 mM NaCl and 50 mM imidazole for 6 hours (two cycles each of 3 hours), followed by two more cycles of dialysis against buffer without imidazole. The dialyzed protein was then concentrated to the starting concentration of 4 mg mL$^{-1}$ using a 100 kDa MWCO centrifuge concentrator. The absorption spectra pre- and post-dialysis were unchanged, indicating that the heme remained attached to HmuS.

*Binding heme to HmuS*. To monitor ferric heme binding to purified HmuS, an aqueous solution of 150 mM hemin was (pH 8) added in 1 mM increments to 600 mL purified HmuS (9.5 mg mL$^{-1}$, 5.5 µM HmuS-heme complex) and monitored by UV/vis spectroscopy. To monitor ferrous heme binding to HmuS, the 150 mM hemin stock was purged with $N_2$ and reduced with a 3-fold excess of dithionite ($Na_2S_2O_4$, 450 mM) in an anaerobic chamber (Coy). The HmuS sample was degassed inside a cuvette and then reduced with a 3-fold excess of dithionite relative to the total HmuS-heme concentration. Stepwise reconstitution (1.5 mM increments) of ferrous heme to the HmuS protein was monitored by

UV/visible absorbance. Samples were allowed to come to equilibrium. The final spectra in the experiments were unchanged after 1h. To saturate one or more binding sites on the protein, HmuS (500 µL, 10 mg $mL^{-1}$) was incubated with excess hemin, aerobically and in the dark. Unbound or loosely bound heme was removed by 3 cycles of centrifuge concentration (50 kDa MWCO) and resuspension in fresh buffer (20 mM tris-HCl, 250 mM NaCl, pH 7.1), and the UV/visible absorbance spectrum measured. The concentrations of protein and heme in the sample were measured by Bradford assay and pyridine hemochromagen assay respectively.

**Monitoring heme turnover to PPIX by HmuS.** Experiments paralleled those carried out with *B theta* cell fractions. Reaction mixtures (3.2 mg $mL^{-1}$ protein, total volume 330 µL) were prepared in 20 mM Tris-HCl buffer, pH 7.1 with 1 mM NADH. 100 µM hemin was added to initiate the reactions, which were incubated anaerobically (2.5% $H_2$/97.5% $N_2$ atmosphere) at room temperature for 40 min. Increasingly pure fractions of recombinant HmuS were assayed: HmuS-containing clarified *E. coli* lysates, ion exchange (IEC) purified fractions, and IEC-SEC (size exclusion chromatography) purified HmuS. In addition, 150 µL *B theta* dialyzed lysate was assayed, with/without an equal volume of IEC-SEC purified HmuS. All reactions were stopped and extracted by adding 1 or 2 volumes of extraction solvent, the proteins were pelleted by centrifugation, and the extracts analyzed by HPLC (above).

**Determining conservation of primary sequences.** Using cblaster (https://github.com/gamcil/cblaster) against the Identical Protein Groups database (https://www.ncbi.nlm.nih.gov/ipg/ ), all HmuS proteins that are found in the same operon with the other 5 Hmu proteins were considered to fulfill the operonic criterion[2]. We selected only complete HmuS sequences with at least 95% coverage and at least 50% identity relative to *B theta* HmuS. This approach yielded 1605 sequences. Using CD-HIT (http://cd-hit.org), this group of proteins was further cleaned to remove redundancies. The resulting 680 non-redundant HmuS operonic fasta sequences were aligned using ClustalΩ. Sequence logo representation was created at https://weblogo.threeplusone.com/create.cgi (Supplementary Fig. 14). To determine sequence conservation within the larger superfamily of chelatase proteins, we started with HmuS as query and performed 5 search iterations with HHblits. Some of the identified ~1800 sequences were short, and we removed all sequences <1000 residues to generate a final alignment of 1513 chelatase sequences (Supplementary Fig. 17). See Supplementary files 03-05 for spreadsheets containing alignment scores and structural representations of conservation.

**Clustering analysis.** Sequences of *B. theta* HmuS, *M. tuberculosis* CobN (PDB code 7C6O) and *Synechocystis spp* magnesium chelatase ChlH (PDB code 6YT0) were used as queries for three iterations of PSI-BLAST searching. All hits were pooled together, and all redundant sequences were removed. After excluding sequences shorter than 1000 residues, 10201 chelatase superfamily members were used for subsequent analyses. ProtT5 protein language model was used to create an average 1024-value vector for each protein sequence[6]. All numerical vectors were subjected to dimensionality reduction by t-SNE to create a 2D grouping of sequences[7]. This type of non-linear embedding preserves the local relationships between vectors, but not necessarily their global distances. An interactive version of this embedding can be found as HTML in Supplementary File 05[8]. Hovering a mouse pointer over each dot will show its BLAST-determined sequence identity to *B theta* HmuS, followed by UniRef annotation of that sequence.

**Single particle analysis**.

*Cryo-TEM sample preparation.* Quantifoil 300 mesh copper R 1.2/1.3 holey carbon grids were plasma cleaned for 45 s at 15 mA with a PELCO easiGlow discharge cleaning system (Ted Pella) and placed into a Vitrobot Mark IV (Thermofisher) blotting apparatus at 4 °C and 95% humidity with Whatman grade 1 blotting paper. 4 µL of HmuS at 1.5 mg $mL^{-1}$ was applied to each grid, blotted for 4 s at blot force 4, and immediately vitrified by plunge-freezing into liquid ethane. Upon freezing, the samples were clipped into autogrids and stored in liquid nitrogen until analyzed.

*Cryo-TEM data collection.* Prepared grids were loaded into a Talos Arctica G2 transmission electron microscope (Thermofisher Scientific) operating at 200 kV and tuned for parallel illumination. Micrographs were collected using a Gatan K3 camera and a total electron exposure of 56 $e^-$ $Å^{-2}$ distributed over 51-frame dose-fractionated movies with a 3.06 s exposure time. SmartScope controlled SerialEM was used to collect 15,994 exposures at a nominal magnification of 45,000 (0.9061 Å $pixel^{-1}$) with target defocus values from -0.6 to -1.5 µm using a 5x5 multi-shot scheme, or beam-image-shift (BIS) distance of 7.5 µm, where appropriate[9,10]. The first 7,800 exposures were collected at $0^o$ tilt angle, the remainder at $15^o$.

*Data Processing.* Single particle reconstruction was performed with cryoSPARC[11,12]; the workflow is presented schematically in Supplementary Fig. 14. Each recorded movie underwent patch motion correction and patch CTF estimation in cryoSPARC Live using default parameters. 3,392 movies were discarded based on quality and resolution of the CTF fit, calculated defocus, total motion, and relative ice thickness, leaving 12,602 for subsequent work. The cryoSPARC blob picker was used to pick 18,955,798 particles using a circular 60-120 Å blob size, an NCC score above 0.3 and local power between -76 and 1301. Particles were extracted with a box size of 416 pixels and binned by 2 (final box size = 208 pixels). The first 100,000 particles were used for an initial 3-class *ab initio* reconstruction. The full particle set was then washed twice using heterogenous refinement seeded with the 3 volumes from the multiclass *ab initio*. The remaining 5,929,8976 particles were re-extracted with an un binned box size of 320 pixels and used for homogeneous refinement with default parameters. This was followed by i) global CTF refinement while fitting tilt, trefoil, tetrafoil and anisotropic magnification, ii) homogenous refinement, iii) local CTF refinement and iv) non-uniform refinement (4), ultimately giving a reconstruction at 2.58 Å, as judged by gold standard Fourier shell coefficient with a 0.143 cutoff.

*Model building and refinement.* A homology model was generated (AlphaFold3)[13] and docked into a sharpened map with Phenix (phenix.autosharpen, phenix.dock_in_map)[14]. The docked model was then rigid body fit by domain (phenix.real_space_refine)[15,16] , and again using 20 smaller segments identified by the TLS Motion server. The model was then completed using iterative building in Coot[17] and real space refinement (phenix.real_space_refine) in Phenix[16,18]. Phenix real-space refinement included global minimization, local grid search, atomic displacement parameters, along with secondary structure, Ramachandran and rotamer outlier restraints. The model was deposited in the Protein Data Bank with accession code PDB ID 9D26 and the map was deposited in the Electron Microscopy Data Bank with accession code EMD-46483. Metrics for model and map validation are presented in Supplementary Table 3. Figures were prepared with Pymol[19], Chimera and ChimeraX[20,21]. Protoporphyrin IX was docked to the HmuS structure using Autodock Vina as previously described.

**Proteomics analysis of excised SDS-PAGE gel bands.** The identities of the expressed protein (158 kDa), its major contaminant band (60 kDa), and a band at 45 kDa which was part of the IEC fraction but later removed by SEC purification were verified by mass spectrometry (**Arkansas IDeA facility**). SDS-PAGE gel bands were excised and subjected to in-gel trypsin digestion. Gel segments were destained in 50% methanol (Fisher), 50 mM ammonium bicarbonate (Sigma-Aldrich), followed by reduction in 10 mM Tris[2-carboxyethyl]phosphine (Pierce) and alkylation in 50 mM iodoacetamide (Sigma-Aldrich). Gel slices were then dehydrated in acetonitrile (Fisher), followed by addition of 100 ng porcine sequencing grade modified trypsin (Promega) in 50 mM ammonium bicarbonate (Sigma-Aldrich) and incubation at 37 $^{o}$C for 12-16 h. Peptide products were then acidified in 0.1% formic acid (Pierce).

Tryptic peptides were separated by reverse phase XSelect CSH C18 2.5 mm resin (Waters) on an in-line 150 x 0.075 mm column using a nanoAcquity ultrahigh pressure liquid chromatography (UPLC) system (Waters). Peptides were eluted using a 60 min gradient from 98:2 to 65:35 buffer A:B ratio. [Buffer A = 0.1% formic acid, 0.5% acetonitrile; buffer B = 0.1% formic acid, 99.9% acetonitrile.] Eluted peptides were ionized by electrospray (2.4 kV) followed by MS/MS analysis using higher-energy collisional dissociation (HCD) on an Orbitrap Fusion Tribrid mass spectrometer (Thermo) in top-speed data-dependent mode. MS data were acquired using the FTMS analyzer in profile mode at a resolution of 240,000 over a range of 375 to 1500 m/z. Following HCD activation, MS/MS data were acquired using the ion trap analyzer in centroid mode and normal mass range with precursor mass-dependent normalized collision energy between 28.0 and 31.0. Proteins were identified by database search using Mascot (Matrix Science) with a parent ion tolerance of 3 ppm and a fragment ion tolerance of 0.5 Da. Scaffold (Proteome Software) was used to verify MS/MS based peptide and protein identifications. Peptide identifications were accepted if they could be established with less than 1.0% false discovery by the Scaffold Local FDR algorithm. Protein identifications were accepted if they could be established with less than 1.0% false discovery and contained at least 2 identified peptides. Protein probabilities were assigned by the Protein Prophet algorithm[22]. Label-free quantitation (LFQ) using signal intensities for average-of-3-most-abundant-peptides per protein, normalized to the number of identifiable peptides for a given protein, was performed using the MassQuant software. The resulting iBAQ metric was computed for the most abundant proteins (Fig S10). (**See Supplementary File 01.**)

**Bottom-up proteomics data measured from partially pure (IEC) and more completely purified (IEC and SEC) HmuS.** Bottom-up proteomic analyses of protein fractions. To identify a possible co-purifying reductase, protein identities and relative abundances were assessed for IEC and IEC-SEC purified HmuS (**University of Notre Dame Mass Spectrometry Facility**). The yellow 20-30 kDa contaminant fractions isolated by SEC were also pooled, centrifuge concentrated (10 kDa MWCO), and analyzed. Samples were dissolved in water at a total protein concentration of 1 µg/µL. Proteins were reduced (100 mM dithiothreitol, MP Biomedicals, LLC) in 25 mM ammonium bicarbonate then heated for 30 min, 65 °C. Solutions were cooled to room temperature then alkylated in the dark for 30 min following the addition of iodoacetamide (Sigma) (40 mM). Trypsin digestion proceeded overnight at 37 °C following the addition of 25 µL of 0.1 ng µL-1 Trypsin Gold (Promega) in 25 mM ammonium bicarbonate. 50 µL of digested solution was desalted via 5 passes of 10 µL using C18 Tips (Pierce). Fractions eluted in 50:50 water:acetonitrile with 0.1% trifluoroacetic acid were evaporated to dryness on a speedvac. Dried samples were reconstituted with 20 µL of 96:4 water:acetonitrile with 0.1% formic acid for subsequent injection on the nanocolumn.

1 µL of tryptic peptides was resolved with nano-scale ultrahigh pressure liquid chromatography (nanoUPLC) coupled with tandem mass spectrometry using a system composed of a Waters M-Class in-line with a Q-Exactive HF mass spectrometer (Thermo Scientific). Solvent A (0.1% formic acid in water, Burdick & Jackson) and solvent B (0.1% formic acid in acetonitrile, Burdick & Jackson) were used as the mobile phase. Peptides were eluted from an Acquity BEH C18 Column, 1.7-µm particle size, 300 Å (Waters) column (100 µm inner diameter × 100 mm long) using a 48-min gradient at a flow rate of 0.9 µL/min (4% B for 8.1 min, 4-7% B 10.0 min, 7-33% B 10-30 min, 33–90% B 30-33 min, 90% B for 3 min, 90-4% B 36-37 min, 4% B 37.1-48 min to equilibrate the column). Data were collected in positive ionization mode. Peaks Online (Bioinformatics Solutions) software was used to identify proteins from tandem mass spectra of

peptide ions. The E. coli K12 protein database was modified to include the sequence of the expressed HmuS protein. Two missed cleavages by trypsin were permitted in the database search which included carbamidomethylation (C) as a fixed modification along with oxidation (M) and deamidation (N,Q) as variable modifications. The mass tolerance for precursor ions was set to 0.010 Daltons and mass tolerance for fragment ions set to 0.8 Da. (**See Supplementary File 02.**)

**Supplementary Figures**

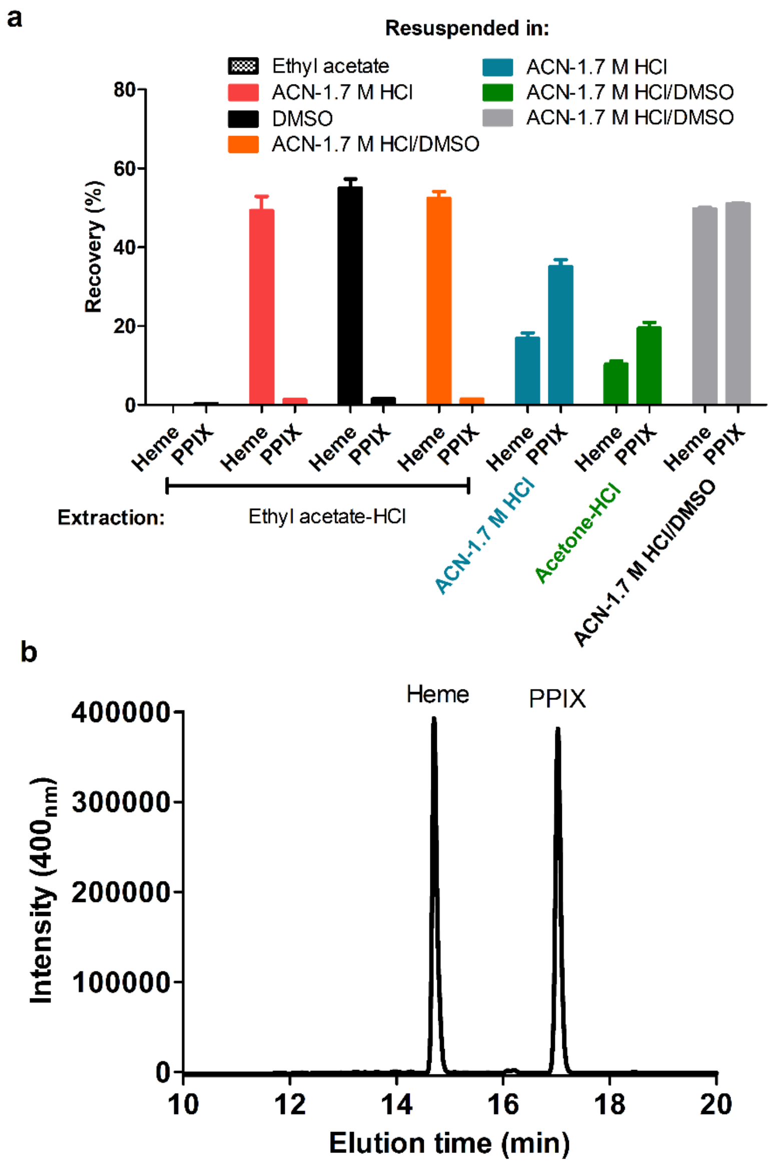

**Figure S1. Validating extraction solvents.** (A) Efficiency of heme and PPIX recovery (plotted as % recovery = [compound measured]/(100 µM + [concentration of compound in no-standard-added control sample]). Solvents used to extract heme and PPIX from the *Escherichia coli* cell pellet are listed in the legend, where ACN = acetonitrile, HCl = 12M hydrochloric acid, and DMSO = dimethylsulfoxide (ethyl acetate, acetone, ACN:1.7 M HCl (82:18, v/v), DMSO and ACN:1.7 M HCl:DMSO (41:9:50, v/v/v). The solvents used for the extraction of heme/PPIX from *E. coli* cells are indicated on the X axis and the solvents used to resuspend the dried samples are indicated with colored bars. (B) Typical HPLC trace illustrating appearance of heme and PPIX (10 µM) and their elution time at 14.7 and 17 min, respectively, wavelength at 400 nm. The chromatography was carried out using a Hypersil GOLD™ column (Thermo Scientific™, 4.6 mm x 250 mm, 5 µm particle size), linear gradient with the solution A (ultrapure water + 0.1% trifluoroacetic acid, TFA) and solution B (Acetonitrile + 0.1% TFA), flow rate of 1 mL $min^{-1}$, oven temperature at 25 °C.

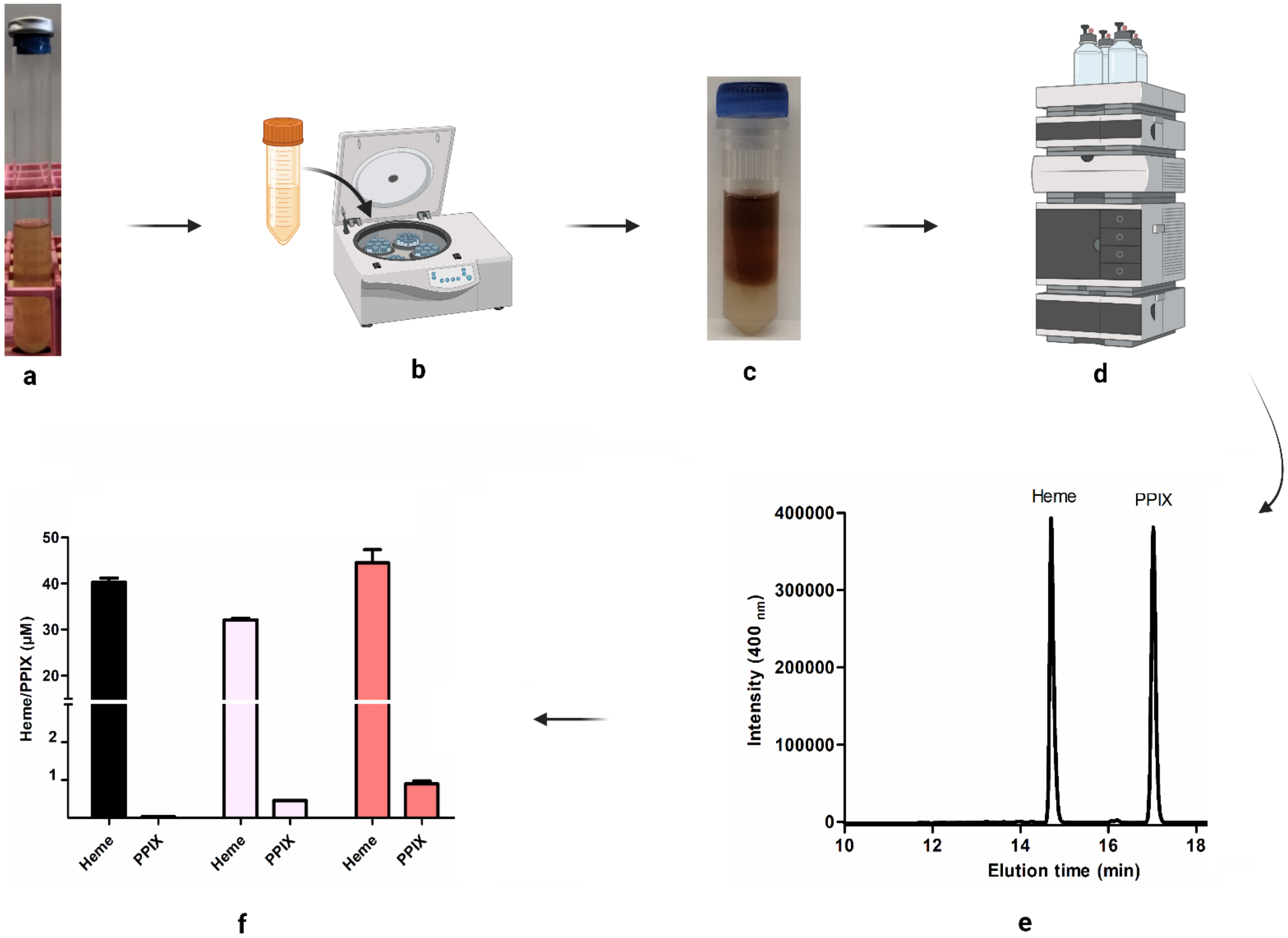

**Figure S2. Schematic illustrating methods for growing *B theta* cells and quantifying heme and PPIX extracted from cell pellets.** (a) Cultivation of *B. thetaiotaomicron* in 10 mL Balch type, crimp-topped tubes containing minimal medium and 15 µM hemin, ±BPS, under an atmosphere composed of 2.5% $H_2$ and 97.5% $N_2$, and incubated at 37 °C, 150 rpm. (b) Bacterial cells were pelleted by centrifugation (11,900 x g, 4 °C, 15 min), washed twice in ultrapure water, and resuspended (0.12 g mL$^{-1}$) in the extraction solvent ACN:HCl:DMSO (41:9:50, v/v/v). (c) The cell suspensions were then transferred into FastPrep Lysis B-matrix tubes and lysed using a FastPrep 24 5g instrument (2 cycles: 6.0 meters/second for 40 sec). (d) Samples were centrifuged (9,600 x g, 25 °C, 15 min) and quantified by peak integration using HPLC. The chromatography was carried out using a Hypersil GOLD™ column (Thermo Scientific™, 4.6 mm x 250 mm, 5 µm particle size) with a linear gradient using the solution A (ultrapure water + 0.1% TFA) and solution B (Acetonitrile + 0.1% TFA), flow rate of 1 mL min$^{-1}$, and oven temperature at 25 °C. (e) Heme and PPIX eluted at 14.7 and 17 min, respectively. (f) Quantification of heme and PPIX was performed using a standard curve (generated by plotting standard concentrations against peak areas at 400 nm) and results were presented as bar graphs.

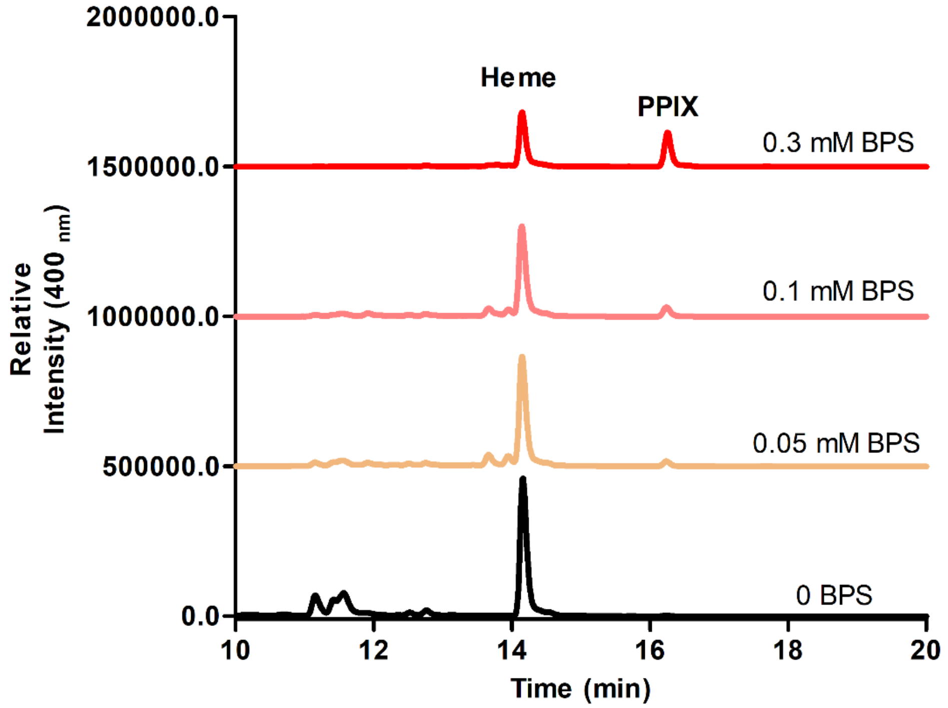

**Figure S3. Representative HPLC data illustrating quantification of heme and PPIX extracted from *B theta* cells grown in minimal medium with increasing concentrations of BPS.** Representative HPLC data used in generating the data plotted in bar chart in Figure 1a are shown. Numerical data were measured from 3 biological replicates, and values are given in Table S1.

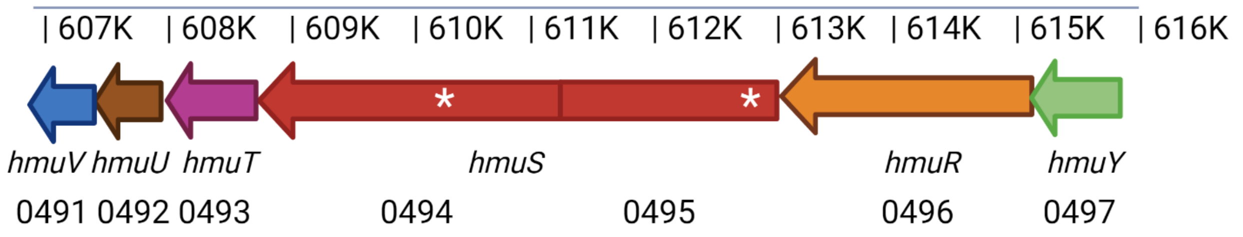

**Figure S4. Composition of the *hmu* operon from B theta VPI-5482 and locations of transposon insertions.** The magnified region of the complete genome between nucleotides 607K-616K (GenBank accession AE015928.1) is shown. Arrows indicate each of the *hmuYRSTUV* genes encoded on the antisense strand, relative to the nucleotide numbering shown along the top of the figure. BT loci numbers are given beneath the gene names. (Note: the *hmuS* gene was erroneously assigned two locus numbers in this genome because of a frameshift, which was later corrected.) Two *hmuS* transposon insertion mutants were generated, mapping to nucleotide positions 610,214 (*hmuS* 2, BT0495 P295-G04) and 612,897 (*hmuS* 1, BT0494 P289-C05), respectively. Approximate locations of transposon insertion are marked with a white asterisk. The transposon sequence and details of its use to generate a genomic library of mutants is described by Shiver et al[4].

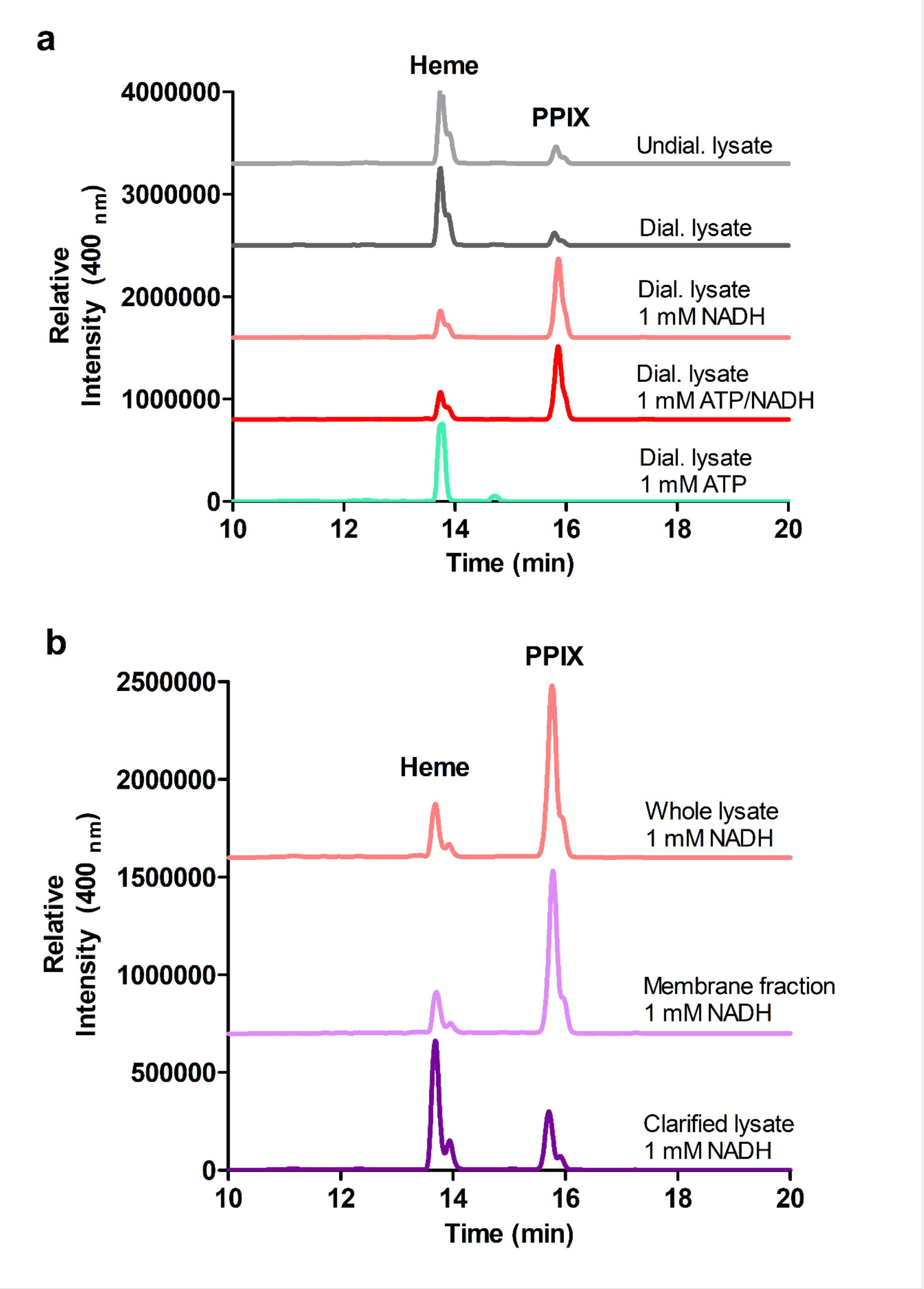

**Figure S5. Representative HPLC data illustrating the production of PPIX by *B theta* cell fractions incubated with heme and NADH or ATP.** Representative HPLC data used in generating the data plotted in bar charts in Figure 2a and b are shown. Numerical data were measured from 3 biological replicates, and values are given in Table S1.

**(A)** N-
MKKKSKIILGGCIVVAALIGLSVWNTWFS***ATKIAFVNFQTIQQGSISKANDNSFIKLSEVSLDNLDRLTSYDMVFING***
***MGLRIVEEQRQQIQQAADKGIPVYTSMATNPANNICNLDSIQQNLIRGYLSNGGKTNYRNMLNYIRKAIDGKASA***
***VPEVEDPIERPSDMLYHAGISNPDDEQEFLTVADYEKFMQENNLYKEGARKIMITGQMADATDLIKALENAGYN***
***VYPVQSMTRFMSFIEEVQPDAVINMAHGRMGDKMVDYLKRNILLFAPLTINSLVDEWENDPMGMSGGFMSQSI***
***VTPEIDGAIRPFALFAQYEDKEGLRHSYAVPERLKTFVSTIDNYLNLKTKPNFEKKVAIYYYKGPGQNALTAAGM***
***EVVPSLYNLLLRMKQEGYNISGLPANAQELGKMIQAQGAVFNAYAEGAFNDFMQNGHPELITKEQYESWVKES***
***LRPEKYQEVVDAFGEFPGNYMVTPDGKLGIARLQFGNVVLLPQNAAGSGDNSFQVVHGTDMAPPHTYIASYLW***
***MQHGFKADALIHFGTHGSLEFTPRKQVALCSNDWPDRLVGAVPHYYLYSIGNVGEGMMAKRRSYATLQSYLT***
***PPFLESSVRGIYRELMEKIKIYNNSQKANKDQESLAVKTLTVKMGIHRDLGLDSMANKPYTEDEIARVENFAEEL***
***ATEKITGQLYTMGVPYEPERITSSVYAMATEPIAYSLFALDKQRGKATESAEKHRSVFTQQYLMPARLLVERLM***
***ANPSLATDELICHTAGITPQELAKARQIEAERNAPKGMMAMMMAAAAKKDQADNEPSGNGHPASAKMEKGP***
***HGKMPAGMKEAMKKMGANMDPEKAMEMAKSMGASPEALKKMEASMKANKDTSTDASGKPAMAGKTEKPQ***
***GMSAMMAAMGKAPKEYSKEEVEFALAVAEVERTIKNVGNYKNALLTSPEEELSSLMNALKGGYTAPTPGGDPI***
***ANPNTLPTGRNMYAINAEATPTEAWEKGIALAKQTIDRYKQRHNDSIPRKVSYTLWSSEFIETGGATIAQVLYML***
***GVEPVRDAFGRVSDLKLIPSTELGRPRIDVVVQTSGQLRDLAASRLFLINRAVEMAAAAKDDKYENQVASSVIE***
***AERVLTEKGLSPKDAREISTFRVFGGANGMYGTGIQEMVESGDRWENESEIADTYLNNMGAYYGSEKNWEVF***
***QKFAFEAALTRTDVVVQPRQSNTWGALSLDHVYEFMGGMNLAVRNVTGKDPDAYLSDYRNRNHMKMQELKE***
***AVGVESRTTILNPTYIKEKMKGGASSASEFAEVITNTYGWNVMKPAAIDKELWDNIYNVYVKDELNLGVKQYFE***
***QQNPAALEEMTAVMLESARKGLWQASEEQVAELSKLHTEIVNTYRPSCSGFVCDNAKLRDFIASKADAQTATQ***
***YKENISKIR***AKASGSNKGVVMKKEEMNQTAENQTNTLSNVAVGIAVIIVILALILFVRKRRKSSQM-C

**(B)**

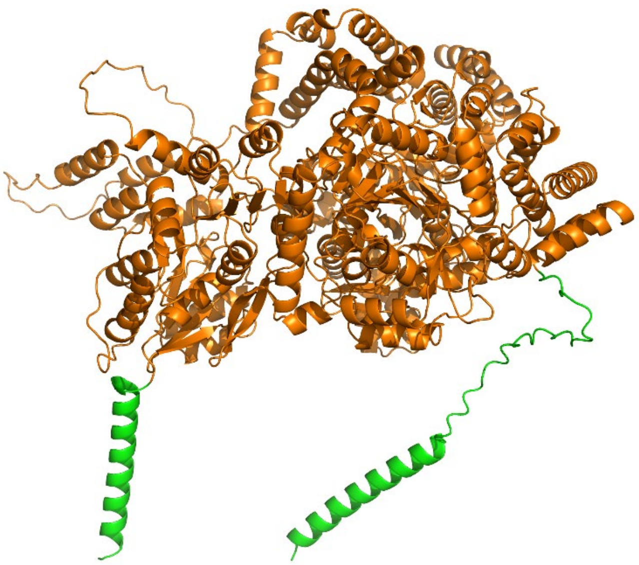

**(C)**

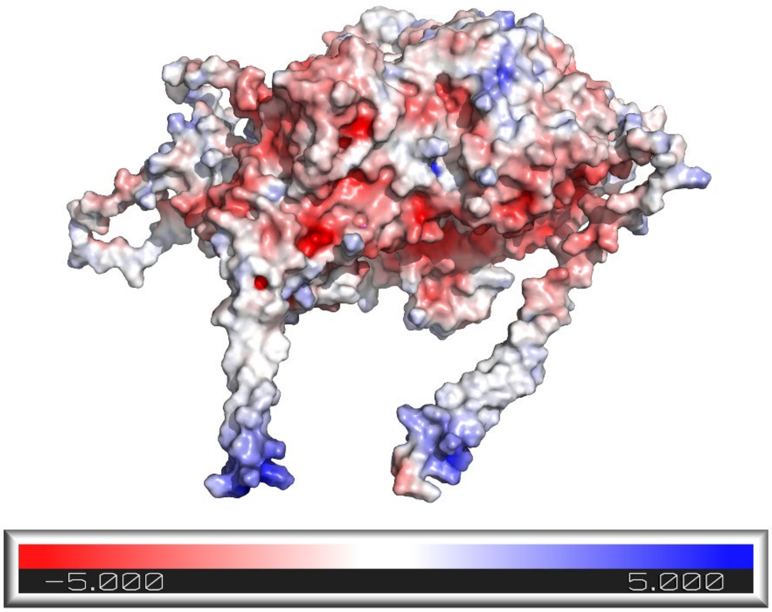

**Figure S6. Annotated protein sequence of HmuS and AlphaFold2-predicted structure used in designing the expression construct.** (A) The full-length 1463 coding sequence (RefSeq: WP_008765019.1) was truncated to eliminate predicted single-transmembrane spanning alpha helices (prediction using Phobius, https://phobius.sbc.su.se/cgi-bin/predict.pl) at the N- and C-termini (residues 1-29 and 1406-1463, highlighted in green; N- and C-termini are labeled). The remaining 1377 amino acid sequence (orange, underlined/bold-italics) was reverse-translated by GenScript and used to generate a synthetic gene for expression in *E. coli* strain BL21DE3. (B) AlphaFold2 structure of the full-length native HmuS, color-coded to match the expressed/unexpressed regions delineated in (A). The N-terminal His tag is not shown. A series of residues predicted to form a poorly structured coil were omitted at the C-terminus to avoid aggregation of the expressed protein. (C) Electrostatic potential surface of the AlphaFold structure was generated using the Adaptive Poisson-Boltzmann Solver (APBS) tool in Pymol. Negative surfaces are in red, positive in blue, and neutral/nonpolar in white. Contour levels are +/- 5 kT/e. Note the pair of hydrophobic helices predicted to span the membrane, and the large stretch of negative charge on the protein's surface between them.

5'-
tggcgaatgggacgcgccctgtagcggcgcattaagcgcggcgggtgtggtggttacgcgcagcgtgaccgctacacttgccagcgccctagcgcccgctcctttcgctttc
ttcccttcctttctcgccacgttcgccggctttccccgtcaagctctaaatcgggggctccctttagggttccgatttagtgctttacggcacctcgaccccaaaaaacttgattagg
gtgatggttcacgtagtgggccatcgccctgatagacggtttttcgccctttgacgttggagtccacgttctttaatagtggactcttgttccaaactggaacaacactcaaccctat
ctcggtctattcttttgatttataagggattttgccgatttcggcctattggttaaaaaatgagctgatttaacaaaaatttaacgcgaattttaacaaaatattaacgtttacaatttca
ggtggcacttttcggggaaatgtgcgcggaacccctatttgtttatttttctaaatacattcaaatatgtatccgctcatgaattaattcttagaaaaactcatcgagcatcaaatga
aactgcaatttattcatatcaggattatcaataccatatttttgaaaaagccgtttctgtaatgaaggagaaaactcaccgaggcagttccataggatggcaagatcctggtatc
ggtctgcgattccgactcgtccaacatcaatacaacctattaatttcccctcgtcaaaaataaggttatcaagtgagaaatcaccatgagtgacgactgaatccggtgagaat
ggcaaaagtttatgcatttctttccagacttgttcaacaggccagccattacgctcgtcatcaaaatcactcgcatcaaccaaaccgttattcattcgtgattgcgcctgagcgag
acgaaatacgcgatcgctgttaaaaggacaattacaaacaggaatcgaatgcaaccggcgcaggaacactgccagcgcatcaacaatattttcacctgaatcaggatatt
cttctaatacctggaatgctgttttcccggggatcgcagtggtgagtaaccatgcatcatcaggagtacggataaaatgcttgatggtcggaagaggcataaattccgtcagcc
agtttagtctgaccatctcatctgtaacatcattggcaacgctacctttgccatgtttcagaaacaactctggcgcatcgggcttcccatacaatcgatagattgtcgcacctgatt
gcccgacattatcgcgagcccatttatacccatataaatcagcatccatgttggaatttaatcgcggcctagagcaagacgtttcccgttgaatatggctcataacaccccttgt
attactgtttatgtaagcagacagttttattgttcatgaccaaaatcccttaacgtgagttttcgttccactgagcgtcagaccccgtagaaaagatcaaaggatcttcttgagatcc
tttttttctgcgcgtaatctgctgcttgcaaacaaaaaaaccaccgctaccagcggtggtttgtttgccggatcaagagctaccaactctttttccgaaggtaactggcttcagcag
agcgcagataccaaatactgtccttctagtgtagccgtagttaggccaccacttcaagaactctgtagcaccgcctacatacctcgctctgctaatcctgttaccagtggctgct
gccagtggcgataagtcgtgtcttaccgggttggactcaagacgatagttaccggataaggcgcagcggtcgggctgaacggggggttcgtgcacacagcccagcttgga
gcgaacgacctacaccgaactgagatacctacagcgtgagctatgagaaagcgccacgcttcccgaagggagaaaggcggacaggtatccggtaagcggcagggtc
ggaacaggagagcgcacgagggagcttccagggggaaacgcctggtatctttatagtcctgtcgggtttcgccacctctgacttgagcgtcgatttttgtgatgctcgtcaggg
gggcggagcctatggaaaaacgccagcaacgcggcctttttacggttcctggccttttgctggccttttgctcacatgttctttcctgcgttatcccctgattctgtggataaccgtat
taccgcctttgagtgagctgataccgctcgccgcagccgaacgaccgagcgcagcgagtcagtgagcgaggaagcggaagagcgcctgatgcggtattttctccttacgc
atctgtgcggtatttcacaccgcatatatggtgcactctcagtacaatctgctctgatgccgcatagttaagccagtatacactccgctatcgctacgtgactgggtcatggctgc
gccccgacacccgccaacacccgctgacgcgccctgacgggcttgtctgctcccggcatccgcttacagacaagctgtgaccgtctccgggagctgcatgtgtcagaggtt
ttcaccgtcatcaccgaaacgcgcgaggcagctgcggtaaagctcatcagcgtggtcgtgaagcgattcacagatgtctgcctgttcatccgcgtccagctcgttgagtttctc
cagaagcgttaatgtctggcttctgataaagcgggccatgttaagggcggtttttttcctgtttggtcactgatgcctccgtgtaagggggatttctgttcatgggggtaatgataccg
atgaaacgagagaggatgctcacgatacgggttactgatgatgaacatgcccggttactggaacgttgtgagggtaaacaactggcggtatggatgcggcgggaccaga
gaaaaatcactcagggtcaatgccagcgcttcgttaatacagatgtaggtgttccacagggtagccagcagcatcctgcgatgcagatccggaacataatggtgcagggc
gctgacttccgcgtttccagactttacgaaacacggaaaccgaagaccattcatgttgttgctcaggtcgcagacgttttgcagcagcagtcgcttcacgttcgctcgcgtatcg
gtgattcattctgctaaccagtaaggcaaccccgccagcctagccgggtcctcaacgacaggagcacgatcatgcgcacccgtggggccgccatgccggcgataatggc
ctgcttctcgccgaaacgtttggtggcgggaccagtgacgaaggcttgagcgagggcgtgcaagattccgaataccgcaagcgacaggccgatcatcgtcgcgctccag
cgaaagcggtcctcgccgaaaatgacccagagcgctgccggcacctgtcctacgagttgcatgataaagaagacagtcataagtgcggcgacgatagtcatgccccgc
gcccaccggaaggagctgactgggttgaaggctctcaagggcatcggtcgagatcccggtgcctaatgagtgagctaacttacattaattgcgttgcgctcactgcccgcttt
ccagtcgggaaacctgtcgtgccagctgcattaatgaatcggccaacgcgcggggagaggcggtttgcgtattgggcgccagggtggtttttcttttcaccagtgagacggg
caacagctgattgcccttcaccgcctggccctgagagagttgcagcaagcggtccacgctggtttgccccagcaggcgaaaatcctgtttgatggtggttaacggcgggata
taacatgagctgtcttcggtatcgtcgtatcccactaccgagatatccgcaccaacgcgcagcccggactcggtaatggcgcgcattgcgcccagcgccatctgatcgttgg
caaccagcatcgcagtgggaacgatgccctcattcagcatttgcatggtttgttgaaaaccggacatggcactccagtcgccttcccgttccgctatcggctgaatttgattgcg
agtgagatatttatgccagccagccagacgcagacgcgccgagacagaacttaatgggcccgctaacagcgcgatttgctggtgacccaatgcgaccagatgctccacg
cccagtcgcgtaccgtcttcatgggagaaaataatactgttgatgggtgtctggtcagagacatcaagaaataacgccggaacattagtgcaggcagcttccacagcaatg
gcatcctggtcatccagcggatagttaatgatcagcccactgacgcgttgcgcgagaagattgtgcaccgccgctttacaggcttcgacgccgcttcgttctaccatcgacac
caccacgctggcacccagttgatcggcgcgagatttaatcgccgcgacaatttgcgacggcgcgtgcagggccagactggaggtggcaacgccaatcagcaacgactg
tttgcccgccagttgttgtgccacgcggttgggaatgtaattcagctccgccatcgccgcttccactttttcccgcgttttcgcagaaacgtggctggcctggttcaccacgcggg
aaacggtctgataagagacaccggcatactctgcgacatcgtataacgttactggtttcacattcaccaccctgaattgactctcttccgggcgctatcatgccataccgcgaa
aggttttgcgccattcgatggtgtccgggatctcgacgctctcccttatgcgactcctgcattaggaagcagcccagtagtaggttgaggccgttgagcaccgccgccgcaag
gaatggtgcatgcaaggagatggcgcccaacagtcccccggccacggggcctgccaccatacccacgccgaaacaagcgctcatgagcccgaagtggcgagcccg
atcttccccatcggtgatgtcggcgatataggcgccagcaaccgcacctgtggcgccggtgatgccggccacgatgcgtccggcgtagaggatcgagatctcgatcccgc
gaaattaatacgactcactataggggaattgtgagcggataacaattcccctctagaaataattttgtttaactttaaga**aggaga**tatacccat**atggcaacaaaaatagct**
**ttcgtaaattttcaaacgatccagcaaggttctatcagcaaagccaacgataatagctttatcaagctgagcgaagtttctctcgataacttagaccgtctgacga**
**gctatgatatggtgttcatcaatggcatgggcttgcgtattgtggaagaacagcgtcaacaaatccaacaggccgccgacaagggtattccggtttacaccag**
**catggcgactaatccggcgaacaatatttgcaatctggatagcatccagcagaaccttatccgtggttacctgtccaacggcggcaagaccaactaccgtaac**
**atgctcaactatatccgtaaagcaatcgatggtaaggcctctgccgtgccggaggtagaggacccgattgaacgtccgtccgatatgttgtaccatgcaggca**
**tcagcaacccggatgatgagcaagagttcctgaccgttgcggactacgaaaagtttatgcaggaaaacaatctgtacaaagagggtgcgagaaagattatga**
**ttaccggccagatggcagacgcgaccgatcttattaaagcacttgagaacgcaggttataacgtctatccggtccaaagcatgacccgttttatgtccttcattga**
**agaggtgcagccggacgcggtgatcaatatggcacatggtcggatgggggataagatggtcgattatttgaaagcgcgcaatatcctgttgttcgcaccgctg**
**accattaacagtctggtggatgagtgggaaaacgatccgatgggtatgagcggcggttttatgagccagagcatcgttactccggagatcgacggcgcaatc**
**cgtccgtttgcactgttcgcgcagtatgaggataaagagggtctacgccacagctacgcggtgccggaacgtttgaaaacttttgtcagcaccattgacaacta**
**cttgaacttgaagaccaagccgaactttgaaaagaaagttgcgatttattactacaagggtccgggtcaaaatgctctgacggcagcgggcatggaggttgta**
**ccgagcctctacaacctgttgctgcgtatgaagcaggaaggttataacattagcggactgcctgcaaatgcgcaagaactgggtaagatgattcaggctcagg**
**gcgctgtgtttaacgcatacgcggaaggtgcgttcaacgacttcatgcagaatggccacccagagctgattaccaaagagcagtatgagtcttgggttaaaga**

**gagcttgcgcccagagaagtaccaggaagttgttgatgcgtttggagaatttccgggtaactacatggtgacaccagacggcaaattaggcatcgcacgtctg
caatttggcaacgtggtgctgctgccgcaaaacgcggccggttcgggcgacaacagcttccaggttgttcatggtacagacatggctccgccacacacctac
attgcatcgtacctgtggatgcagcatggcttcaaagcggacgcgctgatacactttggtacccatggcagcctcgagttcaccccgagaaagcaagttgcac
tttgcagcaacgactggccggatcgtttggttggggcagtcccgcactattacctttacagcatcggtaatgttggtgaaggcatgatggccaagcgccgttcgt
acgcgaccctgcagagctacttgaccccgccttttctggagagcagcgtgcgtggtatctatcgggaattaatggaaaaaatcaaaatttataacaactcccag
aaagcgaacaaagatcaagaaagcttggcggtcaagaccctgacggttaaaatgggtattcaccgtgatctaggtctggattctatggctaacaaaccgtata
ccgaagacgaaatcgctcgcgttgagaacttcgcggaagaattggcgaccgagaagatcaccggtcaattgtacaccatgggcgtgccgtacgaaccgga
gcgcattacgtcgagcgtgtatgccatggcgaccgagccgattgcgtacagcctcttcgcactggataagcagcgtggcaaagcgaccgaatcggccgaa
aagcaccgctctgtttttacccagcaatacctgatgccggcacgcctcttggttgagcgtctgatggctaacccgagcctcgccacggacgaactgatatgcc
ataccgctggcattaccccgcaggagctggcgaaagcccgtcaaatcgaagctgagcgcaatgcgccaaaaggaatgatggcgatgatgatggccgcgg
cagcgaaaaaggaccaggcagataacgagccgagcggcaacggtcacccggcgtccgcgaagatggaaaaaggccctcacggcaagatgccggccg
gcatgaaggaggcaatgaagaaaatgggcgctaatatggatccggaaaaggcgatggaaatggccaaaagcatgggtgctagcccagaagcgttgaaaa
aaatggaagccagcatgaaagcgaataaggacacctccactgatgcgtccggcaagccggctatggctggcaaaacggagaagccacaaggtatgtctg
caatgatggcggctatgggtaaggcgccaaaagaatattccaaagaagaggtggagttcgctctggctgtggccgaggttgaacgtaccattaaaaatgttgg
taattacaagaatgcgttgctgacgagcccggaggaggaattgtcctctctgatgaatgctcttaagggcggctacacggcgccgaccccaggtggtgaccc
tatcgcgaacccgaacaccctgccgaccggtagaaacatgtacgcgatcaacgccgaagcgacccccaccgagtcagcgtgggaaaagggcattgccct
ggcgaagcaaaccattgaccgctataagcaacgtcacaatgatagcatcccgcgcaaggtgagctacaccctgtggagctcggagttcatcgagacaggc
ggtgcaacgatcgcccaggttctgtacatgctgggggtggagccggttcgtgatgcgttcggccgtgtgtccgatctgaaactgattccgtctaccgagctgg
gtcgtccgagaattgacgtggtggtccagactagcggacagctgcgtgacctggccgcgtctcgtttatttctgatcaaccgtgcagtggaaatggcagctgc
ggcgaaggacgacaaatacgaaaaccaggtggcgagcagcgtcatcgaggctgaacgtgtcctgaccgaaaaaggcctgtccccgaaagatgcgcgcg
aaatttccacctttcgtgttttcggtggtgcgaacggtatgtatggtaccggtatccaagagatggttgagtccggcgaccgctgggaaaatgagagcgaaatc
gccgacacatacctgaacaacatgggtgcatattacggcagcgagaagaactgggaagtgtttcagaagttcgccttcgaagcggcgctgactcgcacgga
cgttgtggtgcaaccgcgtcaatctaatacctggggtgcgctgagcctggaccatgtgtacgagttcatgggtggaatgaacttggcggtgcgcaatgtcacc
ggcaaggacccggatgcctatttaagcgactatagaaaccgcaaccacatgaagatgcaagagcttaaagaggcagttggcgttgagtcccgtaccaccatt
ctgaatccgacctatatcaaagagaagatgaaaggtggtgcgtcctctgcatcagaattcgctgaggttattacgaacacgtatggctggaatgttatgaaacc
ggctgcaatcgataaagagctgtgggataacatctataacgtgtatgtgaaagacgagttaaaccttggcgttaaacaatacttcgagcagcagaacccggct
gctttggaggaaatgaccgcggtgatgctggaaagcgcccgtaaaggtctgtggcaggcctccgaggaacaagtggcagagctgagcaaactgcataccg
aaatcgttaatacctatcgtccgtcttgtagcggcttcgtttgtgataatgcgaaactgcgtgactttatcgccagcaaggcggacgctcagactgcgacgcagt
acaaggaaaacatctcgaagattagataatga**ctcgagcaccaccaccaccaccactgagatccggctgctaacaaagcccgaaaggaagctgagttggctgctgc
caccgctgagcaataactagcataaccccttggggcctctaaacgggtcttgaggggttttttgctgaaaggaggaactatatccggat-3’

**(B)**

MATKIAFVNFQTIQQGSISKANDNSFIKLSEVSLDNLDRLTSYDMVFINGMGLRIVEEQRQQIQQAADKGIPVYTSMATNPA
NNICNLDSIQQNLIRGYLSNGGKTNYRNMLNYIRKAIDGKASAVPEVEDPIERPSDMLYHAGISNPDDEQEFLTVADYEKFM
QENNLYKEGARKIMITGQMADATDLIKALENAGYNVYPVQSMTRFMSFIEEVQPDAVINMAHGRMGDKMVDYLKARNILLF
APLTINSLVDEWENDPMGMSGGFMSQSIVTPEIDGAIRPFALFAQYEDKEGLRHSYAVPERLKTFVSTIDNYLNLKTKPNFE
KKVAIYYYKGPGQNALTAAGMEVVPSLYNLLLRMKQEGYNISGLPANAQELGKMIQAQGAVFNAYAEGAFNDFMQNGHP
ELITKEQYESWVKESLRPEKYQEVVDAFGEFPGNYMVTPDGKLGIARLQFGNVVLLPQNAAGSGDNSFQVVHGTDMAPP
HTYIASYLWMQHGFKADALIHFGTHGSLEFTPRKQVALCSNDWPDRLVGAVPHYYLYSIGNVGEGMMAKRRSYATLQSYL
TPPFLESSVRGIYRELMEKIKIYNNSQKANKDQESLAVKTLTVKMGIHRDLGLDSMANKPYTEDEIARVENFAEELATEKITG
QLYTMGVPYEPERITSSVYAMATEPIAYSLFALDKQRGKATESAEKHRSVFTQQYLMPARLLVERLMANPSLATDELICHTA
GITPQELAKARQIEAERNAPKGMMAMMMAAAAKKDQADNEPSGNGHPASAKMEKGPHGKMPAGMKEAMKKMGANMD
PEKAMEMAKSMGASPEALKKMEASMKANKDTSTDASGKPAMAGKTEKPQGMSAMMAAMGKAPKEYSKEEVEFALAVA
EVERTIKNVGNYKNALLTSPEEELSSLMNALKGGYTAPTPGGDPIANPNTLPTGRNMYAINAEATPTESAWEKGIALAKQTI
DRYKQRHNDSIPRKVSYTLWSSEFIETGGATIAQVLYMLGVEPVRDAFGRVSDLKLIPSTELGRPRIDVVVQTSGQLRDLAA
SRLFLINRAVEMAAAAKDDKYENQVASSVIEAERVLTEKGLSPKDAREISTFRVFGGANGMYGTGIQEMVESGDRWENES
EIADTYLNNMGAYYGSEKNWEVFQKFAFEAALTRTDVVVQPRQSNTWGALSLDHVYEFMGGMNLAVRNVTGKDPDAYL
SDYRNRNHMKMQELKEAVGVESRTTILNPTYIKEKMKGGASSASEFAEVITNTYGWNVMKPAAIDKELWDNIYNVYVKDEL
NLGVKQYFEQQNPAALEEMTAVMLESARKGLWQASEEQVAELSKLHTEIVNTYRPSCSGFVCDNAKLRDFIASKADAQTA
TQYKENISKIR

**Figure S7. Annotated DNA sequence of 9373 nucleotide expression construct used to generate HmuS.** (A) The full-length gene encoding the HmuS protein from *Bacteroides thetaiotaomicron* (strain VPI 5482, ATCC 29148, locus numbers BT0495-BT0494, protein accession: WP_022471467.1) was codon-optimized for expression in *E. coli* and synthetically introduced between the NdeI/XhoI sites of the sense strand of vector pET28a(+) by GenScript. The ribosome binding site (RBS) is shown in red. The expressed gene sequence encoding HmuS is shown in **orange**. The sequence in **violet** at the 3’ end of the gene corresponds to two STOP codons inserted into the construct. Sequences encoding the Lac repressor protein and an enzyme providing kanamycin resistance (KanR) are encoded in the opposite direction from the *hmuS* gene, on the minus strand (not shown). (B) FASTA sequence of the expressed protein (1377 amino acids), translated from part (A). A methionine was added at the N-terminus, corresponding to the start codon.

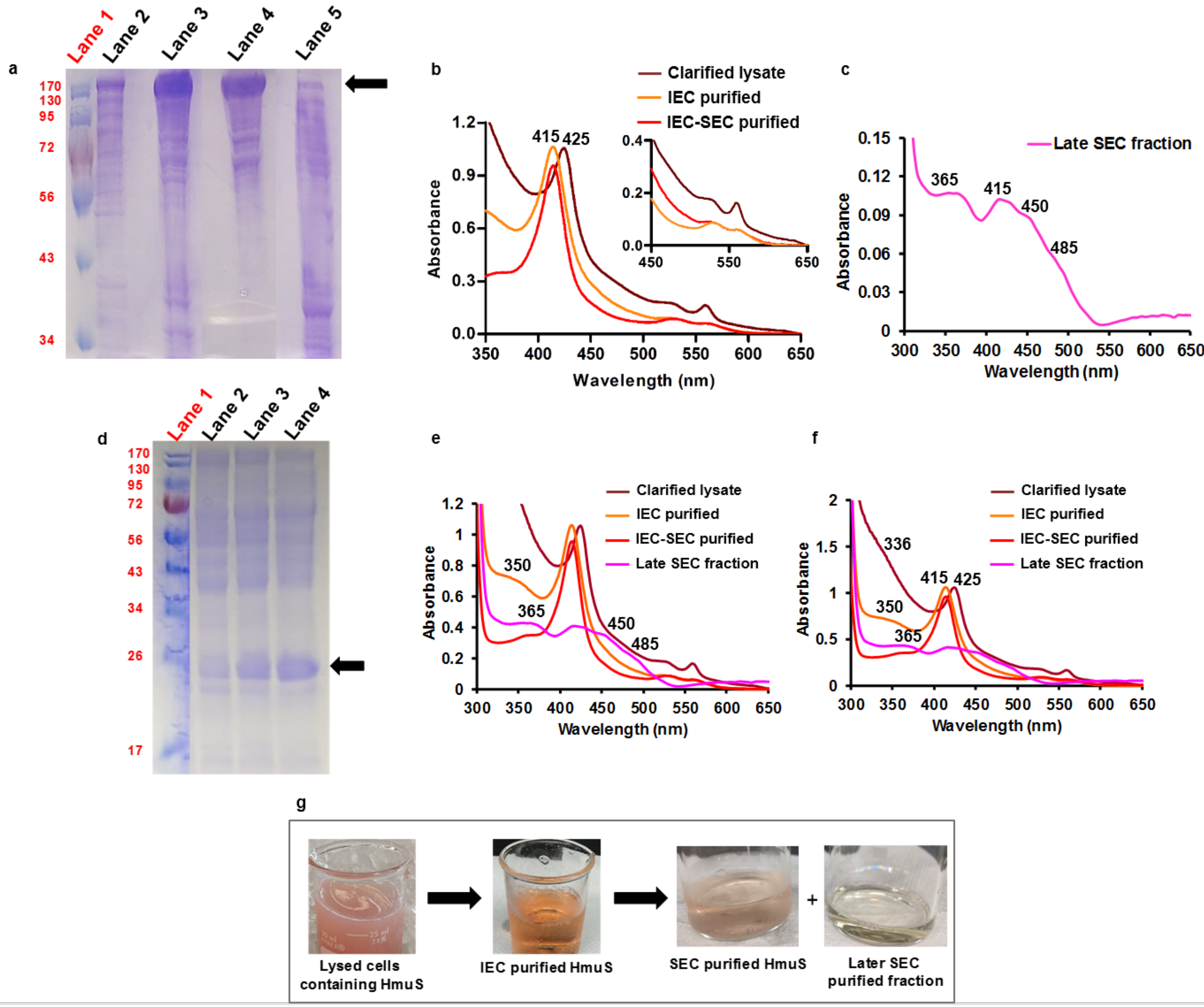

**Figure S8. Steps in the purification of recombinant HmuS illustrated by SDS-PAGE, UV/visible absorbance, and fraction color.** (a) 12% SDS-PAGE analyses of heterologously expressed HmuS in the soluble portion of the host/*E. coli* lysate (centrifuge-clarified lysate) (lane 2, 20 µg), following purification by ion exchange (lane 3, 30 µg), and after further purification by size exclusion chromatography (lane 4, 30 µg). Lane 5 contains pooled, low-molecular weight fractions that eluted after the HmuS (late SEC fraction) (90 mg total protein). The molecular weight size marker was loaded in lane 1. The arrow indicated the 158 KDa HmuS protein. (b) UV/visible absorbance spectra of heterologously expressed HmuS were measured for the same fractions as in (a), but with approximately equal [heme] in each fraction (7.5-9.5 µM as measured by the pyridine hemochromagen assay). The clarified *E. coli* lysate spectrum is shown in rust (22 mg $mL^{-1}$ total protein, roughly 10% HmuS based on (a)); following purification by ion exchange (orange line, 15 mg $mL^{-1}$ total protein, >30% HmuS); and after further purification by size exclusion chromatography (red line, 10 mg $mL^{-1}$ total protein, ≥70% HmuS). As-isolated HmuS contains substoichiometric, reduced heme. The heme oxidized and partly dissociated from the protein during purification. (c) UV/visible absorbance spectrum and (d) 12% SDS-PAGE of the late-eluting, yellow SEC fraction (16 mg $mL^{-1}$ total protein). The arrow indicates a 26 kDa contaminant protein that was commonly observed. The molecular marker was loaded in lane 1. Panels (e) and (f) show the same data as in panels (a) and (b) overlaid. The spectrum from panel (b) is shown on an expanded scale (x 4) to illustrate its peak positions. Panel (g) shows digital images of purification fractions, illustrating their colors.

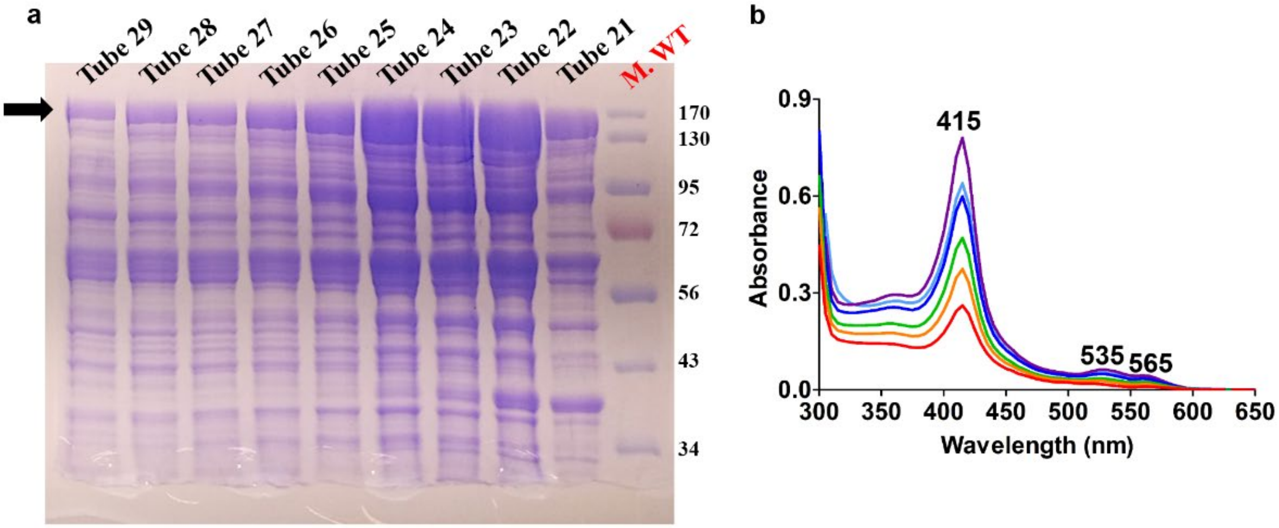

**Figure S9. HmuS expressed in *E. coli* is partially heme-bound.** HmuS was heterologously expressed in *E. coli* and the cells were pelleted and frozen as described above. The cells were thawed, resuspended, lysed, and clarified via centrifugation. A rose-colored soluble fraction resulted, from which HmuS was subsequently purified by anion exchange chromatography (DEAE Sepharose resin, Cytiva). The column was pre-equilibrated with 20 mM Tris-HCl, pH 7, before loading with the clarified lysate and eluting with the same buffer containing a linear gradient of 0-500 mM NaCl. (a) A 12% acrylamide gel illustrates 3 mL fractions collected over time. The black arrow indicates the expected position of HmuS (158 kDa) on the gel. Major contaminants were identified near 90 and 60 kDa. (b) UV/visible absorption spectra of fractions labeled tube 27-22, in descending order: red, orange, green, blue, cyan, violet. These identified a bound ferric heme (Soret band: 415 nm, Q-bands: 535 and 565 nm). The intensities of the UV/visible spectra and the band corresponding to HmuS (158 kDa, black arrow) were correlated, as expected for a heme-binding protein.

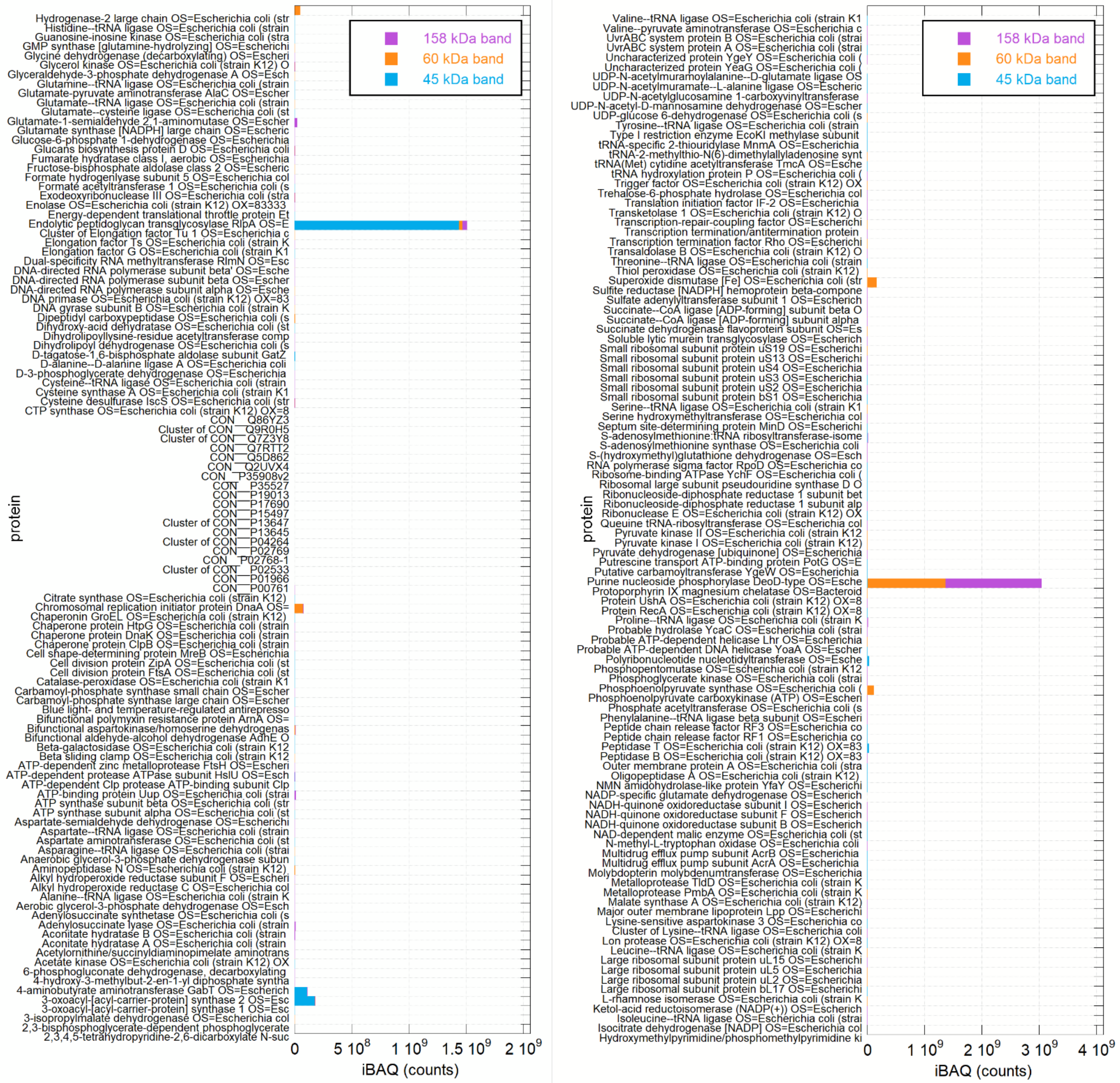

**Figure S10. Mass spectrometric analysis of heterologously expressed HmuS confirms its identity and integrity and shows its major contaminant band is principally composed of HmuS-derived peptides.** HmuS was expressed and purified by IEC as described in the text, and the IEC-purified protein analyzed by SDS-PAGE (12% acrylamide). Bands corresponding to purified HmuS (158 kDa), its major contaminant at 60 kDa, and a contaminant that was later successfully removed by SEC (45 kDa, control) were excised from the gel, proteolyzed with trypsin, and eluted from the gel matrix as described in the Supplementary Methods section above. The peptides from each band were injected onto a UPLC in-line with an Orbitrap mass spectrometer for MS identification of peptides and MS/MS analysis of their sequence. Peptides were mapped onto the *B theta* HmuS sequence or sequences from the *E. coli* host, and proteins were quantified using the iBAQ approach (plotted as superimposed columns above for the top 220 protein hits). Analysis of the 158 kDa band showed HmuS ("protoporphyrin IX $Mg^{2+}$ chelatase," indicated with an arrow on the plotted proteins and their relative amounts above) as the top hit. 91 peptides were detected in 1222 total spectra (4.7% of the total), representing 75% coverage (1073/1427 amino acids) of the HmuS sequence. Peptides were detected from across the entire sequence, with no systematic gaps. The iBAQ score relative to the summed iBAQ scores (riBAQ) indicated 73% of the band consisted of HmuS. (b) Analysis of the 60 kDa band indicated 84 HmuS peptides detected in 783 spectra, representing 70% coverage of the HmuS sequence. The riBAQ(HmuS) for this band was 37%.

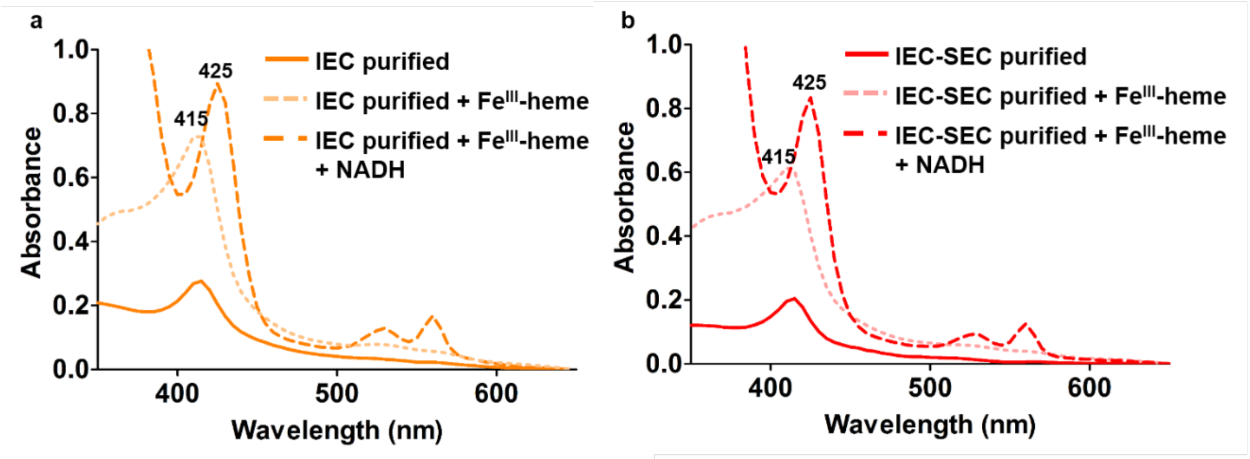

**Figure S11. UV/visible absorption spectroscopy illustrating the ferric HmuS-heme complex purified by (a) IEC and (b) IEC and SEC chromatographies, incubated with additional but substoichiometric $Fe^{III}$-heme, and then reduced with NADH.** Solid lines illustrate proteins as-purified. Lighter shade, dashed lines illustrate proteins following reconstitution with additional, substoichiometric $Fe^{III}$-heme. Darker dashed lines show proteins following anaerobic incubation with 1 mM NADH. (a) IEC purified HmuS (3.2 mg $mL^{-1}$ protein, 2.5 mM $Fe^{III}$-heme) was incubated with 4.5 μM $Fe^{III}$-heme and then reduced with 1 mM NADH. (b) IEC-SEC purified HmuS (3.2 mg $mL^{-1}$ protein, 2.0 mM $Fe^{III}$-heme) was incubated with 4.5 μM $Fe^{III}$-heme and then reduced with 1 mM NADH. All the experiments were carried out in 20 mM Tris-HCl, 250 mM NaCl, pH 7.1.

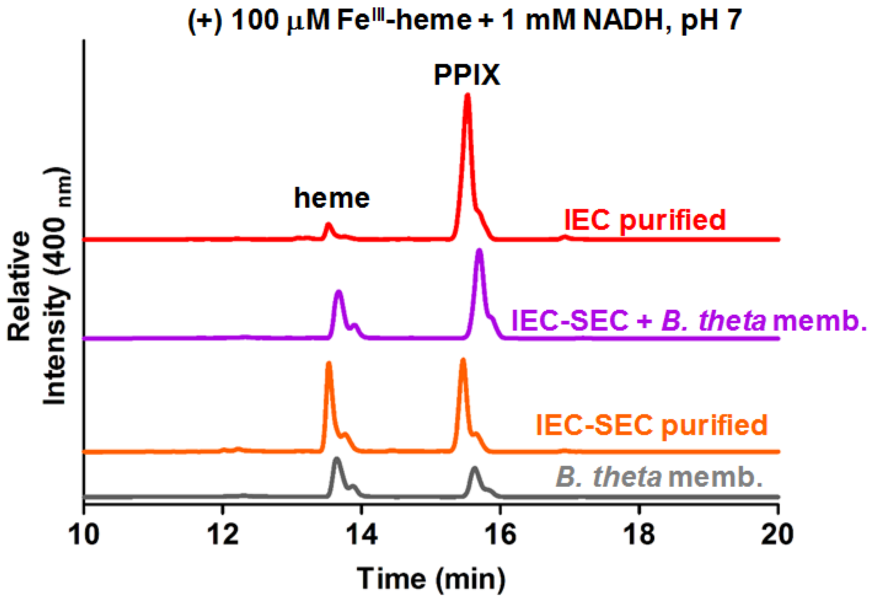

**Figure S12. Heterologously expressed HmuS converts hemin to Fe(II) and PPIX: representative HPLC analyses.** Reaction mixtures contained 100 μM hemin, 1 mM NADH, in 330 μL total reaction volumes (20 mM tris-HCl, 250 mM NaCl, pH 7) and the following protein fractions: partially (IEC) purified HmuS (3.2 mg $mL^{-1}$) (red); IEC and SEC purified HmuS (3.2 mg $mL^{-1}$) (orange); *B. theta* membrane fraction (150 μL from a stock prepared from 0.1 g cells $mL^{-1}$) (gray); the same mixture as in gray but with added purified HmuS (3.2 mg $mL^{-1}$ ) (purple). All the reactions were incubated for 40 min at room temperature prior to extraction and analysis by HPLC. Extractions used 300 μL ACN:12 M HCl:DMSO (41:9:50, v/v/v) with the exception of the IEC+SEC purified HmuS (3.2 mg $mL^{-1}$) reaction mixture which was extracted with 600 μL ACN:12 M HCl:DMSO (41:9:50, v/v/v).

```
HmuS  ------------------------------MKKKSKIILGGCIVVAALIGLSVWNTWFS 29
                                      12345678901234567890123456789

       β I-1      α I-1          β I-2     α I-2       β I-3
HmuS  ATKIAFVNFQTIQQGSISKANDN-SFIKLSEVSLDNLD-RLT---SYDMVFINGMGLRIV 84
      01234567890123456789012 34567890123456 789   012345678901234
       * :. .: .          :         :                *     .**:

       α I-3          β I-4    α I-4      α I-5          α I-6
HmuS  EEQRQQIQQAADKGIPVYTSMATNPANNICNLDSIQQNLIRGYLSNGGKTNYRNMLNYIR 144
      567890123456789012345678901234567890123456789012345678901234
        .   :                  .* * . .         :  *       *  .    :*

                                β II-1        β II-2     α II-1
HmuS  KAIDGKA-SAVPEVEDPIERPSDMLYH-AGISNP---DDEQEFLTVADYEKFMQENNLY- 198
      5678901 234567890123456789 0 123456   789012345678901234567 8
        :* *                     .

                β II-3                α II-2      β II-4     α II-3
HmuS  KE----------GARKIMITGQMADAT--------DLIKALENAGYNVYPVQSMTRFMS- 239
      90          123456789012345          678901234567890123456789
                          : :   :                  :   ::

       α II-3    β II-5            α II-4       β II-     α II-5
HmuS  -FIEEVQPDAVINMAHGRMG----DKMVDYLKARNILLFAPLTINSLVDEWENDPMGMSG 294
       012345678901234567 8    901234567890123456789012345678901234
          :     : : ***:.     :     :    :: ::  :         *   .   : *

       α II-6   α II-7        β II-7       β II-8       α II-8
HmuS  GFMSQSIVTPEIDGAIRPFALFAQYED-KEGLRHSYAVPERLKTFVSTIDNYLNLKTKPN 353
      56789012345678901234567890 1 234567890123456789012345678901 23
      * ::*::  *::**        :          .          *    :    :      :

      α III-1  β III-1                α III-2                 α III-3
HmuS  FEKKVAIYYYKGPGQNA-LTAAGMEVVPSLYNLLLRMKQEGYNISGLPANAQELGKMIQA 412
      45678901234567890 123456789012345678901234567890123456789012
        *:: : : ** *     : *  :..  *:. .*  :.          :       :   .

                   α III-4       β III-2  α III-5        α III-6
HmuS  QGAVFNAYAEGAFNDFMQNGHPELITKEQYESWVKESLRPEKYQEVVDAFGEFPGNYMVT 472
      345678901234567890123456789012345678901234567890123456789012
       *     *  .    :         :        *     .            .  *.  :

                β III-3 β III-4 β III-5                    α III-7
HmuS  P---DGKLGIARLQFGNVVLLPQNAAGSGDNSFQVVHGTDMAPPHTYIASYLWMQHGFKA 529
      3   45678901234567890123456789012345678901234567890123456789
                .:     :*:: ::**    .    : * : **.  .  : : . **:     : .

        β III-6   α III-8       α III-9      β III-7     α III-10   β III-8
HmuS  DALIHFGTHGSLEFTPRKQVALCSNDWPDRLVGAVPHYYLYSIGNVGEGMMAKRRSYATL 589
      012345678901234567890123456789012345678901234567890123456789
      : ..***:**.:*:** :* .:     * : : *  ** * ::   ::**.: ***   *

      β III-8         α IV-1
HmuS  QSYLTPPFLESSVRGIYRELMEKIKIYNNSQKANK-----------------------DQ 626
      0123456789012345678901234567890123 4                       56
       . :*  :  .         *   :  :

         α IV-2       α IV-3            α IV-4
HmuS  ESLAVKTLTVKMGIHRDLGLDSMA-NKPYTEDEIARVENFAEELATEKITGQLYTMGVPY 685
```

```
      78901234567890123456789O 1234567890123456789012345678901234 5
         ::       :*     * :.             :  :  : :*:    :: .  *  *  *
      α IV-5        α IV-6            α IV-7  α IV-8    α IV-9
HmuS  EPERITSSVYAMATEPIAYSLFALDKQRGKATESAEKHRSVFTQQYLMPARLLVERLMA- 744
      67890123456789012345678901234567890123456789012345678901234
             :   :  : :* .    :
     α IV-10 α IV-11    α IV-12
HmuS  NPSLATDELICHTAGITPQELAKARQIEAERNAPKG-MMAM-----MMAAAAKKDQADNE 798
      567890123456789012345678901234567890 1234     5678901234567 8
                          :  :

HmuS  PSG-------------------NGHPASA----KMEKGPHGKMPAGMKEAMK------KMG 830
      901                   2345678    9012345678901234567      890

HmuS  ANMDPEK--AMEMAK-SM---------------------GA-SPEAL----KKMEASMKA 861
      1234567  890123 45                     67 89012    345678901

HmuS  NKDTST------------------------------------------------------ 867
      234567

HmuS  ----------------------DASG--KPAMA---GKTEKPQGMSAMMAAMG--KAPKEY 899
                            8901  23456   78901234567890123  456789

         α IV-12              α IV-12, α III-11
HmuS  SKEEVEFALAVAEVERTIKNVGNYKNALLTSPEEELSSLMNALKGGYTAPTPGGDPIANP 959
      012345678901234567890123456789012345678901234567890123456789
                :    : : .        :       *.       :    :   *: .** : .
   α IV-12      α V-1        α V-2                    β V-1    α V-3
HmuS  NTLPTGRNMYAINAEATPTESAWEKGIALAKQTIDRYKQRHNDSIPRKVSYTLWSSEFIE 1019
      012345678901234567890123456789012345678901234567890123456789
        : :* *::.:* *   *:  ** :        ::        : .   * *: :*:**.**::
        α V-4     β V-2     β V-3   α V-5        β V-4  α V-6   α V-7
HmuS  TGGATIAQVLYMLGVEPVRDAFGRVSDLKLIPSTELGRPRIDVVVQTSGQLRDLAASRLF 1079
      012345678901234567890123456789012345678901234567890123456789
      * *:::.: : :**:.*: * :  : *: *      * * *:*  *****:**: :* *
      α V-7                α V-8              α V-9
HmuS  LINRAVEMAAAAKDD-KYENQVASSVIEAERVLTEKGLSPKDAREISTFRVFGGANGMYG 1138
      012345678901234 56789012345678901234567890123456789012345678
      :: :*: **:             * *   .     :. *   *: *  *    :  *:**. .. **
      α V-10 α V-11  α V-12                  α V-13    β V-5
HmuS  TGIQEMVESGDRWENESEIADTYLNNMGAYYGSEKNWEVFQKFAFEAALTRTDVVVQPRQ 1198
      901234567890123456789012345678901234567890123456789012345678
      :.*   :    ..:*    ::    *: **.   *        *         :   .*  :* ::****
           α V-14          β V-6     β V-7  α VI-1
HmuS  SNTWGALSLDHVYEFMGGMNLTVRNVTGKDPDAYLSDYRNRNHMKMQELKEAVGVESRTT 1258
```

```
      901234567890123456789012345678901234567890123456789012345678
      .* *..*****::*****::::    **::*:**::* *    . .:* ::**:.:  * :

         α VI-2          α VI-3         αVI-4       α VI-5
HmuS  ILNPTYIKEKMKGGASSASEFAEVITNTYGWNVMKPAAIDKELWDNIYNVYVKDELNLGV 1318
      901234567890123456789012345678901234567890123456789012345678
      ::*  ::   : .       .   : *::.*:* .   :    *   :  *  *   :

       α VI-6       α VI-7                   α VI-8
HmuS  KQYFEQQNPAALEEMTAVMLESARKGLWQASEEQVAELSKLHTEIVNTYRPSCSGFVCDN 1378
      901234567890123456789012345678901234567890123456789012345678
             .  :   ::: * :   ** *             .

         α VI-8           α VI-8
HmuS  AKLRDFIASKAD-A--QTATQYKENISKIREAKASGSN--KGVVMKKEEMN-Q-TAENQT 1431
      901234567890 1  23456789012345678901234  5678901234 5 678901
        :

HmuS  NTL-SNVA-VGIA--VIIVILALILFVRKRRKSSQM----------- 1463
      234 5678 9012  345678901234567890123
```

**Figure S13. Strictly conserved residues from alignments of HmuS orthologues in complete *hmuYRSTUV* operons.** Conserved residues are indicated by an *. Secondary structure elements from the cryo-EM single particle structure are indicated above the *B theta* HmuS sequence. Domain boundaries are: Domain I 30-162; Domain II 163-350; Domain III 351-594; Domain IV 595-779 and 899-930; MRI 780-898; Domain V 972-1245; Domain VI 1246-1405.

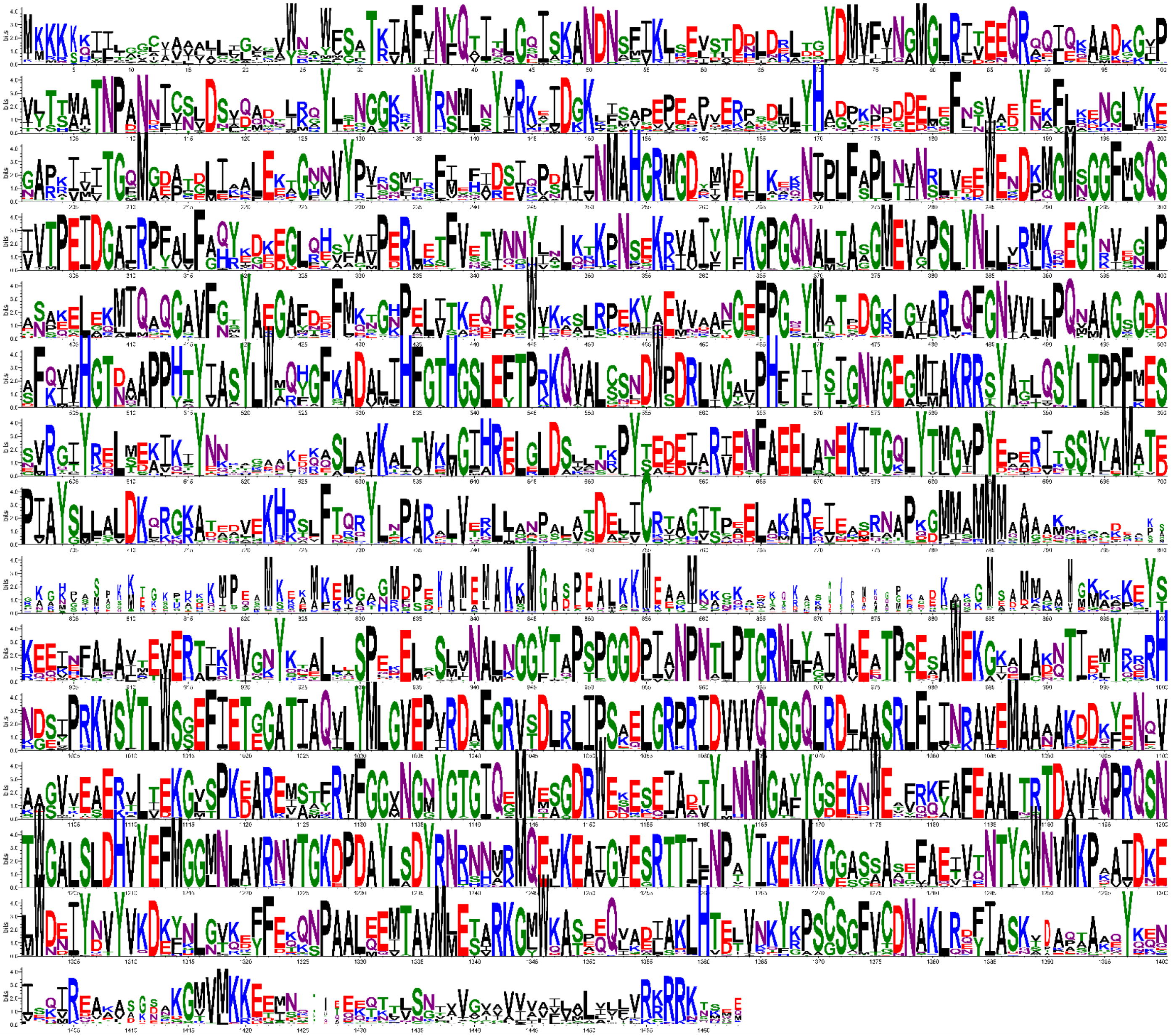

**Figure S14. Sequence logo representation illustrating conservation among operonic HmuS sequences.** As we included only operonic HmuS sequences with at least 50% identity and 95% coverage to *B theta* HmuS, this logo is most representative of organisms related to *B theta*. See also Supplementary Data File 04.

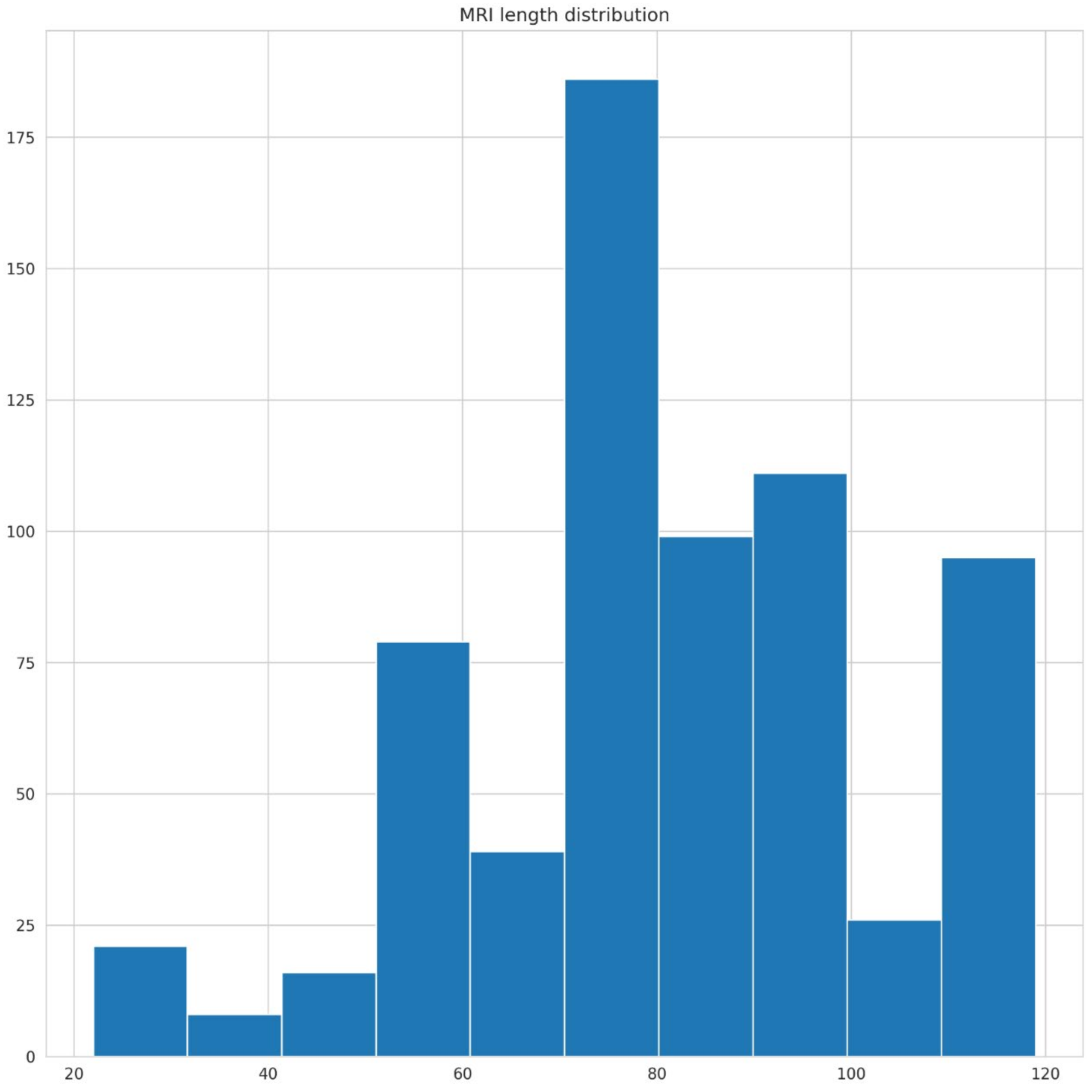

**Figure S15. Length distribution of MRIs from the operonic set of HmuS sequences.** A subset of HmuS sequences corresponding to *B. theta* HmuS residues 780-898 was excised from the multiple sequence alignment. Fragment lengths were determined for all sequences and their distribution is shown as a histogram.

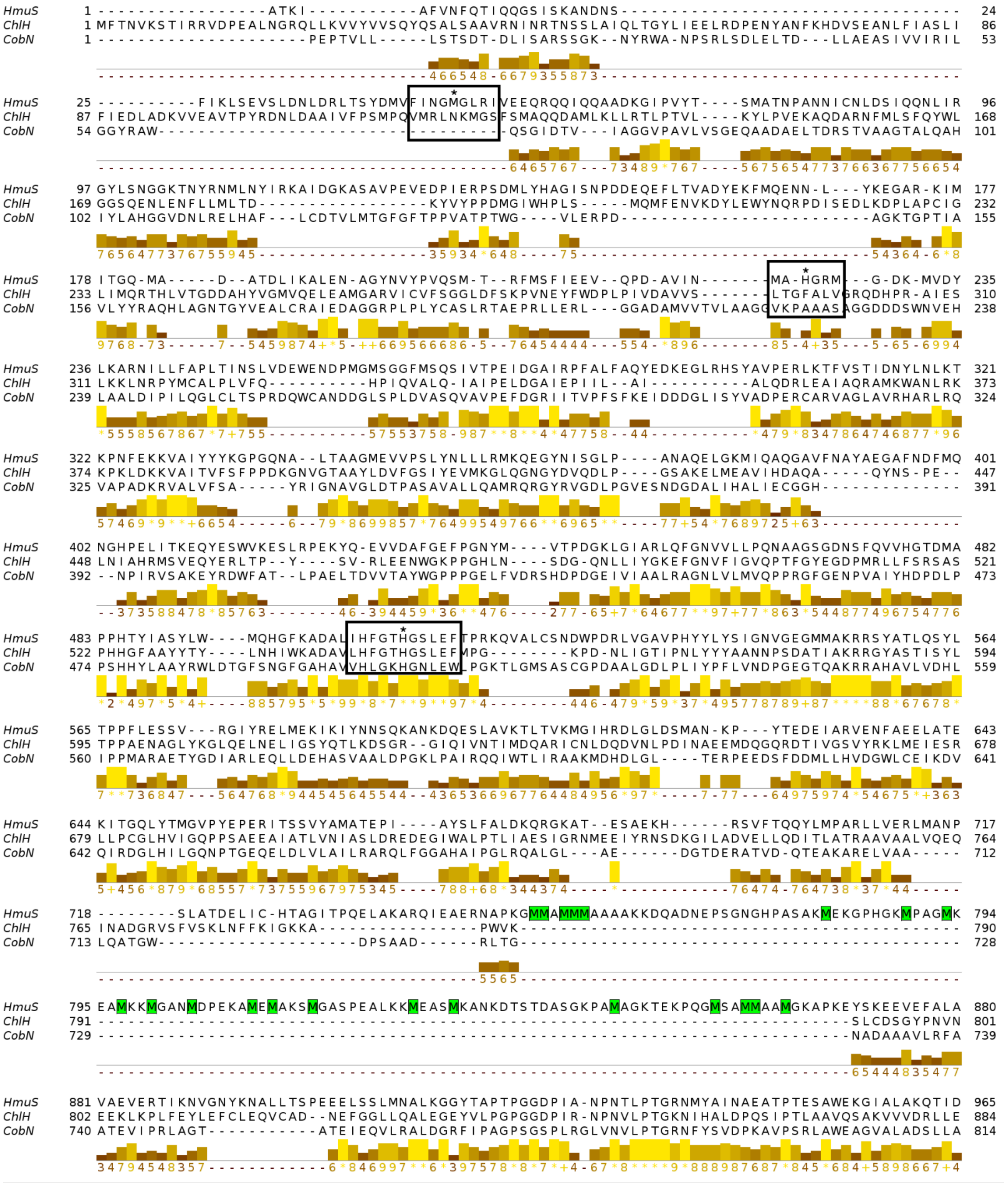

HmuS 1 -----------------ATKI-----------AFVNFQTIQQGSISKANDNS----------------------------------- 24
ChlH 1 MFTNVKSTIRRVDPEALNGRQLLKVVYVVSQYQSALSAAVRNINRTNSSLAIQLTGYLIEELRDPENYANFKHDVSEANLFIASLI 86
CobN 1 ---------------------PEPTVLL-----LSTSDT-DLISARSSGK--NYRWA-NPSRLSDLELTD---LLAEASIVVIRIL 53
HmuS 25 ----------FIKLSEVSLDNLDRLTSYDMVFINGMGLRIVEEQRQQIQQAADKGIPVYT----SMATNPANNICNLDSIQQNLIR 96
ChlH 87 FIEDLADKVVEAVTPYRDNLDAAIVFPSMPQVMRLNKMGSFSMAQQDAMLKLLRTLPTVL----KYLPVEKAQDARNFMLSFQYWL 168
CobN 54 GGYRAW-----------------------------------QSGIDTV---IAGGVPAVLVSGEQAADAELTDRSTVAAGTALQAH 101
HmuS 97 GYLSNGGKTNYRNMLNYIRKAIDGKASAVPEVEDPIERPSDMLYHAGISNPDDEQEFLTVADYEKFMQENN-L---YKEGAR-KIM 177
ChlH 169 GGSQENLENFLLMLTD-----------------KYVYPPDMGIWHPLS-----MQMFENVKDYLEWYNQRPDISEDLKDPLAPCIG 232
CobN 102 IYLAHGGVDNLRELHAF---LCDTVLMTGFGFTPPVATPTWG----VLERPD-------------------------AGKTGPTIA 155
HmuS 178 ITGQ-MA-----D--ATDLIKALEN-AGYNVYPVQSM-T--RFMSFIEEV--QPD-AVIN-------MA-HGRM---G-DK-MVDY 235
ChlH 233 LIMQRTHLVTGDDAHYVGMVQELEAMGARVICVFSGGLDFSKPVNEYFWDPLPIVDAVVS-------LTGFALVGRQDHPR-AIES 310
CobN 156 VLYYRAQHLAGNTGYVEALCRAIEDAGGRPLPLYCASLRTAEPRLLERL---GGADAMVVTVLAAGGVKPAAASAGGDDDSWNVEH 238
HmuS 236 LKARNILLFAPLTINSLVDEWENDPMGMSGGFMSQSIVTPEIDGAIRPFALFAQYEDKEGLRHSYAVPERLKTFVSTIDNYLNLKT 321
ChlH 311 LKKLNRPYMCALPLVFQ----------HPIQVALQ-IAIPELDGAIEPIIL--AI-----------ALQDRLEAIAQRAMKWANLRK 373
CobN 239 LAALDIPILQGLCLTSPRDQWCANDDGLSPLDVASQVAVPEFDGRIITVPFSFKEIDDDGLISYVADPERCARVAGLAVRHARLRQ 324
HmuS 322 KPNFEKKVAIYYYKGPGQNA--LTAAGMEVVPSLYNLLLRMKQEGYNISGLP----ANAQELGKMIQAQGAVFNAYAEGAFNDFMQ 401
ChlH 374 KPKLDKKVAITVFSFPPDKGNVGTAAYLDVFGSIYEVMKGLQGNGYDVQDLP----GSAKELMEAVIHDAQA-----QYNS-PE-- 447
CobN 325 VAPADKRVALVFSA-----YRIGNAVGLDTPASAVALLQAMRQRGYRVGDLPGVESNDGDALIHALIECGGH-------------- 391
HmuS 402 NGHPELITKEQYESWVKESLRPEKYQ-EVVDAFGEFPGNYM----VTPDGKLGIARLQFGNVVLLPQNAAGSGDNSFQVVHGTDMA 482
ChlH 448 LNIAHRMSVEQYERLTP--Y----SV-RLEENWGKPPGHLN----SDG-QNLLIYGKEFGNVFIGVQPTFGYEGDPMRLLFSRSAS 521
CobN 392 --NPIRVSAKEYRDWFAT--LPAELTDVVTAYWGPPPGELFVDRSHDPDGEIVIAALRAGNLVLMVQPPRGFGENPVAIYHDPDLP 473
HmuS 483 PPHTYIASYLW---MQHGFKADALIHFGTHGSLEFTPRKQVALCSNDWPDRLVGAVPHYYLYSIGNVGEGMMAKRRSYATLQSYL 564
ChlH 522 PHHGFAAYYTY----LNHIWKADAVLHFGTHGSLEFMPG-------KPD-NLIGTIPNLYYYAANNPSDATIAKRRGYASTISYL 594
CobN 474 PSHHYLAAYRWLDTGFSNGFGAHAVVHLGKHGNLEWLPGKTLGMSASCGPDAALGDLPLIYPFLVNDPGEGTQAKRRAHAVLVDHL 559
HmuS 565 TPPFLESSV---RGIYRELMEKIKIYNNSQKANKDQESLAVKTLTVKMGIHRDLGLDSMAN-KP---YTEDEIARVENFAEELATE 643
ChlH 595 TPPAENAGLYKGLQELNELIGSYQTLKDSGR--GIQIVNTIMDQARICNLDQDVNLPDINAEEMDQGQRDTIVGSVYRKLMEIESR 678
CobN 560 IPPMARAETYGDIARLEQLLDEHASVAALDPGKLPAIRQQIWTLIRAAKMDHDLGL----TERPEEDSFDDMLLHVDGWLCEIKDV 641
HmuS 644 KITGQLYTMGVPYEPERITSSVYAMATEPI----AYSLFALDKQRGKAT--ESAEKH------RSVFTQQYLMPARLLVERLMANP 717
ChlH 679 LLPCGLHVIGQPPSAEEAIATLVNIASLDREDEGIWALPTLIAESIGRNMEEIYRNSDKGILADVELLQDITLATRAAVAALVQEQ 764
CobN 642 QIRDGLHILGQNPTGEQELDLVLAILRARQLFGGAHAIPGLRQALGL---AE------DGTDERATVD-QTEAKARELVAA----- 712
HmuS 718 --------SLATDELIC-HTAGITPQELAKARQIEAERNAPKGMMAMMMAAAAKKDQADNEPSGNGHPASAKMEKGPHGKMPAGMK 794
ChlH 765 INADGRVSFVSKLNFFKIGKKA----------------PWVK-------------------------------------------- 790
CobN 713 LQATGW-------------------DPSAAD------RLTG--------------------------------------------- 728
HmuS 795 EAMKKMGANMDPEKAMEMAKSMGASPEALKKMEASMKANKDTSTDASGKPAMAGKTEKPQGMSAMMAAMGKAPKEYSKEEVEFALA 880
ChlH 791 -------------------------------------------------------------------------SLCDSGYPNVN 801
CobN 729 -------------------------------------------------------------------------NADAAAVLRFA 739
HmuS 881 VAEVERTIKNVGNYKNALLTSPEEELSSLMNALKGGYTAPTPGGDPIA-NPNTLPTGRNMYAINAEATPTESAWEKGIALAKQTID 965
ChlH 802 EEKLKPLFEYLEFCLEQVCAD--NEFGGLLQALEGEYVLPGPGGDPIR-NPNVLPTGKNIHALDPQSIPTLAAVQSAKVVVDRLLE 884
CobN 740 ATEVIPRLAGT-----------ATEIEQVLRALDGRFIPAGPSGSPLRGLVNVLPTGRNFYSVDPKAVPSRLAWEAGVALADSLLA 814

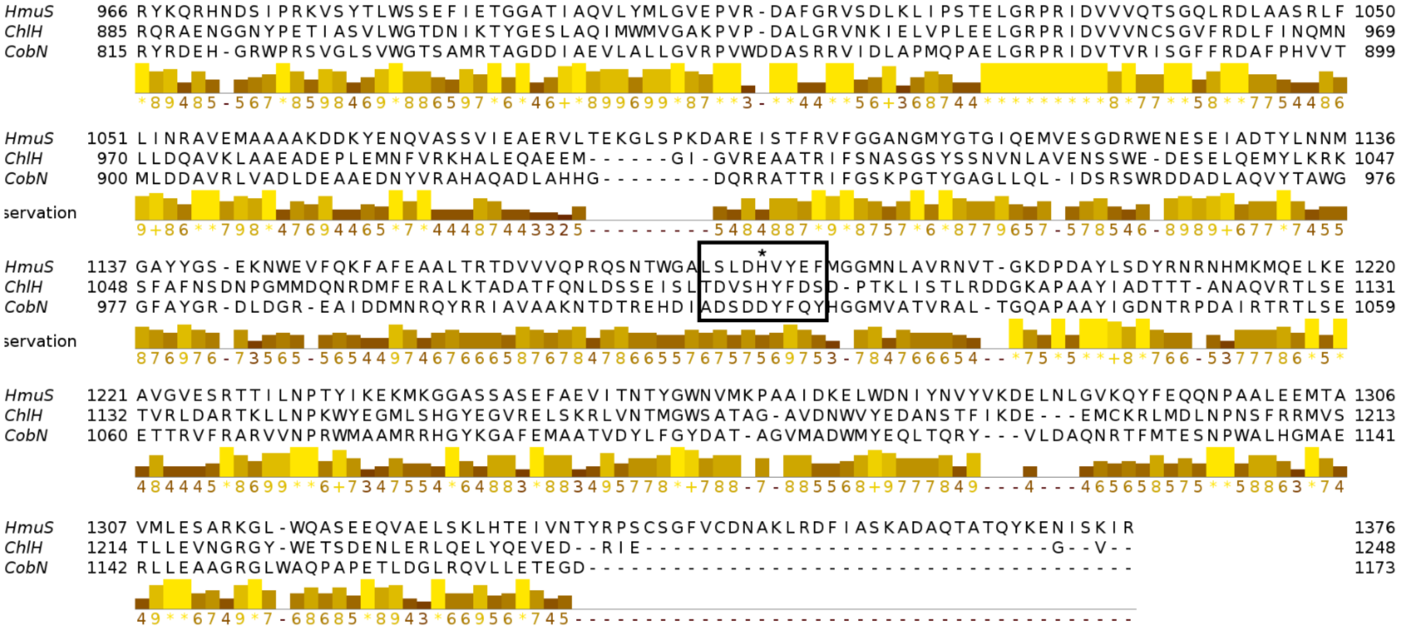

**Figure S16. Structure-based sequence alignment of CobN (6yt0), HmuS (this work) and CobN (7c6o).** Structures were aligned using default parameters in FoldMason (https://www.biorxiv.org/content/10.1101/2024.08.01.606130v2). High numbers and gold/yellow conservation block below the alignment indicate residues that are well-conserved in all members. Dark colors or the lack of colored blocks indicate low levels of conservation. Over-represented methionines in the MRI (missing in the structure) are highlighted in green. Black rectangles of arbitrary width are drawn to include the neighborhood around four residues strictly conserved in HmuS proteins: Met-79, His-254, His-538, and His-1209. The residues are marked by black asterisks. Met-79 and His-254 coordinate the heme molecule at the head/neck interface. These residues are not strictly conserved in other type 1 chelatase superfamily members. His-538 (third rectangle) is involved in coordinating the heme molecule in the putative reaction center and is strictly conserved in all chelatase superfamily members. His-1209 (fourth rectangle) is also predicted to coordinate the heme molecule in the putative reaction center, but it is not universally conserved.

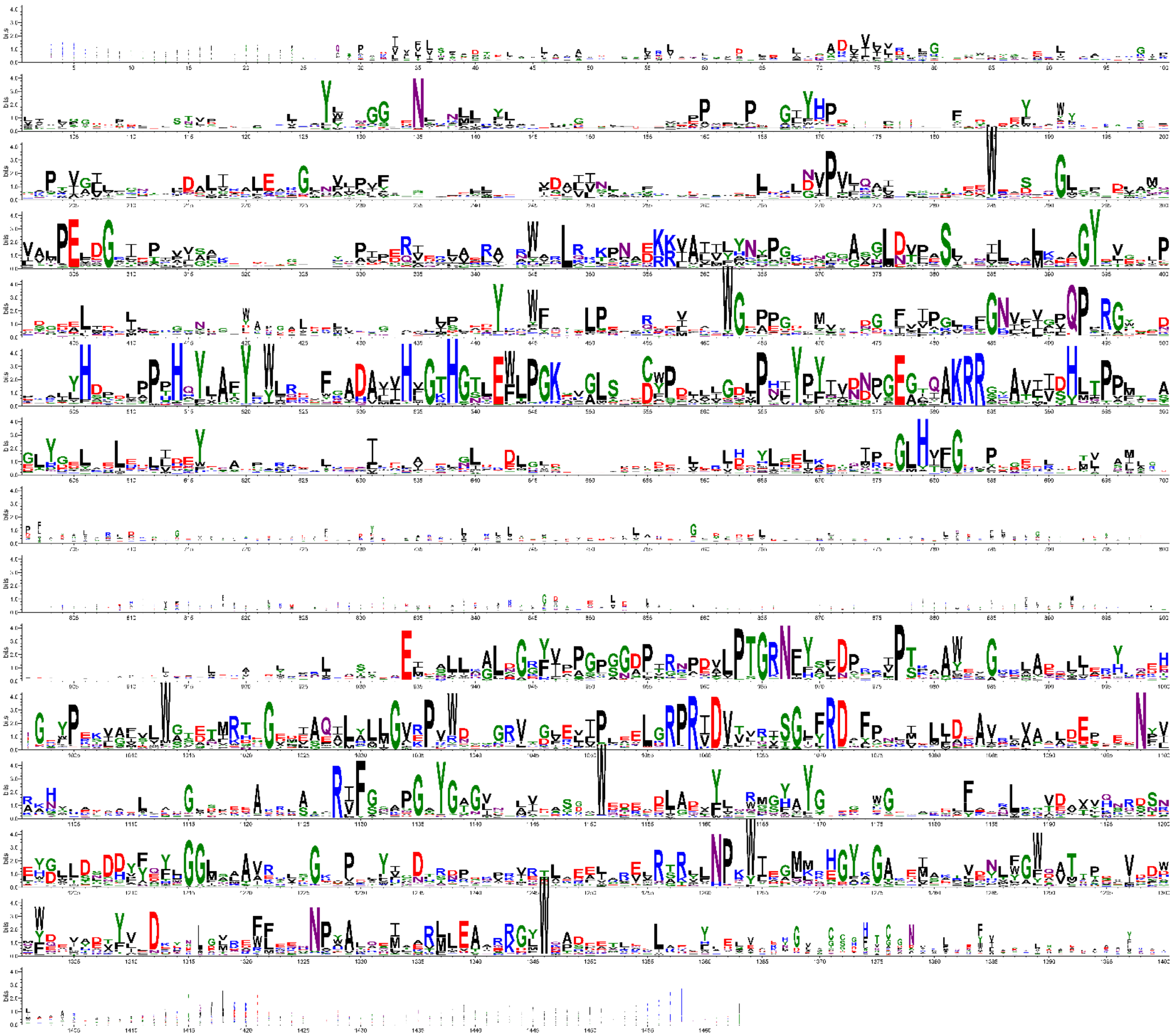

**Figure S17. Sequence logo representation illustrating conservation among type 1 chelatase sequences.** As the MRI is not found in chelatase sequences other than HmuS, the middle part of the sequence logo has almost no conservation.

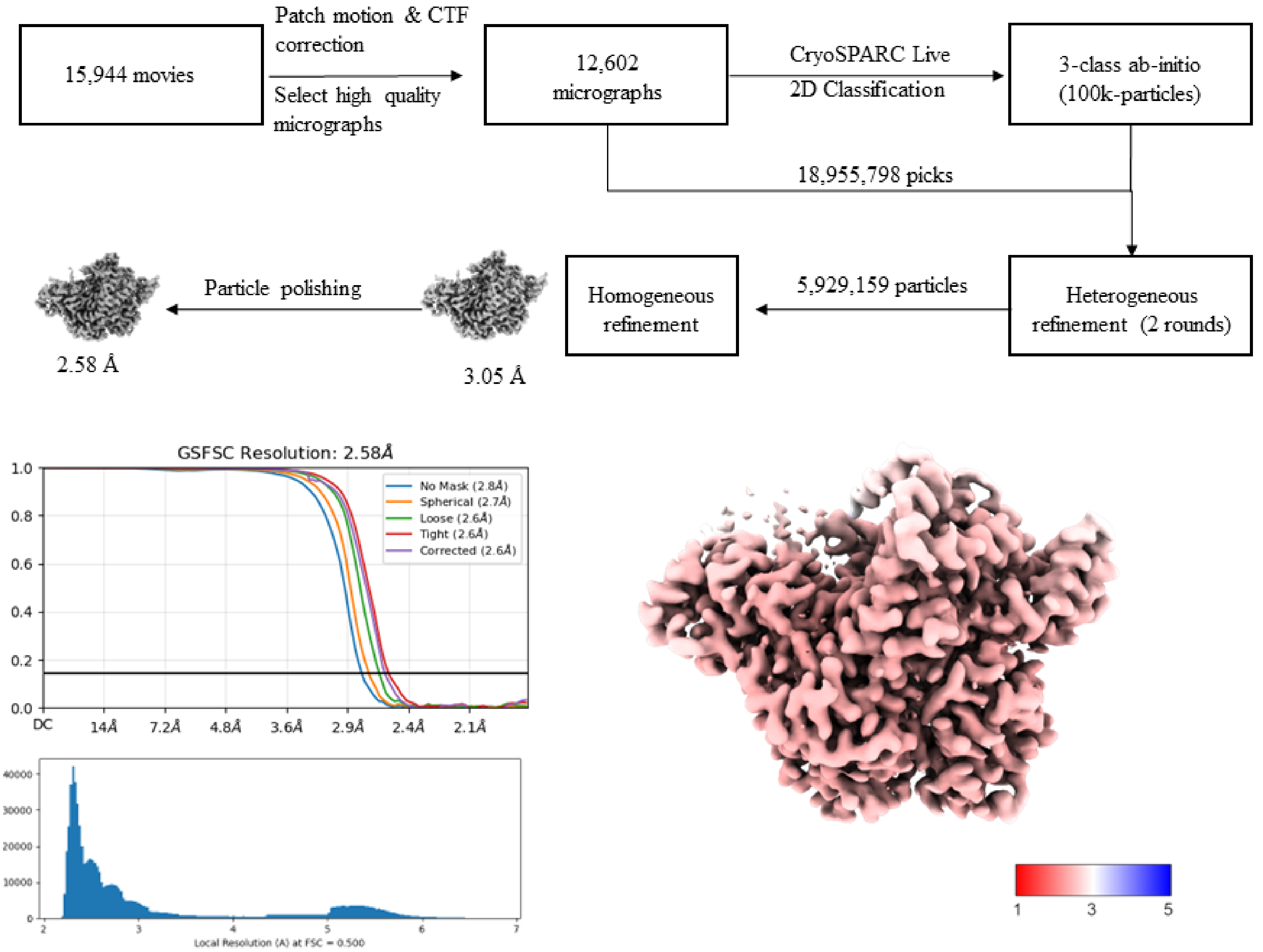

**Figure S18. Single particle reconstruction workflow for HmuS. See Methods for details.**

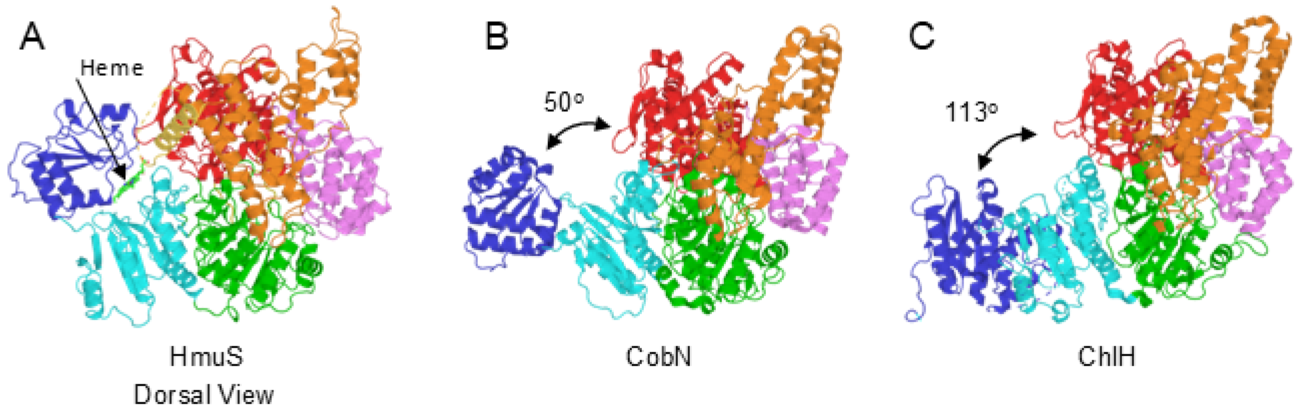

**Figure S19. Head domain conformations.** A) HmuS is colored as in Figure 1, with the head domain in blue, the neck in cyan, and domains III-VI in green, orange, red and purple. The orientation is a “dorsal” view, i.e., looking down on the domain IV “backbone”, with the membrane roughly in the plane of the paper underneath HmuS, and heme bound at the head/neck interface. The first helix of the methionine rich insertion (MRI) is shown in yellow, with the dashed line indicating a rough position for the 120 disordered residues of the MRI, potentially in position to interact with the heme binding site at the neck/head interface. B) CobN shown in an equivalent orientation. Domains II-VI superpose well on HmuS, but the head domain is rotated away from the neck domain by 50°. C) ChlH in the same relative orientation, showing even greater movement of the head domain relative to the neck and body domains. Superposition of the HmuS head domain on the ChlH head domain requires a 113° rotation. The differing orientations of the head domain in CobN and ChlH suggest the HmuS head domain may also sample open conformations in the absence of heme.

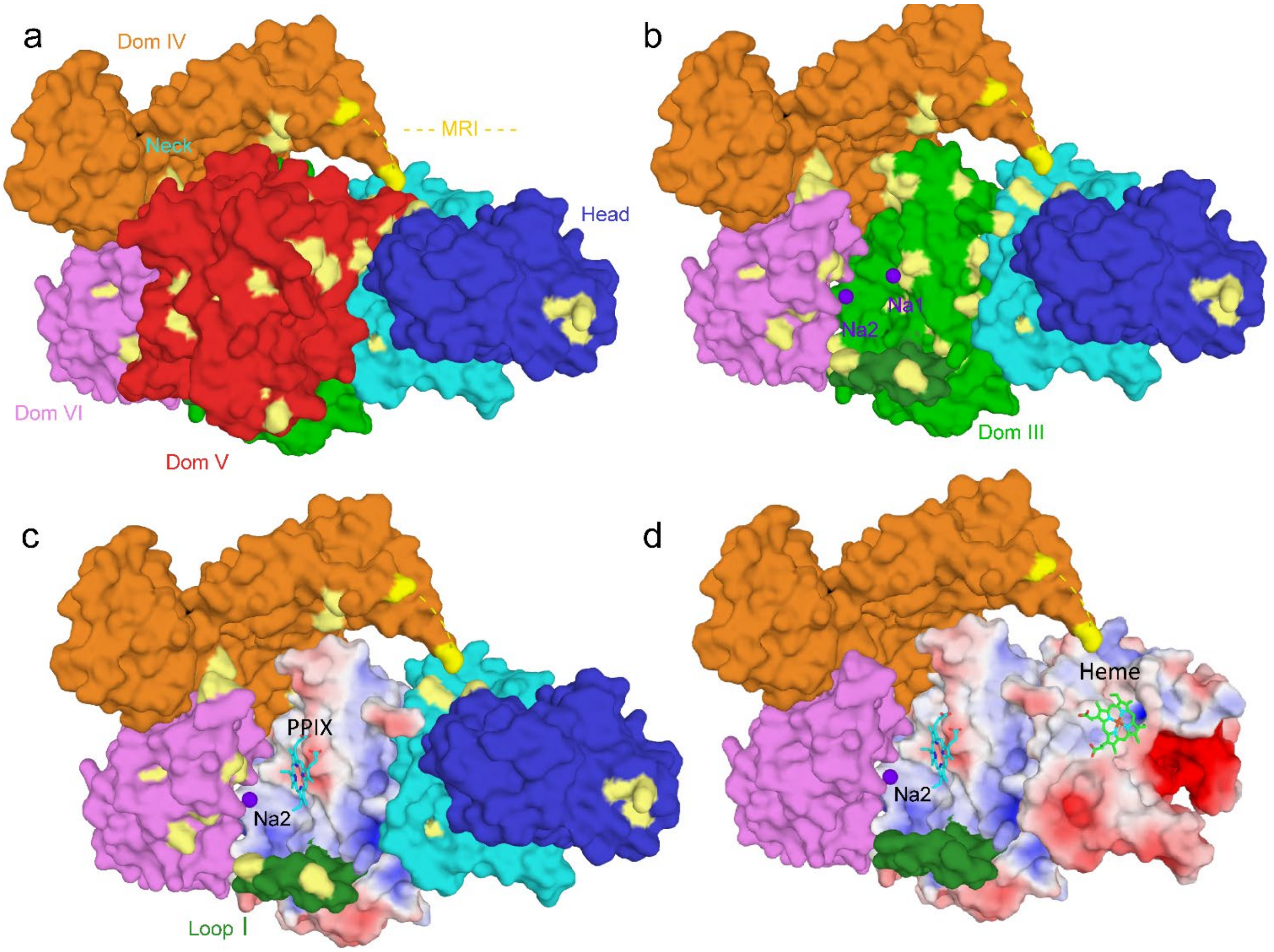

**Figure S20. Ligand binding sites, backside view.** (a) HmuS is shown from the "backside", a 180$^o$ rotation about the vertical axis relative to Figure 9. Domains are colored as in Figures 8 and 9, with strictly conserved residues in pale yellow. The absence of the MRI is indicated by the yellow dashed line and yellow anchor points on domain IV (orange). (b) Domain V has been removed, revealing the central cleft with 2 bound Na ions (purple), Na1 and Na2, and Loop I at the bottom of the structure. Na1 is coordinated by strictly conserved His-1209, which lies in the center of the exposed domain III face. The domain III face of the central cleft is less conserved than the domain V face. (c) The docked protoporphyrin IX (PPIX) and Na2 are shown, along with the electrostatic surface domain III (+/- 5kT/e). (d) Domain I (blue) is also removed, revealing the heme binding site at the Head/Neck interface. The heme and PPIX molecules are separated by nearly 30 Å. Electrostatic surfaces are also shown for domain II.

**Supplementary Tables**

**Table S1. Substrate and product analyses for heme-PPIX conversion reactions† using *B theta* cellular fractions and/or heterologously expressed HmuS. These data are plotted as bar charts in Figure 3.**

| Sample # | Fraction assayed | Fig | Preparation of protein fraction | Amount of fraction assayed | Assay additions | Unreacted [heme] post-incubation (mM) | [PPIX] product post-reaction (mM) | % turn-over: [PPIX] ([heme] + [PPIX])$^{-1}$ x 100% |
|---|---|---|---|---|---|---|---|---|
| 1 | *B theta* whole cell lysate | 3a | 1 g cell pellet resuspended in 15 mL of 20 mM Tris-HCl (pH 7.1). | 300 µL | None | 30 ± 1.6 | 5.2 ± 1 | 15 |
| 2 | *B theta* whole cell lysate, dialyzed | 3a | Sample 1, dialyzed 10 kDa MWCO tubing | 300 µL | None | 32 ± 0.7 | 4.6 ± 0.4 | 13 |
| 3 | *B theta* whole cell lysate, dialyzed + NADH | 3a | Sample 2 | 300 µL | 1 mM NADH | 12 ± 0.3 | 29 ± 0.9 | 71 |
| 4 | *B theta* whole cell lysate, dialyzed + ATP | 3a | Sample 2 | 300 µL | 1 mM ATP | 30 ± 0.8 | 0 | 0 |
| 5 | *B theta* whole cell lysate, dialyzed + NADH + ATP | 3a | Sample 2 | 300 µL | 1 mM NADH, 1 mM ATP | 13 ± 0.7 | 26 ± 0.3 | 67 |
| 6 | *B theta* membranes | 3b | Sample 2 (4 mL) was pelleted, washed, and resuspended in 4 mL reaction buffer | 300 µL | 1 mM NADH | 8.3 ± 0.5 | 36 ± 0.3 | 81 |
| 7 | *B theta* soluble lysate | 3b | Supernatant from pelleted sample 6 | 300 µL | 1 mM NADH | 23 ± 2 | 13 ± 2 | 36 |
| 8 | Heterologously expressed HmuS, IEC purified | 3c | 3.2 mg mL$^{-1}$ protein stock, containing 2.3 mM HmuS-heme | 300 µL | 1 mM NADH | 7.1 ± 0.3 | 82 ± 1 | 92 |
| 9 | Heterologously expressed HmuS, IEC-SEC purified | 3c | 3.2 mg mL$^{-1}$ protein stock; 2 mM HmuS-heme | 300 µL | 1 mM NADH | 32 ± 2 | 37 ± 0.2 | 54 |
| 10 | *B theta* membrane fractions | 3c | Sample 6 | 150 µL | 1 mM NADH | 21 ± 0.5 | 16 ± 0.5 | 44 |
| 11 | Purified HmuS and *B theta* membrane fractions: mixed | 3c | 150 µL sample 6, 150 µL sample 9 | 300 µL total | 1 mM NADH | 22 ± 2 | 46 ± 0.5 | 68 |

†Standard reaction conditions: 100 µM hemin, 40 min room temperature, 330 µL final reaction volume, 20 mM Tris-HCl buffer, pH 7. Extraction conditions: 300 µL or 600 µL ACN + 12M HCl + DMSO (41:9:50). Extraction efficiencies (Fig S1) are near 50% for both heme and PPIX in this solvent. Analyte concentrations in the extracts were measured via HPLC peak integration and comparison with standard curves, and corrected to reflect the concentration in the initial reaction volume. ≥3 reactions were analyzed for substrate-product turnover and analyte concentrations averaged.

**Table S2. Typical outcomes for recombinant HmuS expression and purification†**

| Fraction | Volume (mL) | Protein concentration (mg $mL^{-1}$) | Total protein (mg) | Yield after step (% by mass) | Total yield (% by mass) |
|---|---|---|---|---|---|
| HmuS-containing clarified lysate | 100 | 23 | 2300 | 100 | 100 |
| Anion exchange | 40 | 15 | 600 | 26 | 26 |
| Size exclusion chromatography | 20 | 5 | 100 | 16 | 4.4 |
| Centrifuge concentration (50 MWCO) | 9 | 10 | 90 | 15 | 3.9 |

† For HmuS purification, 14 g cell pellets were generated per L of culture, frozen, and resuspended in 100 mL of 20 mM Tris-HCl, 250 mM NaCl, pH 7.1 buffer, lysed by sonication, and ultracentrifuged as described in the text. Protein concentrations were measured by Bradford analysis (BSA, 0.1 mg $ml^{-1}$, Bradford dye as the working dye, standard curve was constructed with triplicate data points at 595 nm).

**Table S3. Cryo-EM data collection, processing, model refinement and validation.**

| | **EMD-46483 PDB 9D26** |
|---|---|
| **Data collection** | |
| Microscope | Talos Arctica |
| Voltage (kV) | 200 |
| Detector | K3 (Counting) |
| Magnification (nominal/calibrated) | ×45,000/×55,187 |
| Exposure navigation | Image shift to 25 holes |
| Electron exposure ($e^{-}$/Å$^{2}$) | 56 $e^{-}$ Å$^{2}$ |
| Data acquisition software | SmartScope/SerialEM |
| Total electron exposure ($e^{-}$/Å$^{2}$) | 56 |
| Exposure rate (e-/pixel/sec) | 18.3 |
| Frame length (ms) | 60 |
| Number of frames per micrograph | 51 |
| Pixel size (Å) | 0.9061 |
| Defocus range (µm) | -0.6 to -1.5 |
| Micrographs collected ($0^{o}$ tilt) | 7,800 |
| Micrographs collected ($15^{o}$ tilt) | 8,194 |
| **Reconstruction** | |
| Image processing package | cryoSPARC |
| Total extracted particles (blob picks) | 18,955,798 |
| Final number particles | 5,929,8976 |
| Symmetry imposed | C1 |
| Resolution (Å) | |
| FSC 0.143 (masked/unmasked) | 2.8/2.6 |
| **Model Composition (#)** | |
| Chains | 1 |
| Atoms | 9993 |
| Residues | Protein: 1257 |
| Water | 66 |
| Ligands | HEM: 1<br>NA: 2 |
| **Model Refinement** | |
| Refinement Package | Phenix |
| Bonds (RMSD) | |
| Length (Å) (# > 4σ) | 0.005 (0) |
| Angles (°) (# > 4σ) | 0.945 (1) |
| **Model Validation** | |
| MolProbity score | 0.97 |
| Clash score | 1.42 |
| Ramachandran (%) | |
| Outliers | 0 |
| Allowed | 2.47 |
| Favored | 97.53 |
| Rama-Z | |
| whole (N = 1254) | 0.55 (0.24) |
| helix (N = 650) | 1.46 (0.22) |
| sheet (N = 139) | 1.29 (0.44) |
| loop (N = 576) | -1.55 (0.25) |
| Outliers | |
| Rotamer outliers (%) | 0.00 |
| Cβ outliers (%) | 0.00 |
| Peptide Plane, Cis Pro/general (%) | 5.5/0.1 |

| | |
|---|---|
| Peptide Plane, Twisted Pro/general (%) | 0.0/0.0 |
| CaBLAM outliers (%) | 1.60 |
| ADP (B-factors) | |
| Iso/Aniso (#) | 9993/0 |
| min/max/mean | |
| Protein | 80.14/302.4/140.83 |
| Ligand | 113.3/212.9/189.2 |
| Water | 87.0/146.2/117.3 |
| Occupancy | |
| Mean | 1.00 |
| occ = 1 (%) | 99.88 |
| 0 < occ < 1 (%) | 0.12 |
| occ > 1 (%) | 0 |
| **Model vs. Data** | |
| Resolution (Å) | Masked Unmasked |
| d FSC model (0/0.143/0.5) | 2.6/2.6/2.7 2.6/2.7/2.8 |
| CC (mask) | 0.88 |
| CC (box) | 0.78 |
| CC (peaks) | 0.8 |
| CC (volume) | 0.86 |
| Mean CC for ligands | 0.67 |

**Supplementary Data File Descriptions**

**Supplementary File 01.** Complete proteomics data measured from gel bands, confirming the composition of heterologously expressed HmuS and its complete peptide coverage, is given. The major contaminant band is also shown to be associated primarily with HmuS peptides. Identities and relative quantities of detected proteins are reported.

**Supplementary File 02**. Bottom-up proteomics data measured from partially pure (IEC) and more completely purified (IEC and SEC) HmuS is reported. Identities and relative quantities of detected proteins are tabulated. The data suggest an *E. coli* protein that could serve as a promiscuous reductase in the HmuS reaction.

**Supplementary File 03**. Conservation scores on a 0-1 scale were calculated from alignments for each HmuS residue. A HmuS conservation score column was derived from an alignment of 680 operonic sequences (Fig S14). A chelatase conservation column score was derived from an alignment of 1513 ChlH, CobN and HmuS sequences (Fig S16).
